# CLIJ: GPU-accelerated image processing for everyone

**DOI:** 10.1101/660704

**Authors:** Robert Haase, Loic A. Royer, Peter Steinbach, Deborah Schmidt, Alexandr Dibrov, Uwe Schmidt, Martin Weigert, Nicola Maghelli, Pavel Tomancak, Florian Jug, Eugene W. Myers

**Affiliations:** Max Planck Institute for Molecular Cell Biology and Genetics, Pfotenhauerstr. 108, 01307 Dresden, Germany; Center for Systems Biology Dresden, Pfotenhauerstr. 108, 01307 Dresden, Germany; Chan Zuckerberg Biohub, 499 Illinois St, San Francisco, CA 94158, USA; Now with: Helmholtz-Zentrum Dresden-Rossendorf, Bautzner Landstraße 400, 01328 Dresden, Germany

**Author notes:** Correspondence &. both authors contributed equally (Robert Haase, Loic A. Royer).

## Abstract

Graphics processing units (GPU) allow image processing at unprecedented speed. We present CLIJ, a Fiji plugin enabling end-users with entry level experience in programming to benefit from GPU-accelerated image processing. Freely programmable workflows can speed up image processing in Fiji by factor 10 and more using high-end GPU hardware and on affordable mobile computers with built-in GPUs.

---

Modern microscopy generates staggering amounts of multidimensional image data that place increasing demands on processing flexibility and efficiency. One way to speed up image processing is to exploit the parallel processing capabilities of graphics processing units (GPU).

Recently, GPU-acceleration was used in specific image processing tasks such as reconstruction^1, 2^, image quality determination^3^, image restoration^4^, segmentation^5^ and visualisation^6^. However, in these tools, GPU code is fulfilling one specific purpose and is not intended to be reused in other contexts. By contrast, most common image processing tasks are solved by building flexible workflows consisting of simple operations in widely used tools such as ImageJ^7^ and Fiji^8^. Most of these operations were however programmed at a time when GPUs were not commonly used for general purpose processing. Therefore, typical workflows consisting of core ImageJ operations do not take advantage of GPUs. To address this issue we developed a flexible and reusable platform for GPU-acceleration in Fiji.

Our platform, named CLIJ, complements core *I*mage*J* operations with reprogrammed counterparts that take advantage of the Open*CL*^9^ framework to execute on GPUs. Within CLIJ, we implemented a wide range of fundamental image processing functions for morphological filters, spatial transforms, image warping, local and global thresholding, minima/maxima detection, logical operations on binary images, 3D-to-2D projections, and methods of descriptive statistics for quantitative measurements (Suppl. Listing 1).

We then asked how much faster GPU-accelerated versions of individual operations run compared to their counterparts on the central processing unit (CPU). GPUs can do certain operations faster because they have many more processing cores then regular CPUs (Figure 1a). In addition, memory access can be multiple times faster on GPUs depending on the GPU hardware. On the other hand, in order to be processed on GPUs, the data and the compiled program have to be first pushed to GPU memory, and later data have to be pulled back to CPU memory. While this introduces an unavoidable overhead to any GPU operation, once the data are on the GPU, functions we implemented typically run faster on GPU compared to CPU. Furthermore, optimal performance can be achieved by chaining GPU operations and re-using memory to spare memory transfer and allocation time. The speed-up also depends on the image size and for functions that have parameters, the achievable speed may also depend on the values of the parameters (Figure 1b, Suppl. Figures 1, 2 and 3). Furthermore, after the first execution of a CLIJ operation, performance increases because of reuse of the compiled GPU code. We measured execution time and speed-up on two test systems: a consumer laptop (Intel i7-8650U CPU and an Intel UHD 620 GPU), and a professional workstation (two Intel Xeon Silver 4110 CPUs and an Nvidia Quadro P6000 GPU). We observed that tested CLIJ operations (Suppl. Listing 2) were up to about two orders of magnitude faster compared to their counter parts in ImageJ running on the CPU (Figure 1c).

**Figure 1:**
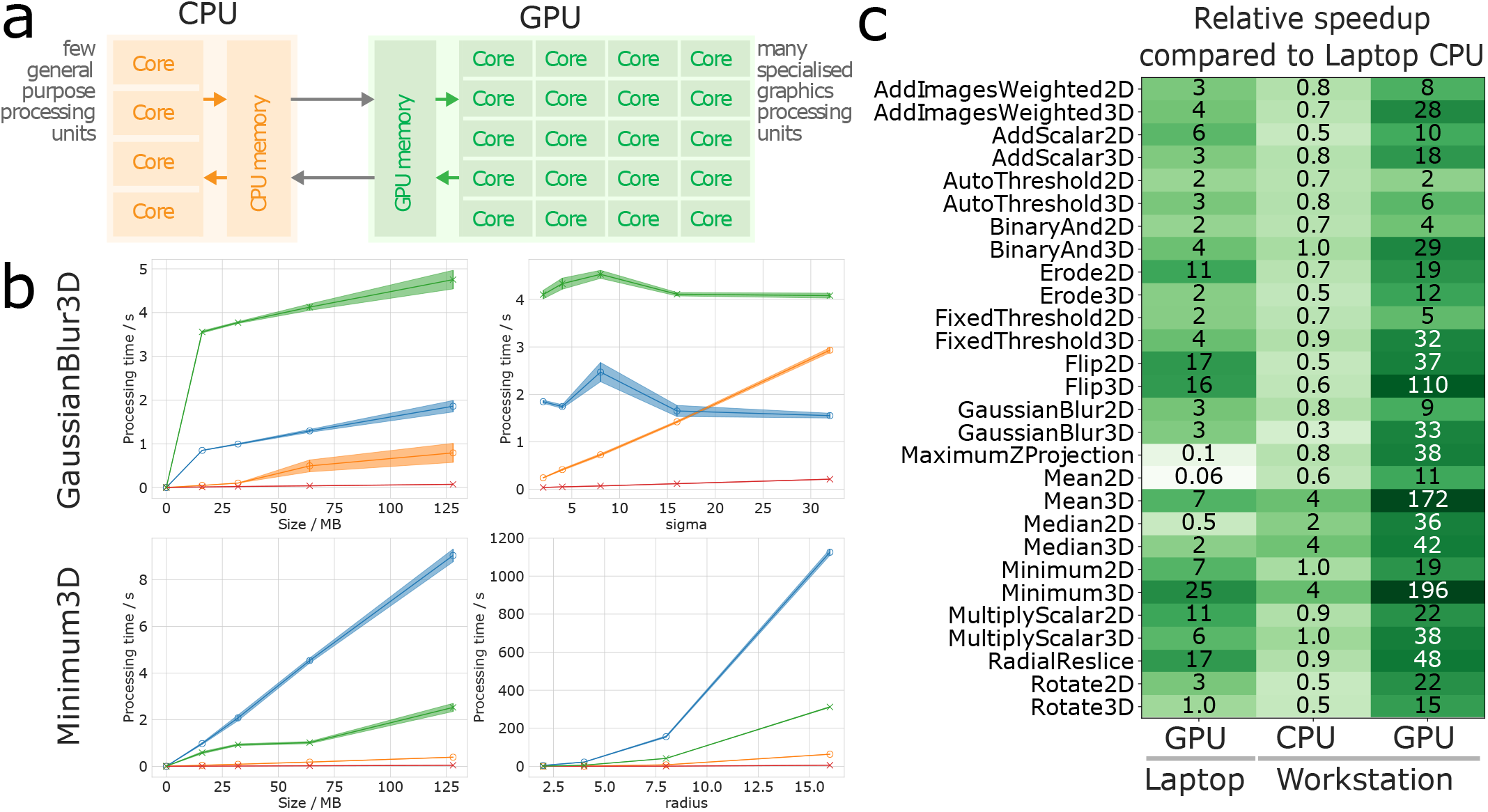
(**a**) GPU-acceleration schematically shows how many GPU cores potentially outperform a CPU with less cores. (**b**) Execution time of the Gaussian blur and minimum filter for different image sizes and parameters. Error bands denote the 99.9% confidence interval. (**c**) Overview of speed-up measurements when applied to 32 MB (2D) / 64 MB (3D) large 16-bit images with respect to computation times on a laptop CPU. Time measurements excluded memory transfer and compilation times.

To demonstrate the utility of CLIJ in practical biological image processing, we chose a multi-step example workflow (Suppl. Figure 4) operating on 3D light sheet microscopy data consisting of 300 time points of a early *Drosophila* embryo expressing histone-RFP to mark the nuclei. The workflow performs Difference-of-Gaussian filtering to reduce background signal and noise, projects the data from 3D to 2D and detects spots to count the nuclei. The processed image stack of each time point consists of 400×1024×121 voxels occupying 189 MB in memory. Since CLIJ operations are new implementations of existing ImageJ functions based on a different computing architecture, we determined how much the output of the GPU-based workflow and the corresponding CPU-based workflow were different. We observed minor absolute differences in the spot count result of 0.9±0.6 percent corresponding to a difference of about 22 in absolute spot count. Furthermore, we observed differences of 0.05±0.04 percent in spot count depending on which GPU hardware CLIJ was used (Suppl. Figure 5). While these differences may in practice be negligible (Suppl. Figure 6, Suppl. Video 1), we think users should be aware that they exist.

The whole time lapse was processed on our laptop within 2 hours and 44 minutes using ImageJ and 11 minutes using CLIJ. On our workstation, processing took 41 minutes using ImageJ and 5 minutes using CLIJ. Thus, these results show that using a consumer laptop, CLIJ enables a speed-up by a factor of 15. Compared to the laptop CPU, execution on the workstation GPU was 33 times faster. We would like to note that all measured runtimes depend on the executed workflow, the image data it is applied on, and the GPU hardware and its drivers. Hence, the precise speed-up of a given pipeline and hardware is hard to predict but will likely be similar in magnitude. We encourage users to consult our FAQ section online (Suppl. Listing 3) to learn more about optimal exploitation of GPU-accelerated image processing and benchmarking. Furthermore, excluding compilation time and file input/output time from the time measurement suggests that real-time image analysis becomes feasible: In a smart microscopy software application, where image data arrives in memory continuously directly from the acquiring camera and GPU code recompilation is not necessary, an estimation of cell count can be made from an image stack in less than 0.5 seconds using the presented CLIJ-based workflow.

Key feature of CLIJ is that it does not require any GPU programming skills, or specialized hardware to be executed. As it is based on the established OpenCL framework, it is not limited to CUDA-compatible GPU devices. The user can assemble CLIJ operations into GPU-accelerated image processing workflows in all programming languages available in Fiji (ImageJ Macro, ImageJ-Ops, BeanShell, JavaScript, Jython, Groovy, and Java). Users can start using CLIJ by simply modifying example code (Suppl. Code 1). Moreover, CLIJ operations can be recorded using ImageJ’s macro recorder and further modified in Fiji’s script editor where we added CLIJ-specific auto-completion functions and online help (Suppl. Figure 7). CLIJ can be used in the cloud or on computing clusters via ImageJ Jupyter notebooks or command-line interface. We tested CLIJ successfully on GPU-hardware from major vendors (Intel, AMD, Nvidia) and operating systems (Windows, MacOS, Linux). Finally, we facilitate providing additional GPU-accelerated operations to be used within the ImageJ ecosystem and extending CLIJ. Specifically, developers can deploy custom OpenCL code using the modern ImageJ2 plugin mechanism^10^ in order to add functionality to CLIJ. For potential CLIJ developers we provide a plugin template together with the full open source code of CLIJ and all data and scripts (Suppl. Listing 4) needed to support our findings at https://clij.github.io/

In summary, CLIJ makes it possible to speed up image processing workflows in Fiji to reduce processing time from hours to minutes. Furthermore, CLIJ allows general purpose realtime image processing, e.g. for smart microscopy applications. In order to facilitate adoption of this enabling technology, we have put special emphasis on documentation, code examples, interoperability, accessibility and user convenience. Therefore, CLIJ enables a wide range of imaging scientists with beginner-level programming experience to benefit from GPU-acceleration.

## Supporting information

Supplementary Code 1: CLIJ usage examples

Supplementary Figure 1: Processing time versus image size for 2D images

Supplementary Figure 2: Processing time versus image size for 3D images

Supplementary Figure 3: Processing time versus parameters

Supplementary Figure 4: Benchmarked workflow

Supplementary Figure 5: Differences between ImageJ and CLIJ workflows

Supplementary Figure 6: Workflow results

Supplementary Figure 7: CLIJ Screenshot

Supplementary Listing 1: CLIJ API Macro reference

Supplementary Listing 2: Benchmarked operations

Supplementary Listing 3: Frequently Asked Questions

Supplementary Listing 4: Used raw data and resulting figures

Supplementary Video 1: Results from processed time lapse for benchmarking workflow

## Acknowledgements

We would like to thank everybody who helped developing and testing CLIJ. In particular thanks goes to Bruno C. Vellutini (MPI-CBG), Curtis Rueden (UW-Madison LOCI), Damir Krunic (DKFZ), Daniel J. White (GE), Gaby G. Martins (IGC), Siân Culley (LMCB MRC), Giovanni Cardone (MPI Biochem), Jan Brocher (Biovoxxel), Johannes Girstmair (MPI-CBG), Juergen Gluch (Fraunhofer IKTS), Kota Miura, Laurent Thomas (Acquifer), Nico Stuurman (UCSF), Peter Haub, Pradeep Rajasekhar (Monash University), Tobias Pietzsch (MPI-CBG), Wilsom Adams (VU Biophotonics). This work was supported by the German Federal Ministry of Research and Education (BMBF) under the code 031L0044 (Sysbio II) and the German Research Foundation (DFG) under the code JU3110/1-1.

## Authors contributions

RH and LAR initiated the research. RH, LAR, PS, DS, AD, US and MW wrote the source code of CLIJ. RH and NM performed the image acquisition of the example data. EWM supervised the project. RH, LAR, US, MW, PT and FJ wrote the manuscript with input from all co-authors.

