## Supplementary Code 1: CLIJ usage examples for "CLIJ: GPU-accelerated image processing for everyone": automaticThreshold.bsh

```
// automaticThreshold.bsh
// ======================
//
// Apply an automatic threshold method to an image on the GPU
//
// Author: Robert Haase,
//         April 2019
//////////////////////////////////////////

import ij.IJ;
import net.haesleinhuepf.clij.CLIJ;


IJ.run("Close All");

// load example image
imp = IJ.openImage("http://wsr.imagej.net/images/blobs.gif");
imp.show();

// init GPU
clij = CLIJ.getInstance();

// push image to GPU
input = clij.push(imp);

// reserve memory for output, same size and type as input
output = clij.create(input);

// apply threshold method on GPU
clij.op().automaticThreshold(input, output, "Otsu");

// show result
clij.pullBinary(output).show();
```
