## Supplementary Code 1: CLIJ usage examples for "CLIJ: GPU-accelerated image processing for everyone": clij_micromanager.bsh

```
// This example script shows how to run CLIJ from MicroManager
// In order to get it run, copy the CLIJ jar-files from the release 
// (https://github.com/clij/clij/releases) in the 
// plugins directory of MicroManager. 
// Tested with MMSetup_64bit_2.0.0-gamma1_20190611
//
// Author: Robert Haase
// June 2019
/////////////////////////////////////////////////////////////////////

import net.haesleinhuepf.clij.CLIJ;
import ij.IJ;
import net.haesleinhuepf.clij.converters.implementations.ClearCLBufferToImagePlusConverter;
import net.haesleinhuepf.clij.converters.implementations.ImagePlusToClearCLBufferConverter;

// initialize CLIJ
clij = CLIJ.getInstance();
IJ.log(clij.clinfo());

// get some test data
imp = IJ.openImage("http://wsr.imagej.net/images/blobs.gif");
imp.show();

// send image to GPU and allocate some memory for result image
gpu_input = clij.push(imp);
gpu_output = clij.create(gpu_input);

// blur the image
clij.op().blur(gpu_input, gpu_output, 10, 10);

// get the result back from the GPU and show it
imp = clij.pull(gpu_output);
imp.show();

// clean up memory on the GPU
gpu_input.close();
gpu_output.close();
```
