## Supplementary Code 1: CLIJ usage examples for "CLIJ: GPU-accelerated image processing for everyone": maximumProjection.bsh

```
// This script shows how generate a maximum Z projection using CLIJ and BeanShell.
//
// Author: Robert Haase (@haesleinhuepf)
// April 2019
//

////////////////////////////////////////

import ij.IJ;
import net.haesleinhuepf.clij.CLIJ;

IJ.run("Close All");

// Init GPU
clij = CLIJ.getInstance();

// get some example data
imp = IJ.openImage("http://imagej.nih.gov/ij/images/t1-head.zip");

// create and fill memory in GPU
imageInput = clij.push(imp);
imageOutput = clij.create(new long[]{imageInput.getWidth(), imageInput.getHeight()}, imageInput.getNativeType());

// process the image
clij.op().maximumZProjection(imageInput, imageOutput);

// show the result
clij.show(imageOutput, "output");

// get the result back as variable
result = clij.pull(imageOutput);

IJ.run("Enhance Contrast", "saturated=0.35");
```
