## Supplementary Code 1: CLIJ usage examples for "CLIJ: GPU-accelerated image processing for everyone": CLIJMacroPluginServiceDemo.java

```
package net.haesleinhuepf.clij.examples.advanced;

import ij.IJ;
import ij.macro.ExtensionDescriptor;
import net.haesleinhuepf.clij.macro.CLIJHandler;
import net.haesleinhuepf.clij.macro.CLIJMacroPluginService;
import net.imagej.ImageJ;

import java.util.Arrays;

/**
 * The CLIJMacroPluginServiceDemo shows how to run CLJI Macro modules from Java.
 * <p>
 * Author: @haesleinhuepf
 * December 2018
 */
public class CLIJMacroPluginServiceDemo {
    public static void main(String... args) {
        ImageJ ij = new ImageJ();
        ij.ui().showUI();

        // get the plugin service from ImageJ
        CLIJMacroPluginService clijMacroPluginService = ij.get(CLIJMacroPluginService.class);

        // print out registered plugins / methods
        for (String name : clijMacroPluginService.getCLIJMethodNames()) {
            System.out.println(name);
        }

        // print some more details; namely parameters
        for (ExtensionDescriptor ed : CLIJHandler.getInstance().getExtensionFunctions()) {
            System.out.println(ed.name + " " + Arrays.toString(ed.argTypes));
        }

        // load image and push it to GPU
        IJ.open("https://github.com/clij/clij-docs/raw/master/src/main/resources/flybrain.tif");
        CLIJHandler.getInstance().handleExtension("CLIJ_push", new Object[] {"flybrain.tif"});

        // execute a processing CLIJ macro operation on it
        Object[] arguments = new Object[]{
                "flybrain.tif",
                "out",
                new Double(5),
                new Double(5),
                new Double(5)
        };
        CLIJHandler.getInstance().handleExtension("CLIJ_mean3DBox", arguments);

        // retrieve result
        CLIJHandler.getInstance().handleExtension("CLIJ_pull", new Object[] {"out"});

        // clean up
        CLIJHandler.getInstance().handleExtension("CLIJ_clear", new Object[]{});

    }
}
```
