## Supplementary Code 1: CLIJ usage examples for "CLIJ: GPU-accelerated image processing for everyone": CustomOpenCLDemo.java

```
package net.haesleinhuepf.clij.examples.advanced;

import ij.IJ;
import ij.ImageJ;
import ij.ImagePlus;
import net.haesleinhuepf.clij.CLIJ;
import net.haesleinhuepf.clij.clearcl.ClearCLBuffer;
import net.haesleinhuepf.clij.kernels.Kernels;
import net.imglib2.RandomAccessibleInterval;
import net.imglib2.img.display.imagej.ImageJFunctions;

import java.io.IOException;
import java.util.HashMap;
import java.util.Map;

/**
 * This example shows how to take input images, process them through
 * by two OpenCL kernels and show the result
 * <p>
 * Author: Robert Haase (http://haesleinhuepf.net) at MPI CBG (http://mpi-cbg.de)
 * February 2018
 */
public class CustomOpenCLDemo {

    public static void main(String... args) throws IOException {
        // Initialize ImageJ and example images
        new ImageJ();
        ImagePlus inputImp = IJ.openImage("https://github.com/clij/clij-docs/raw/master/src/main/resources/flybrain.tif");
        inputImp.show();

        // Startup OpenCL device, convert images to ClearCL format
        CLIJ clij = CLIJ.getInstance();

        ClearCLBuffer inputCLBuffer = clij.push(inputImp);
        ClearCLBuffer outputCLBuffer = clij.create(inputCLBuffer);

        // ---------------------------------------------------------------
        // Example step 1: Downsampling

        Map<String, Object> parameterMap = new HashMap<>();
        parameterMap.put("src", inputCLBuffer);
        parameterMap.put("dst", outputCLBuffer);
        parameterMap.put("factor_x", 0.5f);
        parameterMap.put("factor_y", 0.5f);
        parameterMap.put("factor_z", 1.f);

        clij.execute(Kernels.class, "downsampling.cl", "downsample_3d_nearest", parameterMap);

        // Convert/copy and show intermediate result
        RandomAccessibleInterval intermediateResult = clij.convert(outputCLBuffer, RandomAccessibleInterval.class);

        ImageJFunctions.show(intermediateResult);

        // ---------------------------------------------------------------
        // Example Step 2: Bluring
        HashMap<String, Object> blurParameterMap = new HashMap<>();
        blurParameterMap.put("Nx", 3);
        blurParameterMap.put("Ny", 3);
        blurParameterMap.put("Nz", 3);
        blurParameterMap.put("sx", 2.0f);
        blurParameterMap.put("sy", 2.0f);
        blurParameterMap.put("sz", 2.0f);
        // we reuse memory in the GPU by taking the result from the former
        // step as input here and the input from the former step as
        // output:
        blurParameterMap.put("src", outputCLBuffer);
        blurParameterMap.put("dst", inputCLBuffer);

        clij.execute(Kernels.class, "blur.cl", "gaussian_blur_image3d", blurParameterMap);

        // Convert and show final result
        RandomAccessibleInterval result = clij.convert(inputCLBuffer, RandomAccessibleInterval.class);

        ImageJFunctions.show(result);


    }
}
```
