## Supplementary Code 1: CLIJ usage examples for "CLIJ: GPU-accelerated image processing for everyone": DifferenceOfGaussianCLDemo.java

```
package net.haesleinhuepf.clij.examples.advanced;

import ij.IJ;
import ij.ImageJ;
import ij.ImagePlus;
import ij.plugin.Duplicator;
import net.haesleinhuepf.clij.CLIJ;
import net.haesleinhuepf.clij.clearcl.ClearCLBuffer;
import net.imglib2.RandomAccessibleInterval;
import net.imglib2.img.display.imagej.ImageJFunctions;
import net.imglib2.type.numeric.integer.UnsignedShortType;

import java.io.IOException;
import java.util.HashMap;
import java.util.Map;

/**
 * This
 * <p>
 * Author: Robert Haase (http://haesleinhuepf.net) at MPI CBG (http://mpi-cbg.de)
 * February 2018
 */
public class DifferenceOfGaussianCLDemo {
    public static void main(String... args) throws IOException {
        new ImageJ();
        ImagePlus inputImp = IJ.openImage("https://github.com/clij/clij-docs/raw/master/src/main/resources/flybrain.tif");

        inputImp = new Duplicator().run(inputImp, 25, 25);

        RandomAccessibleInterval<UnsignedShortType> input = ImageJFunctions.wrap(inputImp);

        RandomAccessibleInterval<UnsignedShortType> output = ImageJFunctions.wrap(new Duplicator().run(inputImp));

        ImageJFunctions.show(input);

        CLIJ clij = CLIJ.getInstance();

        // ---------------------------------------------------------------
        // Example 1: Flip image in X
        {
            ClearCLBuffer srcImage = clij.push(input);
            ClearCLBuffer dstImage = clij.push(output);

            Map<String, Object> parameterMap = new HashMap<>();
            parameterMap.put("src", srcImage);
            parameterMap.put("dst", dstImage);
            parameterMap.put("radius", 6);
            parameterMap.put("sigma_minuend", 1.5f);
            parameterMap.put("sigma_subtrahend", 3f);

            clij.execute("src/main/jython/differenceOfGaussian/differenceOfGaussian.cl", "subtract_convolved_images_2d_fast", parameterMap);

            ImagePlus result = clij.pull(dstImage);
            result.show();
            IJ.run(result,"Enhance Contrast", "saturated=0.35");

        }

    }
}
```
