## Supplementary Code 1: CLIJ usage examples for "CLIJ: GPU-accelerated image processing for everyone": AffineTransformDemo.java

```
package net.haesleinhuepf.clij.examples;

import ij.IJ;
import ij.ImageJ;
import ij.ImagePlus;
import net.haesleinhuepf.clij.CLIJ;
import net.haesleinhuepf.clij.clearcl.ClearCLBuffer;
import net.imglib2.realtransform.AffineTransform3D;

/**
 * AffineTransformDemo
 * <p>
 * <p>
 * <p>
 * Author: @haesleinhuepf
 * 01 2019
 */
public class AffineTransformDemo {
    public static void main(String... args) {
        new ImageJ();

        // load example image
        ImagePlus input = IJ.openImage("src/main/resources/flybrain.tif");

        // initialize GPU
        CLIJ clij = CLIJ.getInstance();

        // push image to GPU
        ClearCLBuffer inputOnGPU = clij.push(input);
        // create memory for target
        ClearCLBuffer resultOnGPU = clij.create(inputOnGPU);

        // define affine transform
        AffineTransform3D transform = new AffineTransform3D();
        transform.translate(-inputOnGPU.getWidth() / 2, -inputOnGPU.getHeight() / 2, -inputOnGPU.getDepth() / 2);
        transform.rotate(2, 45);
        transform.translate(inputOnGPU.getWidth() / 2, inputOnGPU.getHeight() / 2, inputOnGPU.getDepth() / 2);

        // apply transform
        clij.op().affineTransform3D(inputOnGPU, resultOnGPU, transform);

        // retrieve result or show it
        ImagePlus result = clij.pull(resultOnGPU);
        clij.show(resultOnGPU, "result");

        // free memory
        inputOnGPU.close();
        resultOnGPU.close();

    }
}
```
