## Supplementary Code 1: CLIJ usage examples for "CLIJ: GPU-accelerated image processing for everyone": ApplyVectorFieldDemo.java

```
package net.haesleinhuepf.clij.examples;

import ij.IJ;
import ij.ImageJ;
import ij.ImagePlus;
import ij.gui.NewImage;
import ij.gui.OvalRoi;
import net.haesleinhuepf.clij.CLIJ;
import net.haesleinhuepf.clij.clearcl.ClearCLBuffer;
import net.haesleinhuepf.clij.coremem.enums.NativeTypeEnum;
import net.imglib2.realtransform.AffineTransform2D;
import net.imglib2.realtransform.AffineTransform3D;

/**
 * This script demonstrates how to apply a vector field
 * to an image in order to transform it non-rigidly
 *
 * Author: @haesleinhuepf
 * June 2019
 */
public class ApplyVectorFieldDemo {
    public static void main(String... args) {

        new ImageJ();

        // get test image
        ImagePlus imp = IJ.openImage("https://samples.fiji.sc/blobs.png");
        IJ.run(imp,"32-bit", "");
        imp.setTitle("blobs");

        // create two images describing local shift
        ImagePlus shiftXimp = NewImage.createFloatImage("shiftX", 256, 254, 1, NewImage.FILL_BLACK);
        ImagePlus shiftYimp = NewImage.createFloatImage("shiftY", 256, 254, 1, NewImage.FILL_BLACK);

        // shift some of the pixels in X
        shiftXimp.setRoi(new OvalRoi(20, 98, 72, 68));
        IJ.run(shiftXimp, "Add...", "value=25");
        IJ.run(shiftXimp, "Select None", "");
        IJ.run(shiftXimp, "Gaussian Blur...", "sigma=15");
        IJ.run(shiftXimp, "Enhance Contrast", "saturated=0.35");

        // init GPU
        CLIJ clij = CLIJ.getInstance();

        // push input to GPU
        ClearCLBuffer blobsGPU = clij.push(imp);
        ClearCLBuffer shiftXgpu = clij.push(shiftXimp);
        ClearCLBuffer shiftYgpu = clij.push(shiftYimp);

        // allocate memory for results on GPU
        ClearCLBuffer rotatedShiftX = clij.create(blobsGPU);
        ClearCLBuffer result = clij.create(blobsGPU);
        ClearCLBuffer resultStack = clij.create(new long[]{256, 254, 36}, NativeTypeEnum.Float);

        for (int i = 0; i < 36; i++) {
            // define affine transform
            AffineTransform2D at = new AffineTransform2D();
            at.translate(-imp.getWidth() / 2, -imp.getHeight() / 2, 0);
            at.rotate((double)i * 10 / 180 * Math.PI);
            at.translate(imp.getWidth() / 2, imp.getHeight() / 2, 0);

            // rotate vector field
            clij.op().affineTransform2D(shiftXgpu, rotatedShiftX, at);

            // apply vector field / transform
            clij.op().applyVectorfield(blobsGPU, rotatedShiftX, shiftYgpu, result);

            // put resulting 2D image in the right plane
            clij.op().copySlice(result, resultStack, i);
        }


        // get result back from GPU
        ImagePlus resultImp = clij.pull(resultStack);
        IJ.run(resultImp , "Invert LUT", "");
        resultImp.setDisplayRange(0, 256);
        resultImp.show();

        // clean up memory
        blobsGPU.close();
        shiftXgpu.close();
        shiftYgpu.close();
        rotatedShiftX.close();
        result.close();
        resultStack.close();

    }
}
```
