## Supplementary Code 1: CLIJ usage examples for "CLIJ: GPU-accelerated image processing for everyone": BackgroundSubtractionandThresholdingDemo.java

```
package net.haesleinhuepf.clij.examples;

import ij.IJ;
import ij.ImagePlus;
import net.haesleinhuepf.clij.CLIJ;
import net.haesleinhuepf.clij.clearcl.ClearCLBuffer;
import net.haesleinhuepf.clij.kernels.Kernels;

/**
 * * Mean filter for background determination,
 * * background subtractionand
 * * thresholding
 *
 * Author: @haesleinhuepf
 * May 2019
 */
public class BackgroundSubtractionandThresholdingDemo {
    public static void main(String... args) {
        ImagePlus imp = IJ.openImage("src/main/resources/droso_crop.tif");

        CLIJ clij = CLIJ.getInstance();

        // conversion
        ClearCLBuffer input = clij.push(imp);
        ClearCLBuffer output = clij.create(input);
        ClearCLBuffer background = clij.create(input);
        ClearCLBuffer backgroundSubtracted = clij.create(input);

        // blur
        clij.op().meanBox(input, background, 2, 2, 2);

        // background subtraction
        clij.op().subtract(input, background, backgroundSubtracted);

        // threshold
        clij.op().automaticThreshold(backgroundSubtracted, output, "Otsu");

        // show results
        clij.show(input, "original");
        clij.show(output, "mask");

        // free memory afterwards
        input.close();
        output.close();
        background.close();
        backgroundSubtracted.close();
    }
}
```
