## Supplementary Code 1: CLIJ usage examples for "CLIJ: GPU-accelerated image processing for everyone": BenchmarkingWorkflowDemo.java

```
package net.haesleinhuepf.clij.examples;

import ij.IJ;
import ij.ImagePlus;
import ij.Prefs;
import ij.plugin.Duplicator;
import ij.plugin.ImageCalculator;
import net.haesleinhuepf.clij.CLIJ;
import net.haesleinhuepf.clij.clearcl.ClearCLBuffer;
import net.imagej.ImageJ;
import net.imglib2.IterableInterval;
import net.imglib2.RandomAccessibleInterval;
import net.imglib2.algorithm.neighborhood.DiamondShape;
import net.imglib2.img.Img;
import net.imglib2.img.display.imagej.ImageJFunctions;
import net.imglib2.roi.Regions;
import net.imglib2.type.BooleanType;
import net.imglib2.type.numeric.RealType;
import net.imglib2.type.numeric.integer.UnsignedByteType;
import net.imglib2.util.Pair;
import net.imglib2.view.Views;

import java.io.IOException;

/**
 * This class (rather its main method) applies a simple pipeline to a given image. It blurs it, creates a mask by
 * thresholding, does some erosion and dilatation to make the edges smoother and finally sets all background pixels in
 * the original image to zero. This pipeline is executed in the ImageJ1 way and in OpenCL to compare performance and
 * results.
 * <p>
 * Author: Robert Haase (http://haesleinhuepf.net) at MPI CBG (http://mpi-cbg.de)
 * March 2018
 */
public class BenchmarkingWorkflowDemo {

    private static ImageJ ij;
    private static CLIJ clij;
    private static double sigma = 3;

    public static void main(String... args) throws IOException {

        // ---------------------------------------
        // Initialize ImageJ, CLIJ and test image
        ij = new ImageJ();
        ij.ui().showUI();
        clij = CLIJ.getInstance();

        input = IJ.openImage("https://github.com/clij/clij-docs/raw/master/src/main/resources/flybrain.tif");
        input.show();
        IJ.run(input, "8-bit", "");
        img = ImageJFunctions.wrapReal(input);

        for (int i = 0; i < 100; i++) {
            // ---------------------------------------
            // three test scenarios
            long timestamp = System.currentTimeMillis();
            demoImageJ1();
            System.out.println("The ImageJ1 way took " + (System.currentTimeMillis() - timestamp) + " msec");

            timestamp = System.currentTimeMillis();
            demoImageJ2();
            System.out.println("The ImageJ2 way took " + (System.currentTimeMillis() - timestamp) + " msec");

            timestamp = System.currentTimeMillis();
            demoCLIJ();
            System.out.println("The CLIJ way took " + (System.currentTimeMillis() - timestamp) + " msec");
            // ---------------------------------------
        }
    }

    private static ImagePlus input;
    private static Img img;

    private static void demoImageJ1() {
        ImagePlus copy = new Duplicator().run(input, 1, input.getNSlices());
        Prefs.blackBackground = false;
        IJ.run(copy, "Gaussian Blur 3D...", "x=" + sigma + " y=" + sigma + " z=" + sigma + "");
        IJ.setRawThreshold(copy, 100, 255, null);
        IJ.run(copy, "Convert to Mask", "method=Default background=Dark");
        IJ.run(copy, "Erode", "stack");
        IJ.run(copy, "Dilate", "stack");
        IJ.run(copy, "Divide...", "value=255 stack");
        ImageCalculator calculator = new ImageCalculator();

        ImagePlus result = calculator.run("Multiply create stack", copy, input);
        result.show();
    }

    private static <T extends RealType<T>, B extends BooleanType<B>> void demoImageJ2() {

        // Thanks to @imagejan, @tpietzsch and @Awalter from forum.imagej.net

        UnsignedByteType threshold = new UnsignedByteType();
        threshold.setReal(100);
        RandomAccessibleInterval gauss = ij.op().filter().gauss(img, sigma);

        // Gaussian blur
        IterableInterval blurredIi = ij.op().threshold().apply(Views.iterable(gauss), threshold);
        RandomAccessibleInterval blurredImg = makeRai(blurredIi);

        // erode
        IterableInterval erodedIi = ij.op().morphology().erode(blurredImg, new DiamondShape(1));
        RandomAccessibleInterval erodedImg = makeRai(erodedIi);

        // dilate
        IterableInterval dilatedIi = ij.op().morphology().dilate(erodedImg, new DiamondShape(1));
        RandomAccessibleInterval dilatedImg = makeRai(dilatedIi);

        // mask original image
        Img output = ij.op().create().img(img);
        IterableInterval<Pair<T, T>> sample = Regions.sample(
                Regions.iterable(dilatedImg),
                Views.pair(output, img)
        );
        sample.forEach(pair -> pair.getA().set(pair.getB()));

        ImageJFunctions.show(output);

    }

    private static RandomAccessibleInterval makeRai(IterableInterval ii) {
        RandomAccessibleInterval rai;
        if (ii instanceof RandomAccessibleInterval) {
            rai = (RandomAccessibleInterval) ii;
        } else {
            rai = ij.op().create().img(ii);
            ij.op().copy().iterableInterval((Img) rai, ii);
        }
        return rai;
    }

    private static void demoCLIJ() throws IOException {
        ClearCLBuffer input = clij.push(img);
        ClearCLBuffer flip = clij.create(input.getDimensions(), input.getNativeType());
        ClearCLBuffer flop = clij.create(input.getDimensions(), input.getNativeType());

        clij.op().blur(input, flop, (float) sigma, (float) sigma, (float) sigma);

        clij.op().threshold(flop, flip, 100.0f);

        clij.op().erodeSphere(flip, flop);
        clij.op().dilateSphere(flop, flip);

        clij.op().multiplyImages(flop, input, flip);

        clij.pull(flip).show();

        flip.close();
        flop.close();
        input.close();
    }
}
```
