## Supplementary Code 1: CLIJ usage examples for "CLIJ: GPU-accelerated image processing for everyone": BlurDemo.java

```
package net.haesleinhuepf.clij.examples;

import ij.IJ;
import ij.ImagePlus;
import net.haesleinhuepf.clij.CLIJ;
import net.haesleinhuepf.clij.clearcl.ClearCLBuffer;

/**
 * Apply a Gaussian blur to an image on the GPU
 *
 * Author: @haesleinhuepf
 * June 2019
 */
public class BlurDemo {
    public static void main(String... args) {
        ImagePlus imp = IJ.openImage("https://samples.fiji.sc/blobs.png");

        CLIJ clij = CLIJ.getInstance();

        // conversion
        ClearCLBuffer input = clij.push(imp);
        ClearCLBuffer output = clij.create(input);

        // blur
        float sigma = 2;
        clij.op().blur(input, output, sigma, sigma);

        ImagePlus result = clij.pull(output);
        result.show();

        // free memory afterwards
        input.close();
        output.close();
    }
}
```
