## Supplementary Code 1: CLIJ usage examples for "CLIJ: GPU-accelerated image processing for everyone": CLIJImageJOpsCombinationDemo.java

```
package net.haesleinhuepf.clij.examples;

import ij.IJ;
import ij.ImagePlus;
import net.haesleinhuepf.clij.CLIJ;
import net.haesleinhuepf.clij.clearcl.ClearCLBuffer;
import net.haesleinhuepf.clij.coremem.enums.NativeTypeEnum;
import net.imagej.ImageJ;
import net.imglib2.RandomAccessibleInterval;
import net.imglib2.algorithm.labeling.ConnectedComponents;
import net.imglib2.img.display.imagej.ImageJFunctions;
import net.imglib2.roi.labeling.ImgLabeling;
import net.imglib2.roi.labeling.LabelRegions;
import net.imglib2.type.logic.BitType;
import net.imglib2.type.numeric.IntegerType;

/**
 * CLIJImageJOpsCombinationDemo
 * <p>
 * <p>
 * <p>
 * Author: @haesleinhuepf
 * 02 2019
 */
public class CLIJImageJOpsCombinationDemo {
    public static void main(String... args) {
        // define parameters
        String filename = "src/main/resources/blobs.tif";
        float sigma = 3;

        // load test data
        ImageJ ij = new ImageJ();
        ij.ui().showUI();
        ImagePlus imp = IJ.openImage(filename);
        imp.show();

        // reserve memory on GPU / transfer images
        CLIJ clij = CLIJ.getInstance();
        ClearCLBuffer input = clij.push(imp);
        ClearCLBuffer blurred = clij.create(input);
        ClearCLBuffer thresholded = clij.createCLBuffer(input.getDimensions(), NativeTypeEnum.Byte);

        // Gaussian blur
        clij.op().blur(input, blurred, sigma, sigma, sigma);

        // Apply Otsu threshold
        clij.op().automaticThreshold(blurred, thresholded, "Otsu");

        // convert back from GPU
        ImagePlus result = clij.pull(thresholded);
        RandomAccessibleInterval<BitType> binaryRai = clij.pullBinaryRAI(thresholded);

        // clean up
        input.close();
        blurred.close();
        thresholded.close();

        // show result from GPU
        result.show();

        //continue with Ops
        ImgLabeling cca = ij.op().labeling().cca(binaryRai, ConnectedComponents.StructuringElement.FOUR_CONNECTED);

        LabelRegions<IntegerType> regions = new LabelRegions(cca);

        System.out.print("Number of objects found: " + regions.getExistingLabels().size());

        // show results of labelling on CPU
        ImageJFunctions.show(cca.getIndexImg());

    }
}
```
