## Supplementary Code 1: CLIJ usage examples for "CLIJ: GPU-accelerated image processing for everyone": ClinfoDemo.java

```
package net.haesleinhuepf.clij.examples;

import net.haesleinhuepf.clij.CLIJ;

/**
 * This demo just shows how to output all information known on available
 * OpenCL devices to stdout.
 * <p>
 * Author: Robert Haase (http://haesleinhuepf.net) at MPI CBG (http://mpi-cbg.de)
 * February 2018
 */
public class ClinfoDemo {
    public static void main(String... args) {

        System.out.println(CLIJ.clinfo());

    }
}
```
