## Supplementary Code 1: CLIJ usage examples for "CLIJ: GPU-accelerated image processing for everyone": CreateObjectOutlinesDemo.java

```
package net.haesleinhuepf.clij.examples;

import ij.IJ;
import ij.ImageJ;
import ij.ImagePlus;
import net.haesleinhuepf.clij.CLIJ;
import net.haesleinhuepf.clij.clearcl.ClearCLBuffer;

/**
 * AutoThresholdDemo
 *
 * This java example shows how to apply an automatic
 * threshold method to an image and get the segmentations
 * outline as binary image in the GPU.
 *
 * Author: @haesleinhuepf
 * June 2019
 */
public class CreateObjectOutlinesDemo {
    public static void main(String[] args) {
        new ImageJ();

        // get test image
        ImagePlus imp = IJ.openImage("https://samples.fiji.sc/blobs.png");
        IJ.run(imp,"32-bit", "");
        imp.setTitle("blobs");


        // init GPU
        CLIJ clij = CLIJ.getInstance();

        // push image to GPU and allocate memory for result
        ClearCLBuffer blobsGPU = clij.push(imp);
        ClearCLBuffer thresholded = clij.create(blobsGPU);
        ClearCLBuffer extended = clij.create(blobsGPU);
        ClearCLBuffer outline= clij.create(blobsGPU);


        // apply threshold
        clij.op().automaticThreshold(blobsGPU, thresholded, "Otsu");

        clij.op().dilateBox(thresholded, extended);

        clij.op().binaryXOr(thresholded, extended, outline);


        // show result
        ImagePlus result = clij.pullBinary(outline);
        result.show();

        // cleanup
        blobsGPU.close();
        thresholded.close();
        extended.close();
        outline.close();

    }
}
```
