## Supplementary Code 1: CLIJ usage examples for "CLIJ: GPU-accelerated image processing for everyone": LocalThresholdDemo.java

```
package net.haesleinhuepf.clij.examples;

import ij.IJ;
import ij.ImagePlus;
import net.haesleinhuepf.clij.CLIJ;
import net.haesleinhuepf.clij.clearcl.ClearCLBuffer;
import net.haesleinhuepf.clij.kernels.Kernels;

/**
 * LocalThresholdDemo
 * <p>
 * <p>
 * <p>
 * Author: @haesleinhuepf
 * 06 2018
 */
public class LocalThresholdDemo {
    public static void main(String... args) {
        ImagePlus imp = IJ.openImage("src/main/resources/droso_crop.tif");

        CLIJ clij = CLIJ.getInstance();

        // conversion
        ClearCLBuffer input = clij.push(imp);
        ClearCLBuffer output = clij.create(input);
        ClearCLBuffer temp = clij.create(input);

        // blur
        Kernels.blur(clij, input, temp, 2f, 2f, 2f);

        // local threshold
        Kernels.localThreshold(clij, input, output, temp);

        Kernels.erodeSphere(clij, output, temp);
        Kernels.erodeSphere(clij, temp, output);

        // show results
        clij.show(input, "original");
        clij.show(output, "mask");

        input.close();
        output.close();
        temp.close();
    }
}
```
