## Supplementary Code 1: CLIJ usage examples for "CLIJ: GPU-accelerated image processing for everyone": MotionCorrectionDemo.java

```
package net.haesleinhuepf.clij.examples;

import ij.IJ;
import ij.ImageJ;
import ij.ImagePlus;
import net.haesleinhuepf.clij.CLIJ;
import net.haesleinhuepf.clij.clearcl.ClearCLBuffer;
import net.haesleinhuepf.clij.macro.modules.Clear;
import net.imglib2.realtransform.AffineTransform3D;

/**
 * MotionCorrectionDemo
 * <p>
 * <p>
 * <p>
 * Author: @haesleinhuepf
 * 06 2019
 */
public class MotionCorrectionDemo {
    public static void main(String[] args) {
        new ImageJ();

        // define move to correct
        ImagePlus imp = IJ.openImage("src/main/resources/motion_correction_Drosophila_DSmanila1.tif");
        IJ.run(imp, "32-bit", "");
        imp.show();

        // define a threshold to differentiate object and background
        float threshold = 50;

        // Init GPU
        CLIJ clij = CLIJ.getInstance();

        // push input to GPU
        ClearCLBuffer input = clij.push(imp);

        // create some slice images on the GPU to work on
        ClearCLBuffer inputSlice = clij.create(new long[]{input.getWidth(), input.getHeight()}, input.getNativeType());
        ClearCLBuffer thresholded = clij.create(inputSlice);
        ClearCLBuffer shifted = clij.create(inputSlice);

        double formerX = 0;
        double formerY = 0;

        // process all slices; only the first stays where it is
        for (int z = 0; z  < imp.getNSlices(); z++) {
            System.out.println("z: " + z);
            clij.op().copySlice(input, inputSlice, z);

            // determine center of mass

            clij.op().threshold(inputSlice, thresholded, threshold);
            double[] centerOfMass = clij.op().centerOfMass(thresholded);

            if (z > 0) {

                // determine shift
                double deltaX = centerOfMass[0] - formerX;
                double deltaY = centerOfMass[1] - formerY;

                // apply translation transformation
                AffineTransform3D at = new AffineTransform3D();
                at.translate(-deltaX, -deltaY, 0);

                clij.op().affineTransform(inputSlice, shifted, at);

                // copy result back
                clij.op().copySlice(shifted, input, z);

            } else {
                formerX = centerOfMass[0];
                formerY = centerOfMass[1];
            }
        }

        // show result
        ImagePlus result = clij.pull(input);
        result.setDisplayRange(imp.getDisplayRangeMin(), imp.getDisplayRangeMax());
        result.setTitle("Motion corrected");
        result.show();

        input.close();
        inputSlice.close();
        thresholded.close();
        shifted.close();
    }
}
```
