## Supplementary Code 1: CLIJ usage examples for "CLIJ: GPU-accelerated image processing for everyone": MultipleGPUDemo.java

```
package net.haesleinhuepf.clij.examples;

import com.sun.org.apache.xpath.internal.operations.Mult;
import ij.IJ;
import ij.ImagePlus;
import net.haesleinhuepf.clij.CLIJ;
import net.haesleinhuepf.clij.clearcl.ClearCLBuffer;

import java.util.ArrayList;

/**
 * This demo runs on a computer with multiple GPUs. It processes images in parallel on both GPUs.
 *
 * Author: Robert Haase, MPI CBG
 * June 2019
 */
public class MultipleGPUDemo {

    public static void main(String... args) {
        new MultipleGPUDemo().run();
    }

    public void run() {
        ArrayList<Integer> deviceIndices = new ArrayList<Integer>();

        int count = 0;
        // print out all GPU names
        for (String deviceName : CLIJ.getAvailableDeviceNames()) {
            System.out.println(deviceName);
            if (deviceName.toLowerCase().contains("gpu") || // Intel HD GPUs
                deviceName.toLowerCase().contains("gfx") || // AMD Ryzen GPUs
                deviceName.toLowerCase().contains("geforce") // NVidia Geforce GPUs
            ) {
                deviceIndices.add(count);
            }
            count++;
        }

        if (deviceIndices.size() < 2) {
            System.out.println("This computer doesn't have several GPUs. This demo needs several GPUs.");
            return;
        }

        // get CLIJ instances operating on different GPUs
        CLIJ clijGPU1 = new CLIJ(deviceIndices.get(0));
        CLIJ clijGPU2 = new CLIJ(deviceIndices.get(1));

        ImagePlus imp = IJ.openImage("src/main/resources/droso_crop.tif");

        // create multiple processors using different GPUs
        Processor processorGPU1 = new Processor(clijGPU1, imp);
        Processor processorGPU2 = new Processor(clijGPU2, imp);

        // start processing
        processorGPU1.start();
        processorGPU2.start();

        // wait until processing is done on both GPUs
        try {
            processorGPU1.join();
            processorGPU2.join();

            Thread.sleep(5000);
        } catch (InterruptedException e) {
            e.printStackTrace();
        }
    }

    class Processor extends Thread {
        CLIJ clij;
        ImagePlus imp;
        public Processor(CLIJ clij, ImagePlus imp) {
            this.clij = clij;
            this.imp = imp;
        }

        @Override
        public void run() {
            String gpuName = clij.getGPUName();
            log(gpuName + " start");

            // send an image to GPU and allocate memory for another image
            ClearCLBuffer input = clij.push(imp);
            ClearCLBuffer output = clij.create(input);
            log(gpuName + " copy done");

            // apply a Gaussian blur
            float sigma = 25;
            clij.op().blur(input, output, sigma, sigma, sigma);
            log(gpuName + " blur done");

            // show results
            clij.show(output, "Result on " + clij.getGPUName());
            log(gpuName + " finished");

            // cleanup memory
            input.close();
            output.close();
        }
    }

    private void log(String text) {
        synchronized (this) {
            System.out.println("" + System.currentTimeMillis() + "\t" + text);
        }
    }
}
```
