## Supplementary Code 1: CLIJ usage examples for "CLIJ: GPU-accelerated image processing for everyone": maximumProjection.js

```
// This script shows how generate a maximum Z projection using CLIJ and javascript.
//
// Author: Robert Haase (@haesleinhuepf)
// April 2019
//

////////////////////////////////////////
importClass(Packages.ij.IJ);
importClass(Packages.net.haesleinhuepf.clij.CLIJ);

IJ.run("Close All");

// Init GPU
clij = CLIJ.getInstance();

// get some example data
imp = IJ.openImage("http://imagej.nih.gov/ij/images/t1-head.zip");

// create and fill memory in GPU
imageInput = clij.push(imp);
dimensions = [imageInput.getWidth(), imageInput.getHeight()];
imageOutput = clij.create(dimensions, imageInput.getNativeType());

// process the image
clij.op().maximumZProjection(imageInput, imageOutput);

// show the result
clij.show(imageOutput, "output");

// get the result back as variable
result = clij.pull(imageOutput);

IJ.run("Enhance Contrast", "saturated=0.35");
```
