## Supplementary Code 1: CLIJ usage examples for "CLIJ: GPU-accelerated image processing for everyone": automaticThreshold.py

```
# automaticThreshold.py
# =====================
#
# Apply an automatic threshold method to an image on the GPU
#
# Author: Robert Haase,
#         April 2019
#########################################

from ij import IJ;
from net.haesleinhuepf.clij import CLIJ;


IJ.run("Close All");

# load example image
imp = IJ.openImage("http://wsr.imagej.net/images/blobs.gif");
imp.show();

# init GPU
clij = CLIJ.getInstance();

# push image to GPU
input = clij.push(imp);

# reserve memory for output, same size and type as input
output = clij.create(input);

# apply threshold method on GPU
clij.op().automaticThreshold(input, output, "Otsu");

# show result
clij.pull(output).show();
IJ.setMinAndMax(0, 1);
```
