## Supplementary Code 1: CLIJ usage examples for "CLIJ: GPU-accelerated image processing for everyone": differenceOfGaussian.py

```
# this example ImageJ/Fiji jython script shows how to apply a
# difference of gaussian (DoG) filter to an image using an
# OpenCL kernel.
#
# Author: Robert Haase (@haesleinhuepf)
# March 2018
#

from net.haesleinhuepf.clij import CLIJ;
from net.imglib2.view import Views;
from ij import IJ;
from java.lang import Float
import os;
import inspect

# retrieve the folder where this script is located (thanks to @mountain_man from the ImageJ forum)
filesPath = os.path.dirname(os.path.abspath(inspect.getsourcefile(lambda:0))) + "/"

# take the current image which is open in ImageJ
imp = IJ.openImage("http://imagej.nih.gov/ij/images/t1-head.zip");

# initialize ClearCL context and convenience layer
clij = CLIJ.getInstance();

# convert imglib2 image to CL images (ready for the GPU)
inputCLImage = clij.push(imp);
tempCLImage = clij.create([inputCLImage.getWidth(), inputCLImage.getHeight()], inputCLImage.getNativeType());
outputCLImage = clij.create([inputCLImage.getWidth(), inputCLImage.getHeight()], inputCLImage.getNativeType());

# crop out a center plane of the 3D data set
clij.op().copySlice(inputCLImage, tempCLImage, 64);

# apply a filter to the image using ClearCL / OpenCL
clij.execute(filesPath + "differenceOfGaussian.cl", "subtract_convolved_images_2d_fast", {
"src":tempCLImage,
"dst":outputCLImage,
"radius":6,
"sigma_minuend":Float(1.5),
"sigma_subtrahend":Float(3)});

# convert the result back to imglib2 and show it
result = clij.pull(outputCLImage);
result.show();
IJ.run("Enhance Contrast", "saturated=0.35");

# clean up
inputCLImage.close();
tempCLImage.close();
outputCLImage.close();
```
