## Supplementary Code 1: CLIJ usage examples for "CLIJ: GPU-accelerated image processing for everyone": maximumProjection.py

```
# This script shows how generate a maximum Z projection using CLIJ and jython.
#
# Author: Robert Haase (@haesleinhuepf)
# March 2018
#

#######################################

from ij import IJ;
from ij import ImagePlus;
from net.haesleinhuepf.clij import CLIJ;

# Init GPU
clij = CLIJ.getInstance();

# get some example data
imp = IJ.openImage("http://imagej.nih.gov/ij/images/t1-head.zip");

# create and fill memory in GPU
imageInput = clij.push(imp);
imageOutput = clij.create([imageInput.getWidth(), imageInput.getHeight()], imageInput.getNativeType());

# process the image
clij.op().maximumZProjection(imageInput, imageOutput);

# show the result
clij.show(imageOutput, "output");

# get the result back as variable
result = clij.pull(imageOutput);
```
