## Supplementary Code 1: CLIJ usage examples for "CLIJ: GPU-accelerated image processing for everyone": mean_detailed_comparison_IJ_CLIJ_radius.ijm

```
// CLIJ example macro: mean_detailed_comparison_IJ_CLIJ_radius.ijm
//
// This macro shows differences between ImageJ and CLIJ when executing a mean filter.
// Starting from a 2D image where a single pixel is set to 255, mean filters with
// different radii are applied.
//
// Author: Robert Haase
// June 2019
// ---------------------------------------------

run("Close All");


for (radius = 1; radius < 5; radius++) {
	// -----------------------------------------------------------------
	
	// create 2D example image
	input = "example2d";
	newImage(input, "8-bit black", 10, 10, 1);
	makeRectangle(4, 5, 1, 1);
	run("Add...", "value=255");
	run("Select None");
	
	// visualise input image
	zoom(10);
	setWindowPosition(0, 0);
	setMinAndMax(0, 1);
	makeOverlay();
	
	// init GPU
	run("CLIJ Macro Extensions", "cl_device=");
	
	// do operation on GPU
	outputGPU = "mean2d_GPU (r=" + radius + ")";
	Ext.CLIJ_push(input);
	Ext.CLIJ_mean2DSphere(input, outputGPU, radius, radius);
	Ext.CLIJ_pull(outputGPU);
	
	// visualise result
	zoom(10);
	setWindowPosition(radius, 0);
	setMinAndMax(0, 1);
	
makeOverlay();
	
	// do operation on CPU
	selectWindow(input);
	run("Duplicate...", "title=[mean2D_CPU (r=" + radius + ")]");
	run("Mean...", "radius=" + radius);
	
	// visualise result
	zoom(10);
	setWindowPosition(radius, 1);
	setMinAndMax(0, 1);
	
makeOverlay();
}
// -----------------------------------------------------------------

function zoom(count) {
	for (i = 0; i < count; i++) {	
		run("In [+]");
	}	
}

function setWindowPosition(x, y) {
	setLocation(x * 330, y * 370);
}

function makeOverlay() {
	for (y = 1; y <= 9; y += 2) {
		makeRectangle(0, y, 10, 1);
		Roi.setStrokeColor("gray");
		run("Add Selection...");
	}
	for (x = 1; x <= 9; x += 2) {
		makeRectangle(x, 0, 1, 10);
		Roi.setStrokeColor("gray");
		run("Add Selection...");	
	}
	run("Select None");
}
```
