## Supplementary Code 1: CLIJ usage examples for "CLIJ: GPU-accelerated image processing for everyone": addImages.ijm

```
// CLIJ example macro: addImages.ijm
//
// This macro shows how add images of different bit-type
//
// Author: Robert Haase
// January 2019
// ---------------------------------------------

run("Close All");

// Get test data
run("Blobs (25K)");
run("8-bit");
rename("original");
getDimensions(width, height, channels, slices, frames)

newImage("background", "16-bit ramp", width, height, slices);

// init GPU
run("CLIJ Macro Extensions", "cl_device=");
Ext.CLIJ_clear();

// push images to GPU
Ext.CLIJ_push("original");
Ext.CLIJ_push("background");


// cleanup imagej
run("Close All");

// create output image with 32 bits
Ext.CLIJ_create2D("originalWithBackground", width, height, 32);

// add images

Ext.CLIJ_addImagesWeighted("original", "background", "originalWithBackground", 1, 0.01);

// show result
Ext.CLIJ_pull("originalWithBackground");
run("Invert LUT");
```
