## Supplementary Code 1: CLIJ usage examples for "CLIJ: GPU-accelerated image processing for everyone": affineTransform.ijm

```
// CLIJ example macro: affineTransform.ijm
//
// This macro shows how to apply an affine transform in the GPU.
//
// Author: Robert Haase
// January 2019
// ---------------------------------------------

run("Close All");

// Get test data
run("Blobs (25K)");
//open("C:/structure/data/blobs.gif");


run("32-bit"); // interplation works better with float images
rename("original");

// init GPU
run("CLIJ Macro Extensions", "cl_device=");
Ext.CLIJ_clear();

// push images to GPU
Ext.CLIJ_push("original");

// cleanup imagej
run("Close All");

transform = "center ";
transform = transform + " rotate=45"; // degrees
transform = transform + " scaleX=2"; // relative zoom factor
transform = transform + " translateY=25"; // pixels
transform = transform + " -center";

Ext.CLIJ_affineTransform2D("original", "target", transform);

// show result
Ext.CLIJ_pull("target");
run("Invert LUT");
```
