## Supplementary Code 1: CLIJ usage examples for "CLIJ: GPU-accelerated image processing for everyone": allocateBigImages.ijm

```
// CLIJ example macro: allocateBigImages.ijm
//
// This macro shows how to process/handle big images in the GPU.
// Basically: They have to pushed in smaller blocks.
//
// Author: Robert Haase
// December 2018
// ---------------------------------------------

run ("Close All");


// Get test data
//open("C:/structure/data/blobs.gif");
run("Blobs (25K)");

// define image names
input = "input";
bigStack = "bigStack";
crop = "crop";

// Init GPU
run("CLIJ Macro Extensions", "cl_device=");
Ext.CLIJ_clear();

// push images to GPU
rename(input);
Ext.CLIJ_push(input);

// CleanUp ImageJ
close();

// create an 8 GB image in GPU memory
Ext.CLIJ_create3D(bigStack, 2048, 2048, 1000, 16);


for (i = 0; i < 10; i++) {
	// fill the image with content
	Ext.CLIJ_copySlice(input, bigStack, i);
}
Ext.CLIJ_crop3D(bigStack, crop, 0, 0, 0, 150, 150, 10);

// Get results back from GPU
Ext.CLIJ_pull(crop);

// report about what's allocated in the GPU memory
Ext.CLIJ_reportMemory();

// Cleanup by the end
Ext.CLIJ_clear();
```
