## Supplementary Code 1: CLIJ usage examples for "CLIJ: GPU-accelerated image processing for everyone": applyVectorField.ijm

```
// This script demonstrates how to apply a vector field
// to an image in order to transform it non-rigidly
//
// Author: Robert Haase,
// March 2019
//

run("Close All");

// get test image
run("Blobs (25K)");
run("32-bit");
rename("blobs");

// create two images describing local shift
newImage("shiftX", "32-bit black", 256, 254, 1);
newImage("shiftY", "32-bit black", 256, 254, 1);

// reserve memory for the result video
newImage("resultStack", "32-bit black", 256, 254, 36);


// shift some of the pixels in X
selectImage("shiftX");
makeOval(20, 98, 72, 68);
run("Add...", "value=25");
run("Select None");
run("Gaussian Blur...", "sigma=15");
run("Enhance Contrast", "saturated=0.35");

// init GPU
run("CLIJ Macro Extensions", "cl_device=");
Ext.CLIJ_push("blobs");
Ext.CLIJ_push("shiftX");
Ext.CLIJ_push("shiftY");
Ext.CLIJ_push("resultStack");

for (i = 0; i < 36; i++) {

	// change the shift from slice to slice
	Ext.CLIJ_affineTransform2D("shiftX", "rotatedShiftX", "center rotate=" + (i * 10) + " -center");
	
	// apply transform
	Ext.CLIJ_applyVectorField2D("blobs", "rotatedShiftX", "shiftY", "transformed");

	// put resulting 2D image in the right plane
	Ext.CLIJ_copySlice("transformed", "resultStack", i);
}


// get result back from GPU
Ext.CLIJ_pull("resultStack");
run("Invert LUT");
```
