## Supplementary Code 1: CLIJ usage examples for "CLIJ: GPU-accelerated image processing for everyone": autoThreshold.ijm

```
// CLIJ example macro: autoThreshold.ijm
//
// This macro shows how to apply an automatic 
// threshold method to an image in the GPU.
//
// Author: Robert Haase
// December 2018
// ---------------------------------------------


// Get test data
run("Blobs (25K)");
//open("C:/structure/data/blobs.gif");
getDimensions(width, height, channels, slices, frames);
input = getTitle();

mask = "Mask";

// Init GPU
run("CLIJ Macro Extensions", "cl_device=");
Ext.CLIJ_clear();

// push data to GPU
Ext.CLIJ_push(input);

// cleanup ImageJ
run("Close All");

// create a mask using a fixed threshold
Ext.CLIJ_automaticThreshold(input, mask, "Otsu");

// show result
Ext.CLIJ_pullBinary(mask);
```
