## Supplementary Code 1: CLIJ usage examples for "CLIJ: GPU-accelerated image processing for everyone": backgroundSubtraction.ijm

```
// CLIJ example macro: backgroundSubtraction.ijm
//
// This macro shows how background subtraction can be done in the GPU.
//
// Author: Robert Haase
// December 2018
// ---------------------------------------------

run ("Close All");

// Get test data
//open("C:/structure/data/t1-head.tif");
run("T1 Head (2.4M, 16-bits)");
input = getTitle();
background = "background";
background_subtracted = "background_subtracted";

// Init GPU
run("CLIJ Macro Extensions", "cl_device=");
Ext.CLIJ_clear();

// push images to GPU
Ext.CLIJ_push(input);

// CleanUp ImageJ
close();

// Blur in GPU
Ext.CLIJ_blur3D(input, background, 10, 10, 1);

// subtraction from original
Ext.CLIJ_addImagesWeighted(input, background, background_subtracted, 1, -1);

// Get results back from GPU
Ext.CLIJ_pull(background_subtracted);
Ext.CLIJ_pull(input);


// Cleanup by the end
Ext.CLIJ_clear();
```
