## Supplementary Code 1: CLIJ usage examples for "CLIJ: GPU-accelerated image processing for everyone": benchmarking.ijm

```
// CLIJ example macro: benchmarking.ijm
//
// This macro shows how to measure performance of GPU and CPU based ImageJ macro code.
//
// Author: Robert Haase
// December 2018
// ---------------------------------------------

// Get test data
run("T1 Head (2.4M, 16-bits)");
input = getTitle();
getDimensions(width, height, channels, slices, frames);

blurred = "Blurred";

// Init GPU
run("CLIJ Macro Extensions", "cl_device=");
Ext.CLIJ_clear();

// Local mean filter in CPU
for (i = 1; i <= 10; i++) {
	time = getTime();
	run("Mean 3D...", "x=3 y=3 z=3");
	print("CPU mean filter no " + i + " took " + (getTime() - time) + " msec");
}

// push images to GPU
time = getTime();
Ext.CLIJ_push(input);
print("Pushing one image to the GPU took " + (getTime() - time) + " msec");

// cleanup ImageJ
run("Close All");

// Local mean filter in GPU
for (i = 1; i <= 10; i++) {
	time = getTime();
	Ext.CLIJ_mean3DBox(input, blurred, 3, 3, 3);
	print("GPU mean filter no " + i + " took " + (getTime() - time) + " msec");
}

// Get results back from GPU
time = getTime();
Ext.CLIJ_pull(blurred);

print("Pulling one image from the GPU took " + (getTime() - time) + " msec");

// Cleanup GPU 
Ext.CLIJ_clear();
```
