## Supplementary Code 1: CLIJ usage examples for "CLIJ: GPU-accelerated image processing for everyone": bigImageTransfer.ijm

```
// CLIJ example macro: bigImageTransfer.ijm
//
// This macro shows how to process big images on the GPU
//
// Author: Robert Haase
// January 2019
// ---------------------------------------------

run("Close All");

// Get test dat - a 4 GB image of type 8-bit
width = 1024;
height = 1024;
slices = 4095;
newImage("original", "8-bit ramp", width, height, slices);

// init GPU
run("CLIJ Macro Extensions", "cl_device=");
Ext.CLIJ_clear();

// push images to GPU
Ext.CLIJ_push("original");

// cleanup imagej
run("Close All");

// add images

Ext.CLIJ_maximumZProjection("original", "max");

// show result
Ext.CLIJ_pull("max");

Ext.CLIJ_reportMemory();
Ext.CLIJ_clear();
```
