## Supplementary Code 1: CLIJ usage examples for "CLIJ: GPU-accelerated image processing for everyone": binaryProcessing.ijm

```
// CLIJ example macro: binaryProcessing.ijm
//
// This macro shows how to deal with binary images, e.g. thresholding, dilation, erosion, in the GPU.
//
// Author: Robert Haase
// December 2018
// ---------------------------------------------

run("Close All");


// Get test data
run("Blobs (25K)");
//open("C:/structure/data/blobs.gif");
getDimensions(width, height, channels, slices, frames);
input = getTitle();
threshold = 128;

mask = "mask";

temp = "temp";


// Init GPU
run("CLIJ Macro Extensions", "cl_device=");
Ext.CLIJ_clear();

// push data to GPU
Ext.CLIJ_push(input);

// cleanup ImageJ
run("Close All");

// create a mask using a fixed threshold
Ext.CLIJ_threshold(input, mask, threshold);

// binary opening: erosion + dilation, twice each
Ext.CLIJ_erodeBox(mask, temp);
Ext.CLIJ_erodeBox(temp, mask);

Ext.CLIJ_dilateBox(mask, temp);
Ext.CLIJ_dilateBox(temp, mask);


// show result
Ext.CLIJ_pullBinary(mask);
```
