## Supplementary Code 1: CLIJ usage examples for "CLIJ: GPU-accelerated image processing for everyone": bitdepthConversion.ijm

```
// CLIJ example macro: bitdepthConversion.ijm
//
// This macro shows how change the bit-depth of 
// images in the GPU.
//
// Author: Robert Haase
// March 2019
// ---------------------------------------------


// create an image with 32-bit
newImage("image32", "32-bit ramp", 20, 20, 1);

// init GPU
run("CLIJ Macro Extensions", "cl_device=");
Ext.CLIJ_clear();

// send image to GPU
Ext.CLIJ_push("image32");

// scale the image pixel intensities with a given factor
intensityScalingFactor = 255;
Ext.CLIJ_multiplyImageAndScalar("image32", "temp", intensityScalingFactor);

// convert it to 8 bit
Ext.CLIJ_convertUInt16("temp", "image16");

// get converted image back from GPU
Ext.CLIJ_pull("image16");
```
