## Supplementary Code 1: CLIJ usage examples for "CLIJ: GPU-accelerated image processing for everyone": blur.ijm

```
// CLIJ example macro: blur.ijm
//
// This macro shows how to blur an image in the GPU.
//
// Author: Robert Haase
// December 2018
// ---------------------------------------------
run("Close All");

// Get test data
run("T1 Head (2.4M, 16-bits)");
input = getTitle();
getDimensions(width, height, channels, slices, frames);

blurred = "Blurred";

// Init GPU
run("CLIJ Macro Extensions", "cl_device=");
Ext.CLIJ_clear();

// push images to GPU
Ext.CLIJ_push(input);

// cleanup ImageJ
run("Close All");

// Blur in GPU
Ext.CLIJ_blur3D(input, blurred, 5, 5, 1);

// Get results back from GPU
Ext.CLIJ_pull(blurred);

// Cleanup by the end
Ext.CLIJ_clear();
```
