## Supplementary Code 1: CLIJ usage examples for "CLIJ: GPU-accelerated image processing for everyone": clInfo.ijm

```
// CLIJ example macro: clInfo.ijm
//
// Outputs information about OpenCL devices
//
// Author: Robert Haase
// December 2018
// ---------------------------------------------


run("CLIJ Macro Extensions", "cl_device=");
Ext.CLIJ_clInfo();
```
