## Supplementary Code 1: CLIJ usage examples for "CLIJ: GPU-accelerated image processing for everyone": convert.ijm

```
// CLIJ example macro: convert,ijm
//
// This macro shows how to convert an image in the GPU.
//
// Author: Robert Haase
// December 2018
// ---------------------------------------------

run("Close All");

cl_device = "";

run("Blobs (25K)");
source = getTitle();

run("CLIJ Macro Extensions", "cl_device=" + cl_device);
Ext.CLIJ_clear();

Ext.CLIJ_push(source);

Ext.CLIJ_convertFloat(source, "float");
Ext.CLIJ_pull("float");

Ext.CLIJ_convertUInt8("float", "uint8");
Ext.CLIJ_pull("uint8");

Ext.CLIJ_convertUInt16(source, "uint16");
Ext.CLIJ_pull("uint16");
```
