## Supplementary Code 1: CLIJ usage examples for "CLIJ: GPU-accelerated image processing for everyone": create_object_outlines.ijm

```
// CLIJ example macro: create_object_outlines.ijm
//
// This macro shows how to deal with binary images, e.g. thresholding, dilation, erosion, in the GPU.
//
// This macro was generate using the ImageJ macro recorder. 
//
// Author: Robert Haase
// December 2018
// ---------------------------------------------

run("Blobs (25K)");

//run("Close All");
//open("C:/structure/data/blobs.gif");
run("CLIJ Macro Extensions", "cl_device=");

Ext.CLIJ_push("blobs.gif");
Ext.CLIJ_mean2DBox("blobs.gif", "CLIJ_mean2DBox_destination_blobs.gif", 2.0, 2.0);
Ext.CLIJ_threshold("CLIJ_mean2DBox_destination_blobs.gif", "CLIJ_threshold_destination_CLIJ_mean2DBox_destination_blobs.gif", 127.0);
Ext.CLIJ_erodeBox("CLIJ_threshold_destination_CLIJ_mean2DBox_destination_blobs.gif", "CLIJ_erodeBox_destination_CLIJ_threshold_destination_CLIJ_mean2DBox_destination_blobs.gif");
Ext.CLIJ_binaryXOr("CLIJ_threshold_destination_CLIJ_mean2DBox_destination_blobs.gif", "CLIJ_erodeBox_destination_CLIJ_threshold_destination_CLIJ_mean2DBox_destination_blobs.gif", "CLIJ_binaryXOr_destination_CLIJ_threshold_destination_CLIJ_mean2DBox_destination_blobs.gif");
Ext.CLIJ_pull("CLIJ_binaryXOr_destination_CLIJ_threshold_destination_CLIJ_mean2DBox_destination_blobs.gif");
```
