## Supplementary Code 1: CLIJ usage examples for "CLIJ: GPU-accelerated image processing for everyone": help.ijm

```
// CLIJ example macro: help.ijm
//
// This macro shows how to get help on CLIJ methods
//
// Author: Robert Haase
// December 2018
// ---------------------------------------------


run("CLIJ Macro Extensions", "cl_device=");
Ext.CLIJ_help("");
```
