## Supplementary Code 1: CLIJ usage examples for "CLIJ: GPU-accelerated image processing for everyone": localMaximum.ijm

```
// localMaximum.ijm
//
// Demonstrates differences between local "maximum" filters in IJ and CLIJ. 
//
// Author: haesleinhuepf
// Jan 2019

run("Close All");

for (radius = 1; radius < 4; radius++) { 

	newImage("original", "8-bit black", 9, 9, 9);
	Stack.setSlice(5);
	makeRectangle(4, 4, 1, 1);
	run("Add...", "value=255 slice");
	run("Select None");
	
	selectWindow("original");
	run("Duplicate...", "title=orignal1slice");
	
	
	run("CLIJ Macro Extensions", "cl_device=");
	Ext.CLIJ_clear();
	
	Ext.CLIJ_push("original");
	Ext.CLIJ_push("orignal1slice");
	
	
	
	Ext.CLIJ_maximum3DSphere("original", "maximum3DSphereCLIJ_" + radius, radius, radius, radius);
	Ext.CLIJ_pull("maximum3DSphereCLIJ_" + radius);
	zoom("maximum3DSphereCLIJ_" + radius);
	
	Ext.CLIJ_maximum2DSphere("orignal1slice", "maximum2DSphereCLIJ_" + radius, radius, radius);
	Ext.CLIJ_pull("maximum2DSphereCLIJ_" + radius);
	zoom("maximum2DSphereCLIJ_" + radius);
	
	selectWindow("original");
	run("Duplicate...", "title=maximum3DIJ_" + radius + " duplicate");
	run("Maximum 3D...", "x=" + radius + " y=" + radius + " z=" + radius);
	zoom("maximum3DIJ_" + radius);
	
	selectWindow("orignal1slice");
	run("Duplicate...", "title=maximum2DIJ_" + radius);
	run("Maximum...", "radius=" + radius);
	zoom("maximum2DIJ_" + radius);
}

function zoom(title) {
	selectWindow(title);
	for (i = 0; i < 10; i++) {
		run("In [+]");
	}
}
```
