## Supplementary Code 1: CLIJ usage examples for "CLIJ: GPU-accelerated image processing for everyone": maximumProjection.ijm

```
// CLIJ example macro: backgroundSubtraction.ijm
//
// This macro shows how maximum projection can be done in the GPU.
//
// Author: Robert Haase
// December 2018
// ---------------------------------------------

run ("Close All");

// Get test data
//open("C:/structure/data/t1-head.tif");
run("T1 Head (2.4M, 16-bits)");
input = getTitle();
background = "background";
background_subtracted = "background_subtracted";
maximum_projected = "maximum_projected";

// Init GPU
run("CLIJ Macro Extensions", "cl_device=");
Ext.CLIJ_clear();

// push images to GPU
Ext.CLIJ_push(input);

// CleanUp ImageJ
close();

// maximum projection
Ext.CLIJ_maximumZProjection(input, maximum_projected);

// Get results back from GPU
Ext.CLIJ_pull(maximum_projected);

// Cleanup by the end
Ext.CLIJ_clear();
```
