## Supplementary Code 1: CLIJ usage examples for "CLIJ: GPU-accelerated image processing for everyone": mean2d.ijm

```
// CLIJ example macro: mean.ijm
//
// This macro shows how the mean average filter works in the GPU.
//
// Author: Robert Haase
// December 2018
// ---------------------------------------------
run("Close All");

// Get test data
run("T1 Head (2.4M, 16-bits)");
run("Duplicate...", " ");
rename("Mean CPU")
input = getTitle();
getDimensions(width, height, channels, slices, frames);

blurred = "Mean GPU";

// Init GPU
run("CLIJ Macro Extensions", "cl_device=");
Ext.CLIJ_clear();

// push images to GPU
Ext.CLIJ_push(input);

// Local mean filter in CPU
selectWindow(input);
run("Mean...", "radius=3");

// Local mean filter in GPU
Ext.CLIJ_mean2DBox(input, blurred, 3, 3);

// Get results back from GPU
Ext.CLIJ_pull(blurred);

// Cleanup GPU 
Ext.CLIJ_clear();


imageCalculator("Subtract create 32-bit stack", "Mean CPU","Mean GPU");
selectWindow("Result of Mean CPU");
rename("Difference between CPU and GPU");
//setSlice(60);
run("Enhance Contrast", "saturated=0.35");
```
