## Supplementary Code 1: CLIJ usage examples for "CLIJ: GPU-accelerated image processing for everyone": mean_detailed_comparison_IJ_CLIJ.ijm

```
// CLIJ example macro: mean_detailed_comparison_IJ_CLIJ.ijm
//
// This macro shows differences between ImageJ and CLIJ when executing a mean filter.
// Starting from an image (2D or 3D) where a single pixel is set to 255, mean filters are applied.
// While we observe differences between 2D and 3D in ImageJ. CLIJ is more consistent.
//
// Author: Robert Haase
// June 2019
// ---------------------------------------------

run("Close All");

radius = 1;


// -----------------------------------------------------------------

// create 2D example image
input = "example2d";
newImage(input, "8-bit black", 10, 10, 1);
makeRectangle(4, 5, 1, 1);
run("Add...", "value=255");
run("Select None");

// visualise input image
zoom(10);
setWindowPosition(0, 0);
setMinAndMax(0, 1);
makeOverlay();

// init GPU
run("CLIJ Macro Extensions", "cl_device=");

// do operation on GPU
outputGPU = "mean2d_GPU";
Ext.CLIJ_push(input);
Ext.CLIJ_mean2DSphere(input, outputGPU, radius, radius);
Ext.CLIJ_pull(outputGPU);

// visualise result
zoom(10);
setWindowPosition(1, 0);
setMinAndMax(0, 1);

makeOverlay();

// do operation on CPU
selectWindow(input);
run("Duplicate...", "title=mean2D_CPU");
run("Mean...", "radius=" + radius);

// visualise result
zoom(10);
setWindowPosition(2, 0);
setMinAndMax(0, 1);

makeOverlay();

// -----------------------------------------------------------------

// create 3D example image
input = "example3d";
newImage(input, "8-bit black", 10, 10, 10);
setSlice(4);
makeRectangle(4, 5, 1, 1);
run("Add...", "value=255");
run("Select None");

// visualise input image
zoom(10);
setWindowPosition(0, 1);
setMinAndMax(0, 1);
makeOverlay();

// init GPU
run("CLIJ Macro Extensions", "cl_device=");

// do operation on GPU
outputGPU = "mean3d_GPU";
Ext.CLIJ_push(input);
Ext.CLIJ_mean3DSphere(input, outputGPU, radius, radius, radius);
Ext.CLIJ_pull(outputGPU);

// visualise result
setSlice(4);
zoom(10);
setWindowPosition(1, 1);
setMinAndMax(0, 1);
makeOverlay();

// do operation on CPU
selectWindow(input);
run("Duplicate...", "title=mean3D_CPU duplicate");
run("Mean 3D...", "x=" + radius + " y=" + radius + " z=" + radius);

// visualise result
zoom(10);
setSlice(4);
setWindowPosition(2, 1);
setMinAndMax(0, 1);

makeOverlay();

// -----------------------------------------------------------------

function zoom(count) {
	for (i = 0; i < count; i++) {	
		run("In [+]");
	}	
}

function setWindowPosition(x, y) {
	setLocation(x * 330, y * 370);
}

function makeOverlay() {
	for (y = 1; y <= 9; y += 2) {
		makeRectangle(0, y, 10, 1);
		Roi.setStrokeColor("gray");
		run("Add Selection...");
	}
	for (x = 1; x <= 9; x += 2) {
		makeRectangle(x, 0, 1, 10);
		Roi.setStrokeColor("gray");
		run("Add Selection...");	
	}
	run("Select None");
}
```
