## Supplementary Code 1: CLIJ usage examples for "CLIJ: GPU-accelerated image processing for everyone": memory_reuse_versus_reallocation.ijm

```
// CLIJ example macro: memory_reuse_versus_reallocation.ijm
//
// This exmple macro shows that processing speed can be gained
// by reusing image memory in the GPU instead of allocating 
// memory during every iteration
//
// Author: Robert Haase
// July 2019
// ---------------------------------------------

run("Close All");

width = 1024;
height = 1024;
slices = 100;

// Get test data
newImage("original", "16-bit ramp", width, height, slices);

// init GPU
run("CLIJ Macro Extensions", "cl_device=");
Ext.CLIJ_clear();

// push images to GPU
Ext.CLIJ_push("original");

// cleanup imagej
run("Close All");

for (j = 0; j < 10; j++) {
	// copy the image 10 times wihile reusing target memory
	time = getTime();
	for (i = 0; i < 10; i++) {
		Ext.CLIJ_copy("original", "copy");
	}
	IJ.log("Copying with memory reusing took " + (getTime() - time));
	
	// copy the image 10 times while releasing and reallocating target memory.
	time = getTime();
	for (i = 0; i < 10; i++) {
		Ext.CLIJ_copy("original", "copy");
		Ext.CLIJ_release("copy");
	}
	IJ.log("Copying with memory reallocation took " + (getTime() - time));
}
```
