## Supplementary Code 1: CLIJ usage examples for "CLIJ: GPU-accelerated image processing for everyone": motionCorrection_compare_stackreg.ijm

```
// CLIJ example macro: motionCorrection_compare_stackreg.ijm
//
// This macro compares performance of motion correction
// on the GPU with StackReg. You need to install the
// BIG-EPFL update site in order to test it.
//
// Author: Robert Haase
// June 2019
// ---------------------------------------------
run("Close All");

// define move to correct
file = "https://github.com/clij/clij-docs/raw/master/src/main/resources/motion_correction_Drosophila_DSmanila1.tif";
//file = "C:/structure/code/clij-docs/src/main/resources/motion_correction_Drosophila_DSmanila1.tif";

// define identifiers for intermediate results in the GPU
inputStack = "inputStack";
slice = "slice";
shifted = "shifted";
binary = "binary";

// define a threshold to differentiate object and background
threshold = 50;

// open image
open(file);
rename(inputStack);
run("32-bit");

// collect info about it
getDimensions(width, height, channels, slices, frames);


// Init GPU
time = getTime();
run("CLIJ Macro Extensions", "cl_device=");
Ext.CLIJ_clear();

Ext.CLIJ_push(inputStack);

formerX = 0;
formerY = 0;

// process all slices; only the first stays where it is
for (z = 0; z  < slices; z++) {
	Ext.CLIJ_copySlice(inputStack, slice, z);

	// determine center of mass
	Ext.CLIJ_threshold(slice, binary, threshold);
	Ext.CLIJ_centerOfMass(binary);
	x = getResult("MassX", nResults() - 1);
	y = getResult("MassY", nResults() - 1);

	if (z > 0) {
		
		// determine shift
		deltaX = x - formerX;
		deltaY = y - formerY;

		// apply translation transformation
		Ext.CLIJ_affineTransform2D(slice, shifted, "translatex=" + deltaX + " translatey=" + deltaY);

		// copy result back
		Ext.CLIJ_copySlice(shifted, inputStack, z);
	
	} else {
		formerX = x;
		formerY = y;
	}
}

// show result
Ext.CLIJ_pull(inputStack);
rename("CLIJ motion corrected");
IJ.log("Motion correction with CLIJ took " + (getTime() - time) + " msec");

// open image again and register with stackreg
open(file);
time = getTime();
rename("StackReg motion corrected");
run("32-bit");
run("StackReg", "transformation=Translation");

IJ.log("Motion correction with StackReg took " + (getTime() - time) + " msec");

open(file);
rename("Original");
run("32-bit");

imageCalculator("Subtract create 32-bit stack", "CLIJ motion corrected", "StackReg motion corrected");
rename("Difference");

images = newArray("Original", "CLIJ motion corrected", "StackReg motion corrected", "Difference");
for (i = 0; i < 4; i++) {
	selectWindow(images[i]);
	setLocation(50 + 300 * i, 50);
	run("Out [-]");
	run("Out [-]");
}
```
