## Supplementary Code 1: CLIJ usage examples for "CLIJ: GPU-accelerated image processing for everyone": orthogonalMaximumProjections.ijm

```
// CLIJ example macro: orthogonalMaximumProjections.ijm
//
// This macro shows how maximum-X, -Y and -Z projections can be created using the GPU.
//
// Author: Robert Haase
// December 2018
// ---------------------------------------------
run("Close All");

// Get test data
run("T1 Head (2.4M, 16-bits)");
//open("C:/structure/data/t1-head.tif");
getDimensions(width, height, channels, slices, frames);
input = getTitle();
downScalingFactorInXY = 0.666; // because the image has slice distance 1.5
downScalingFactorInZ = 1;

downscaled = "Downscaled";

maximumProjectionX = "Maximum projection in X";
maximumProjectionY = "Maximum projection in Y";
maximumProjectionZ = "Maximum projection in Z";

// Init GPU
run("CLIJ Macro Extensions", "cl_device=");
Ext.CLIJ_clear();

// push data to GPU
Ext.CLIJ_push(input);

// cleanup ImageJ
run("Close All");

// process
Ext.CLIJ_downsample3D(input, downscaled, downScalingFactorInXY, downScalingFactorInXY, downScalingFactorInZ);

Ext.CLIJ_maximumXYZProjection(downscaled, maximumProjectionX, 2, 1, 0);
Ext.CLIJ_maximumXYZProjection(downscaled, maximumProjectionY, 2, 0, 1);
Ext.CLIJ_maximumXYZProjection(downscaled, maximumProjectionZ, 0, 1, 2);

// show results
Ext.CLIJ_pull(maximumProjectionX);
Ext.CLIJ_pull(maximumProjectionY);
Ext.CLIJ_pull(maximumProjectionZ);
```
