## Supplementary Code 1: CLIJ usage examples for "CLIJ: GPU-accelerated image processing for everyone": project3D.ijm

```
// CLIJ example macro: project3D.ijm
//
// This macro shows how a 3D projection can be done in the GPU.
//
// Author: Robert Haase
// January 2019
// ---------------------------------------------


run("Close All");

// Get test data
//open("C:/structure/data/t1-head.tif");
run("T1 Head (2.4M, 16-bits)");

angle_step = 10;

run("32-bit");
rename("original");

time  = getTime();
run("3D Project...", "projection=[Brightest Point] axis=Y-Axis slice=1 initial=0 total=360 rotation=" + angle_step + " lower=1 upper=255 opacity=0 surface=100 interior=50 interpolate");
close();
IJ.log("CPU 3D projection took " + (getTime()-time) + " msec");

getDimensions(width, height, channels, depth, frames);


// init GPU
run("CLIJ Macro Extensions", "cl_device=");
Ext.CLIJ_clear();

time = getTime();

// push images to GPU
Ext.CLIJ_push("original");
// reserve the right amount of memory for the result image
Ext.CLIJ_create3D("target", width, height, 360 / angle_step, 32);

// reserve a bit more pixels in Z for translated and rotated image because we
// need space for the shoulders if we rotated the patient around the Y-axis
Ext.CLIJ_create3D("rotated", width, height, depth * 2, 32);
Ext.CLIJ_create3D("translated", width, height, depth * 2, 32);

// cleanup imagej
run("Close All");

// we need to translate the stack in Z to get some space for the shoulders 
// when we rotate the head around the y-axis 
Ext.CLIJ_translate3D("original", "translated", 0, 0, depth / 2);
	
count = 0;
for (angle = 0; angle < 360; angle += angle_step) {
	Ext.CLIJ_rotate3D("translated", "rotated", 0, angle, 0.0, true);
	Ext.CLIJ_maximumZProjection("rotated", "maxProjected");

	// put the maximum projection in the right place in the result stack
	Ext.CLIJ_copySlice("maxProjected", "target", count);
	
	count++;
}

// show result
Ext.CLIJ_pull("target");

IJ.log("GPU 3D projection took " + (getTime()-time) + " msec");
```
