## Supplementary Code 1: CLIJ usage examples for "CLIJ: GPU-accelerated image processing for everyone": push.ijm

```
// CLIJ example macro: push.ijm
//
// This macro shows how an image is pushed to the GPU.
//
// Author: Robert Haase
// December 2018
// ---------------------------------------------


// Get test data
// run("T1 Head (2.4M, 16-bits)");
newImage("Untitled", "16-bit black", 256, 256, 128);
rename("Mean CPU")
input = getTitle();
getDimensions(width, height, channels, slices, frames);

// Init GPU
run("CLIJ Macro Extensions", "cl_device=gfx902");
Ext.CLIJ_clear();

// push images to GPU
Ext.CLIJ_push(input);

Ext.CLIJ_reportMemory();
```
