## Supplementary Code 1: CLIJ usage examples for "CLIJ: GPU-accelerated image processing for everyone": reslicing.ijm

```
// CLIJ example macro: reslicing.ijm
//
// This macro shows how stacks can be resliced in the GPU
//
// Author: Robert Haase
// December 2018
// ---------------------------------------------
run("Close All");

// Get test data
run("T1 Head (2.4M, 16-bits)");
//open("C:/structure/data/t1-head.tif");
getDimensions(width, height, channels, slices, frames);
input = getTitle();

resliced = "Resliced left";

// Init GPU
run("CLIJ Macro Extensions", "cl_device=");
Ext.CLIJ_clear();

// push data to GPU
Ext.CLIJ_push(input);

// reslice
Ext.CLIJ_resliceLeft(input, resliced);

// show results
Ext.CLIJ_pull(resliced);
```
