## Supplementary Code 1: CLIJ usage examples for "CLIJ: GPU-accelerated image processing for everyone": rotate_comparison_IJ_CLIJ.ijm

```
// CLIJ example macro: rotate_comparison_IJ_CLIJ.ijm
//
// This macro shows how stacks can be rotated in the GPU
// and how different results are between CLIJ and ImageJ.
//
// Author: Robert Haase
// July 2019
// ---------------------------------------------
run("Close All");


// Get test data
run("Blobs (25K)");
run("32-bit");

getDimensions(width, height, channels, slices, frames);
input = getTitle();

rotated = "Rotated";

// Init GPU
run("CLIJ Macro Extensions", "cl_device=1070");
Ext.CLIJ_clear();

// push data to GPU
Ext.CLIJ_push(input);

// rotate on CPU
run("Rotate... ", "angle=45 grid=1 interpolation=Bilinear");

// rotate on GPU
Ext.CLIJ_affineTransform2D(input, rotated, "center rotate=45 -center");

// show results
Ext.CLIJ_pull(rotated);

// calculate difference image between CPU and GPU
imageCalculator("Subtract create 32-bit", input, rotated);
```
