## Supplementary Code 1: CLIJ usage examples for "CLIJ: GPU-accelerated image processing for everyone": rotateFree.ijm

```
// CLIJ example macro: rotateFree.ijm
//
// This macro shows how to rotate an image in the GPU.
//
// Author: Robert Haase
// January 2019
// ---------------------------------------------


//run("Close All");

// Get test data
run("Blobs (25K)");
//open("C:/structure/data/blobs.gif");


angle_step = 10;

run("32-bit");
rename("original");

getDimensions(width, height, channels, depth, frames);


// init GPU
run("CLIJ Macro Extensions", "cl_device=");
Ext.CLIJ_clear();

// push images to GPU
Ext.CLIJ_push("original");
// reserve the right amount of memory for the result image
Ext.CLIJ_create3D("target",  width, height, 360 / angle_step, 32);

// cleanup imagej
run("Close All");
	
count = 0;
for (angle = 0; angle < 360; angle += angle_step) {
	Ext.CLIJ_rotate2D("original", "rotated", angle, true);

	// put the rotated image in the right place in the result stack
	Ext.CLIJ_copySlice("rotated", "target", count);
	
	count++;
}

// show result
Ext.CLIJ_pull("target");
run("Invert LUT");
```
