## Supplementary Code 1: CLIJ usage examples for "CLIJ: GPU-accelerated image processing for everyone": scaleFree.ijm

```
// CLIJ example macro: project3D.ijm
//
// This macro shows how to rotate an image in the GPU.
//
// Author: Robert Haase
// January 2019
// ---------------------------------------------


run("Close All");

// Get test data
run("Blobs (25K)");
//open("C:/structure/data/blobs.gif");


zoom_step = 0.03;

run("32-bit");
rename("original");

getDimensions(width, height, channels, depth, frames);

// init GPU
run("CLIJ Macro Extensions", "cl_device=");
Ext.CLIJ_clear();

// push images to GPU
Ext.CLIJ_push("original");
// reserve the right amount of memory for the result image
Ext.CLIJ_create3D("target",  width, height, 1.0 / zoom_step, 32);

// cleanup imagej
run("Close All");
	
count = 0;
for (zoom = 1; zoom > 0; zoom -= zoom_step) {
	Ext.CLIJ_scale("original", "zoomed", zoom, true);

	// put the zoomed image in the right place in the result stack
	Ext.CLIJ_copySlice("zoomed", "target", count);
	
	count++;
}

// show result
Ext.CLIJ_pull("target");
run("Invert LUT");
```
