## Supplementary Code 1: CLIJ usage examples for "CLIJ: GPU-accelerated image processing for everyone": thresholding.ijm

```
// CLIJ example macro: thresholding,ijm
//
// This macro shows how to apply a threshold to an image in the GPU.
//
// Author: Robert Haase
// December 2018
// ---------------------------------------------


// Get test data
run("Blobs (25K)");
//open("C:/structure/data/blobs.gif");
getDimensions(width, height, channels, slices, frames);
input = getTitle();
threshold = 128;

// create memory for mask image
newImage("Untitled", "8-bit black", width, height, slices);
rename("Mask");
mask = getTitle();

// Init GPU
run("CLIJ Macro Extensions", "cl_device=");
Ext.CLIJ_clear();

// push data to GPU
Ext.CLIJ_push(input);
Ext.CLIJ_push(mask);

// cleanup ImageJ
run("Close All");

// create a mask using a fixed threshold
Ext.CLIJ_threshold(input, mask, threshold);

// show result
Ext.CLIJ_pullBinary(mask);
```
