## Supplementary Code 1: CLIJ usage examples for "CLIJ: GPU-accelerated image processing for everyone": topHat.ijm

```
// CLIJ example macro: topHat.ijm
//
// This macro shows how a top-hat filter can be applied in the GPU.
//
// Author: Robert Haase
// December 2018
// ---------------------------------------------

radiusXY = 10;
radiusZ = 0;

run ("Close All");

// Get test data
//open("C:/structure/data/t1-head.tif");
run("T1 Head (2.4M, 16-bits)");

input = getTitle();
median = "median";
temp1 = "temp1";
temp2 = "temp2";
output = "output";

// Init GPU
run("CLIJ Macro Extensions", "cl_device=");
Ext.CLIJ_clear();

// push images to GPU
Ext.CLIJ_push(input);

// CleanUp ImageJ
close();


Ext.CLIJ_medianSliceBySliceSphere(input, median, 1, 1);
Ext.CLIJ_minimum3DBox(median, temp1, radiusXY, radiusXY, radiusZ);
Ext.CLIJ_maximum3DBox(temp1, temp2, radiusXY, radiusXY, radiusZ);
Ext.CLIJ_subtractImages(median, temp2, output);

Ext.CLIJ_pull(input);
Ext.CLIJ_pull(output);

// Cleanup by the end
Ext.CLIJ_clear();
```
