## Supplementary Code 1: CLIJ usage examples for "CLIJ: GPU-accelerated image processing for everyone": translate.ijm

```
// CLIJ example macro: translate.ijm
//
// This macro shows how to translate an image in the GPU
// and make a video of an openign curtain.
//
// Author: Robert Haase
// July 2019
// ---------------------------------------------


//run("Close All");

// Get test data
run("Blobs (25K)");
//open("C:/structure/data/blobs.gif");


step = 10;

run("32-bit");
rename("original");

getDimensions(width, height, channels, depth, frames);

// init GPU
run("CLIJ Macro Extensions", "cl_device=");
Ext.CLIJ_clear();

// push images to GPU
Ext.CLIJ_push("original");
// reserve the right amount of memory for the result image
Ext.CLIJ_create3D("target",  width, height, 200 / step, 32);

// cleanup imagej
run("Close All");
	
count = 0;
for (displacement = 0; displacement < 200; displacement += step) {
	Ext.CLIJ_translate2D("original", "translated", displacement, 0);

	// put the translated image in the right place in the result stack
	Ext.CLIJ_copySlice("translated", "target", count);
	
	count++;
}

// show result
Ext.CLIJ_pull("target");
run("Invert LUT");
```
