## Supplementary Code 1: CLIJ usage examples for "CLIJ: GPU-accelerated image processing for everyone": turn_stack.ijm

```
// CLIJ example macro: turn_stack.ijm
//
// This macro shows how to wrangle voxels in 3D in the GPU.
//
// This marco was created using the macro recorder
//
// Author: Robert Haase
// December 2018
// ---------------------------------------------

run("Close All");

run("T1 Head (2.4M, 16-bits)");
//open("C:/structure/data/t1-head.tif");
run("CLIJ Macro Extensions", "cl_device=");
Ext.CLIJ_push("t1-head.tif");
Ext.CLIJ_resliceLeft("t1-head.tif", "CLIJ_resliceLeft_destination_t1-head.tif");
Ext.CLIJ_rotateRight("CLIJ_resliceLeft_destination_t1-head.tif", "CLIJ_rotateRight_destination_CLIJ_resliceLeft_destination_t1-head.tif");
Ext.CLIJ_pull("CLIJ_rotateRight_destination_CLIJ_resliceLeft_destination_t1-head.tif");
Ext.CLIJ_reportMemory();
```
