## Supplementary Code 1: CLIJ usage examples for "CLIJ: GPU-accelerated image processing for everyone": warpCat.ijm

```
// This script demonstrates how to apply a vector field
// to an image in order to transform it non-rigidly
//
// It uses a picture showing Pixel. 
// Yes, the cats name is Pixel.
//
// Make sure to activate the "CLIJ" update site before 
// running this macro in Fiji.
//
// Author: Robert Haase,
// May 2019
//


run("Close All");

// get test image
open("https://github.com/clij/clij-docs/raw/master/src/main/resources/pixel_cat.tif");
run("32-bit");
rename("cat");

getDimensions(width, height, channels, slices, frames)

// create two images describing local shift
newImage("shiftX", "32-bit black", width, height, 1);
newImage("shiftY", "32-bit black", width, height, 1);

// reserve memory for the result video
newImage("resultStack", "32-bit black", width, height, 36);


// shift some of the pixels in X
selectImage("shiftX");
makeOval(150, 150, 200, 200);
run("Add...", "value=50");
run("Select None");
run("Gaussian Blur...", "sigma=50");
run("Enhance Contrast", "saturated=0.35");

// init GPU
run("CLIJ Macro Extensions", "cl_device=");
Ext.CLIJ_push("cat");
Ext.CLIJ_push("shiftX");
Ext.CLIJ_copy("shiftX", "shiftY");
Ext.CLIJ_push("resultStack");

// clean up imagej
run("Close All");

for (i = 0; i < 36; i++) {

	// change the shift from slice to slice
	Ext.CLIJ_affineTransform2D("shiftX", "rotatedShiftX", "center rotate=" + (i * 10) + " -center");
	Ext.CLIJ_affineTransform2D("shiftY", "rotatedShiftY", "center rotate=" + (i * 10) + " -center");
	
	// apply transform
	Ext.CLIJ_applyVectorField2D("cat", "rotatedShiftX", "rotatedShiftY", "transformed");

	// put resulting 2D image in the right plane
	Ext.CLIJ_copySlice("transformed", "resultStack", i);
}


// get result back from GPU
Ext.CLIJ_pull("resultStack");

Ext.CLIJ_clear();
```
