## Supplementary figures and images for "CLIJ: GPU-accelerated image processing for everyone"

### Supplementary Figure 1: Processing time versus image size for 2D images

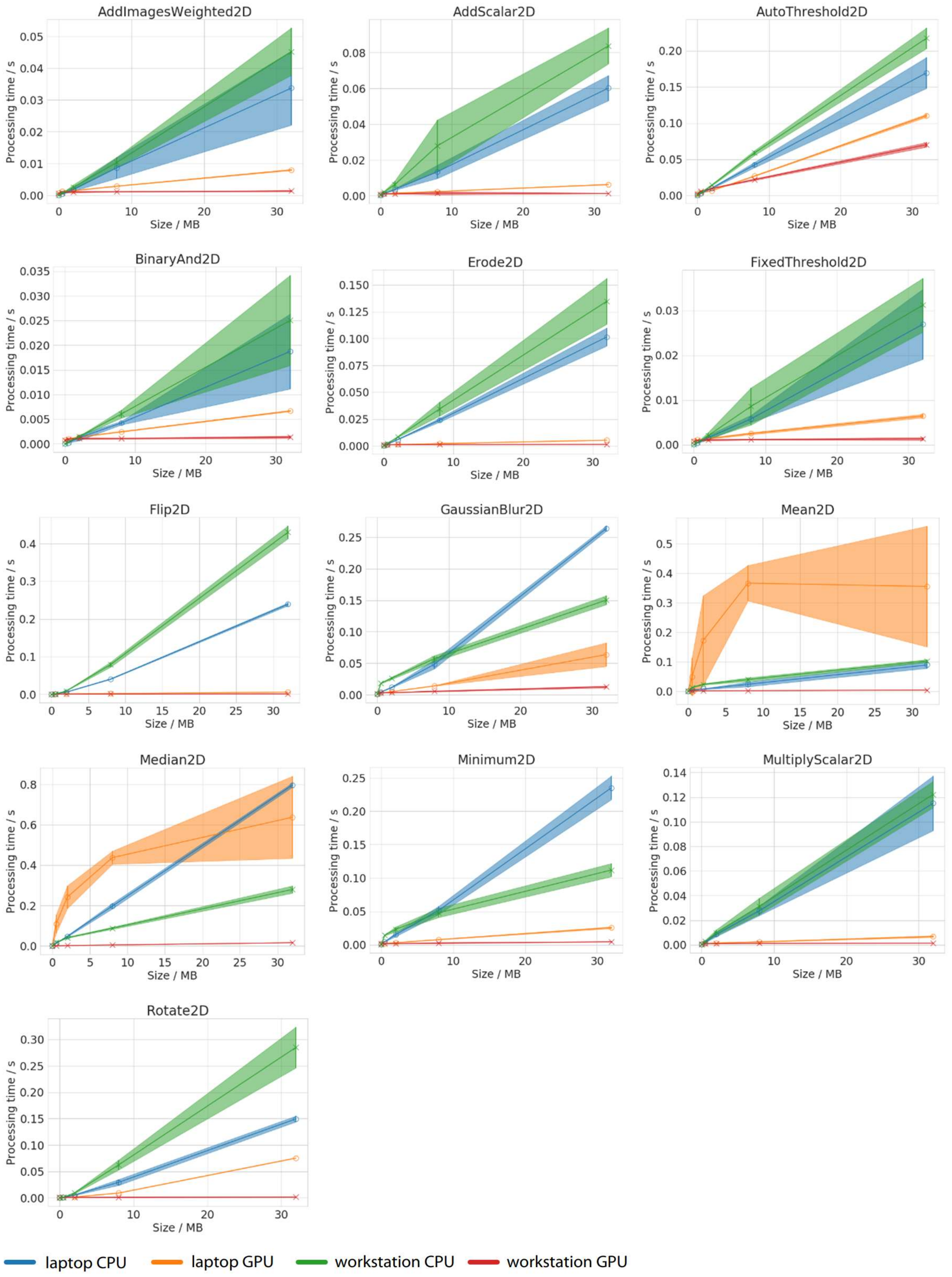


Supplemental Figure 1: Processing time of benchmarked 2D operations versus image size.

### Supplementary Figure 2: Processing time versus image size for 3D images

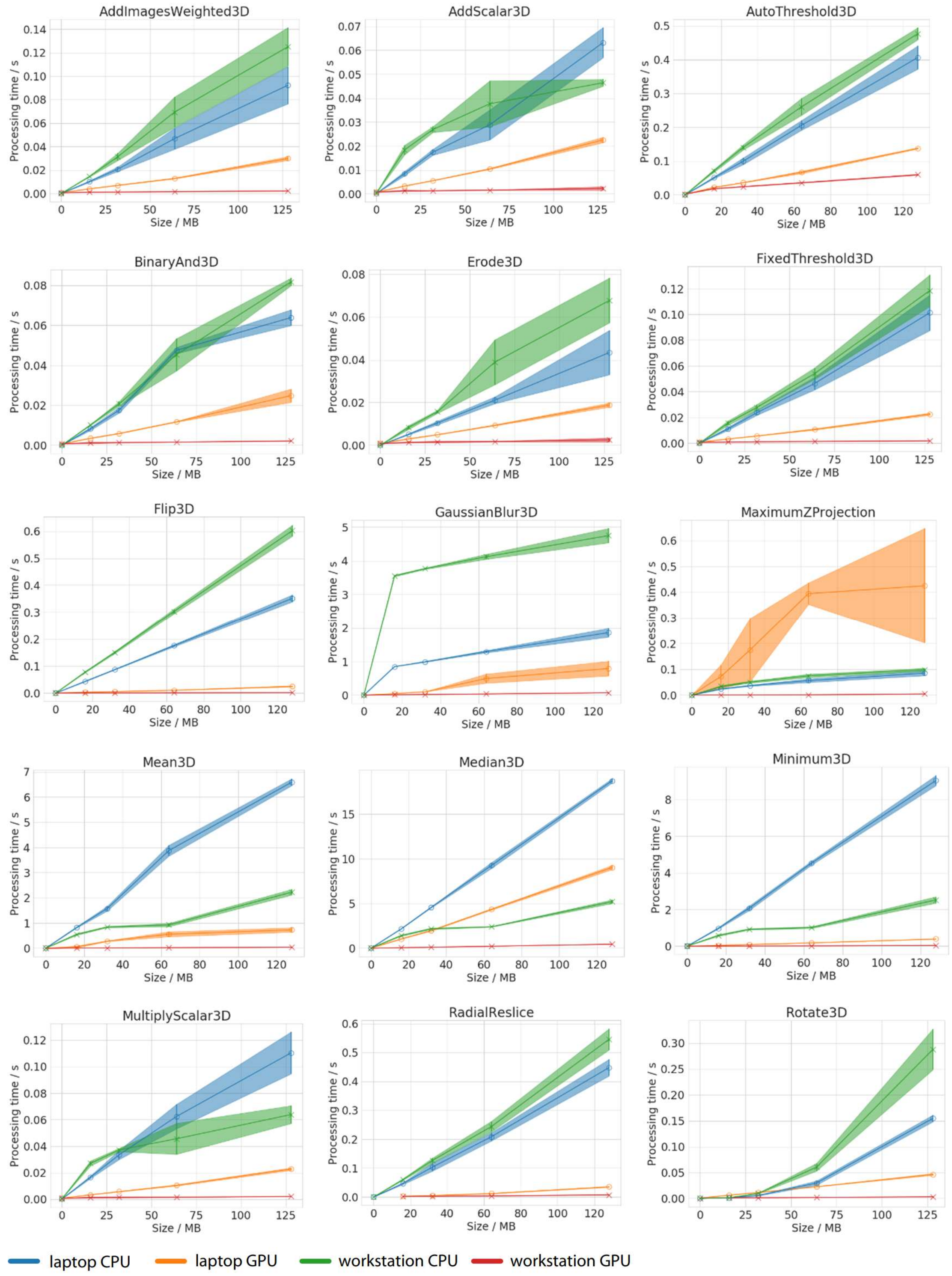


Supplemental Figure 2: Processing time of benchmarked 3D operations versus image size.
