## Supplementary Figure 3: Processing time versus parameters for "CLIJ: GPU-accelerated image processing for everyone"

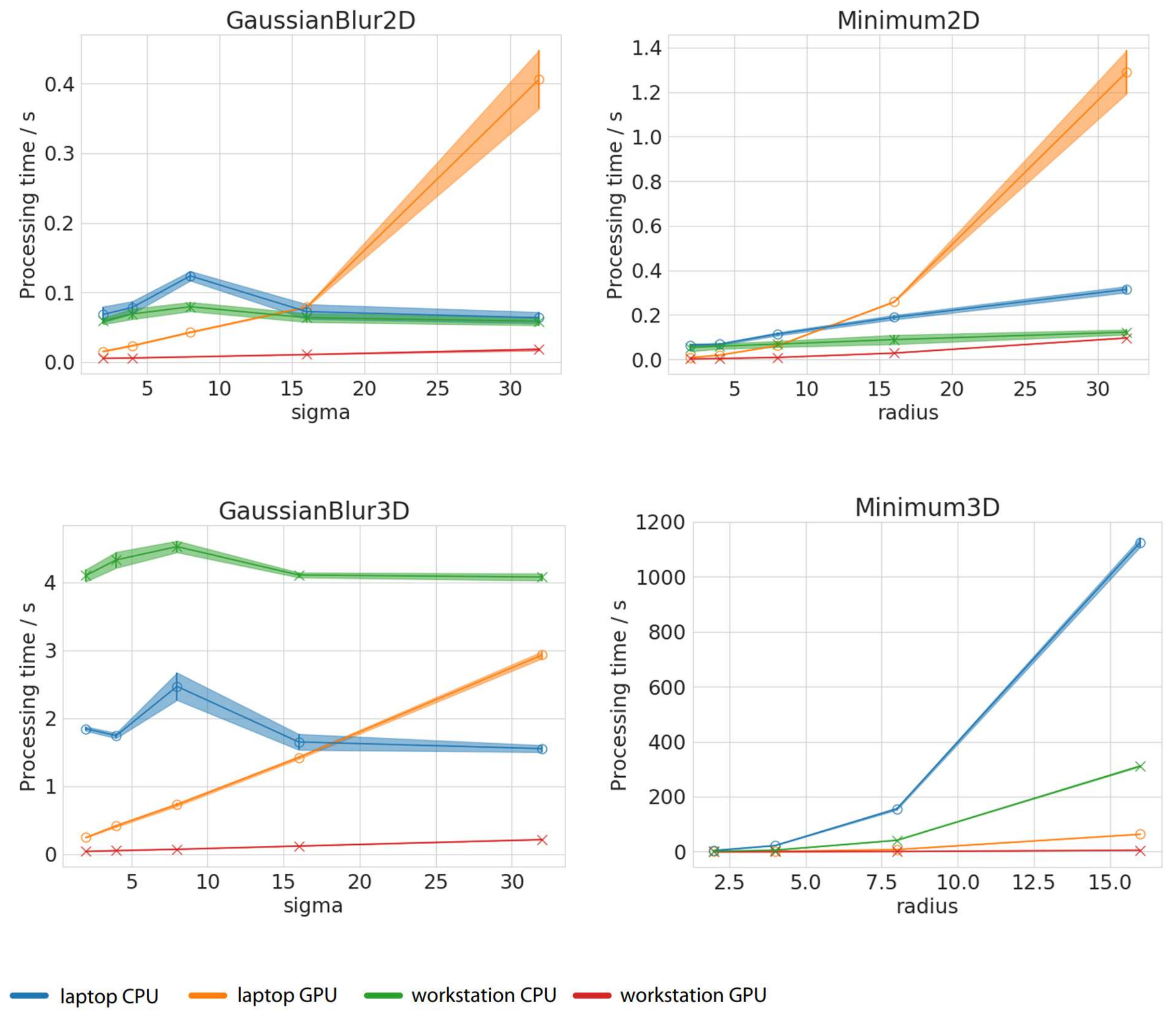


Supplemental Figure 3: Processing time of Gaussian Blur and Minimum filters in 2D and 3D applied to 32 MB (2D) and 64 MB (3D) large images.
