## Supplementary Figure 4: Benchmarked workflow for "CLIJ: GPU-accelerated image processing for everyone"

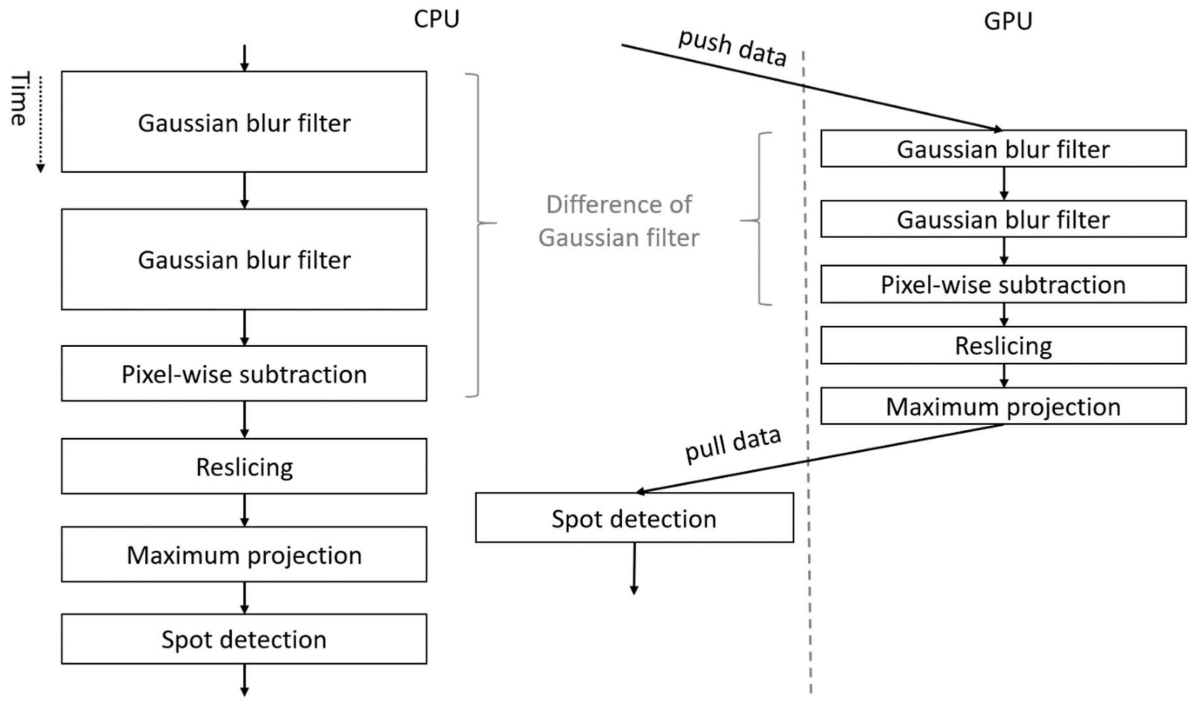


**Supplemental Figure 4:** Sequence diagram visualising that the GPU-accelerated workflow ends earlier even though data has to be pushed to GPU memory and pulled back. Difference-of-Gaussian filtering and a cylindrical maximum projection pre-process the image stack before spot detection is used to estimate nuclei count on the surface projection of a Drosophila embryo.
