## Supplementary Figure 5: Differences between ImageJ and CLIJ workflows for "CLIJ: GPU-accelerated image processing for everyone"

a)

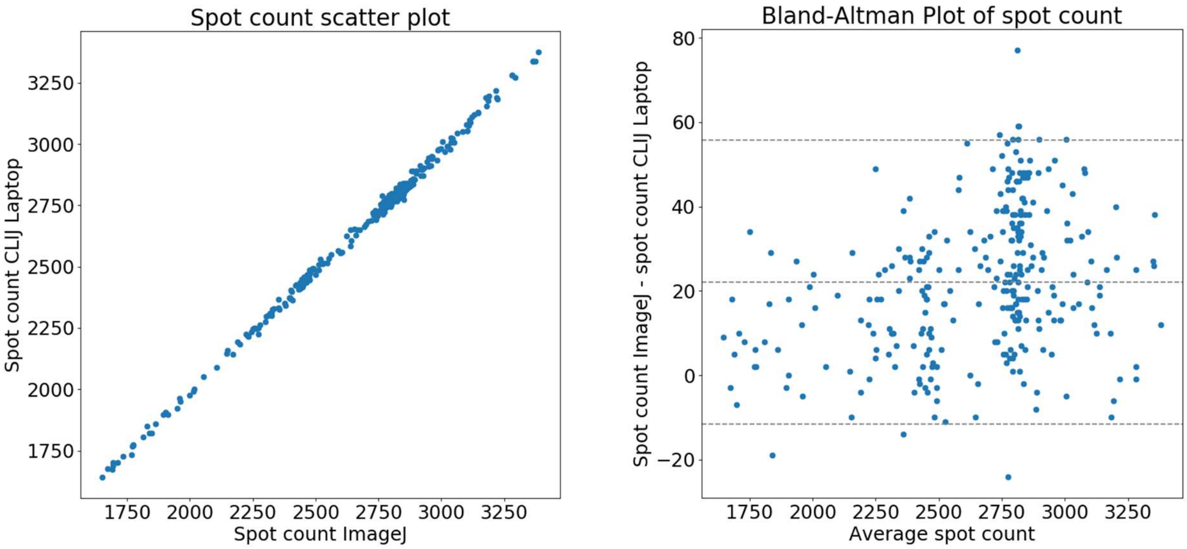


b)

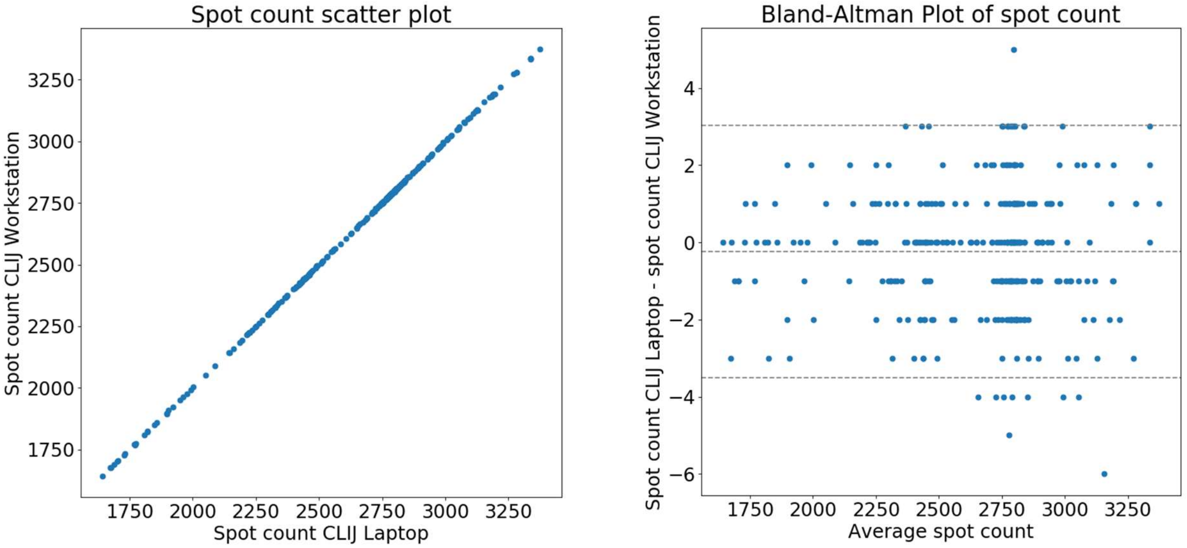


**Supplemental Figure 5:** Comparison of a) derived spot count using ImageJ and CLIJ workflows executed on the test laptop visualized as scatter plot and Bland-Altman plot. In b) differences are shown between the CLIJ workflows executed on laptop (Intel GPU) and workstation (Nvidia GPU). Measured differences are unique for the tested scenarios. They may vary on other operations, workflows, used hardware and/or data.
