## Supplementary Figure 6: Workflow results for "CLIJ: GPU-accelerated image processing for everyone"

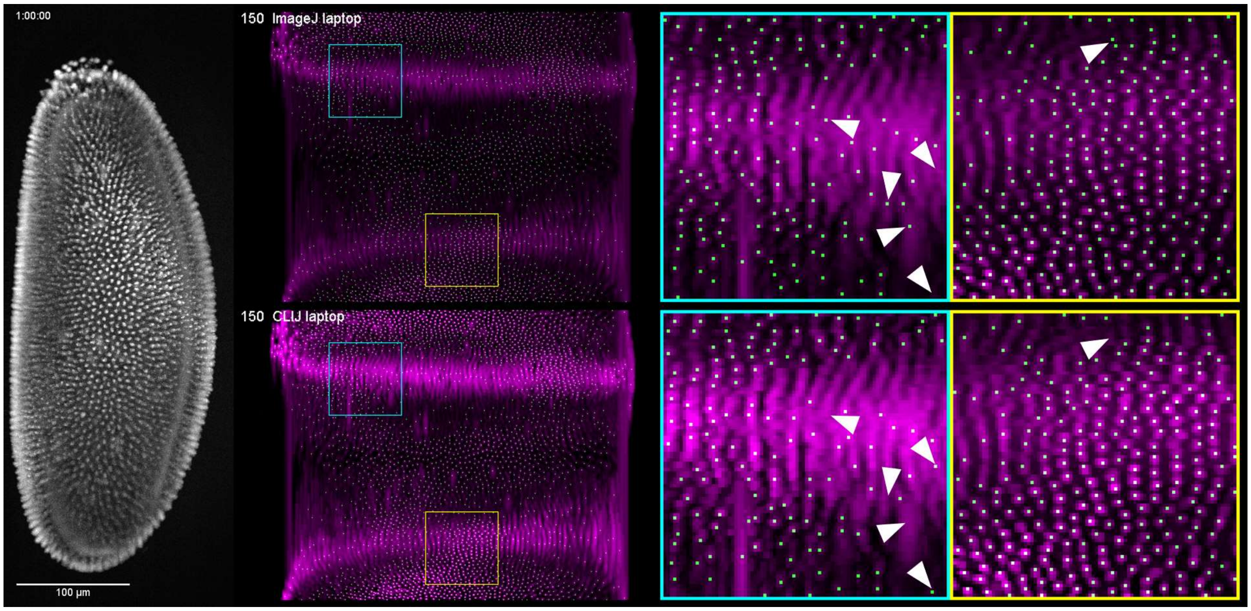


**Supplementary Figure 6**: The benchmarked workflow processes a 3D image stack showing a Drosophila melanogaster embryo expressing histone-RFP (left, shown as background-subtracted maximum projections), projects it on a 3D surface (center) and counts spots (green dots). Zoomed views show regions where spot detection is harder (cyan) than in other regions (yellow) because of limited differentiability of nuclei. In some regions differences, exemplary marked with white triangles, appear more often between both workflows (top: ImageJ, bottom: CLIJ).
