## Supplementary Figure 7: CLIJ Screenshot for "CLIJ: GPU-accelerated image processing for everyone"

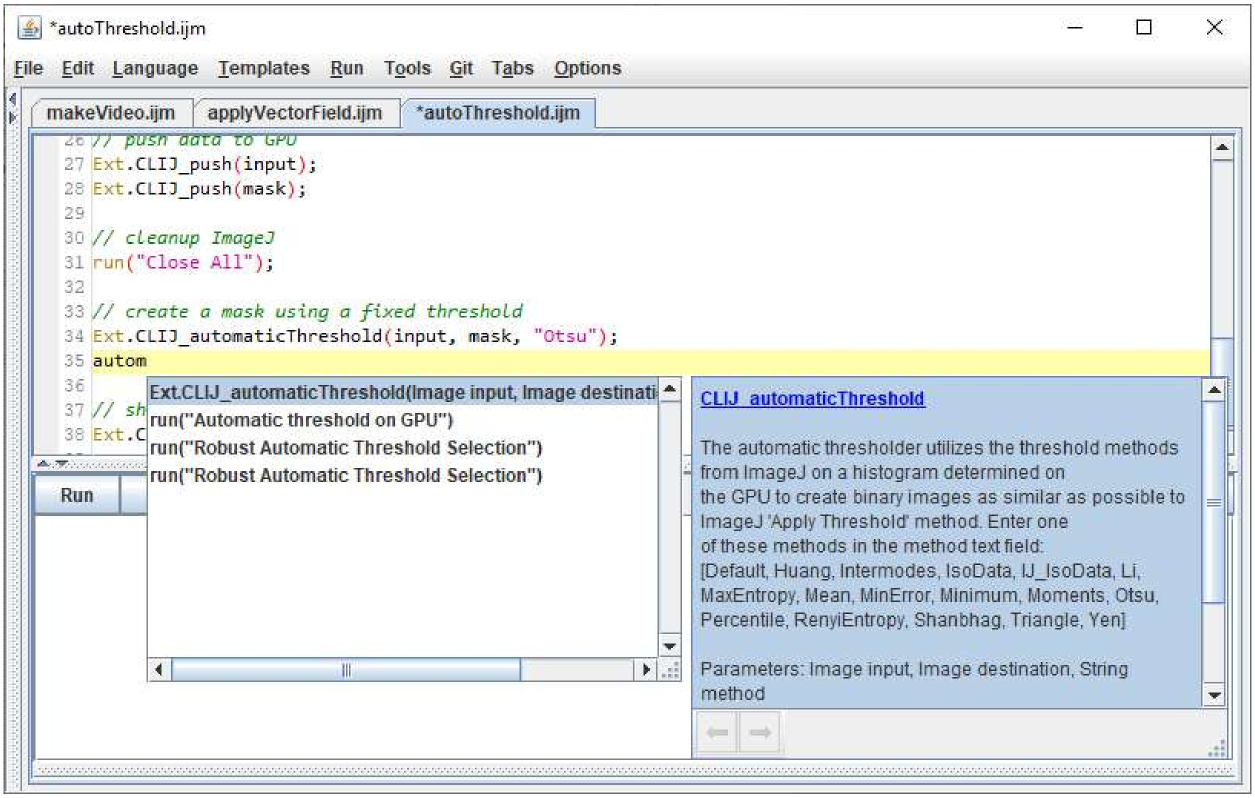


**Supplementary Figure 7:** Fijis script editor has been enriched with CLIJ specific auto-completion and online-help enabling users to assemble GPU-accelerated workflows without the need for reading the full documentation of CLIJ.
