## Supplementary Listing 1: CLIJ API Macro reference for "CLIJ: GPU-accelerated image processing for everyone"

### Supplementary Listing 1: CLIJ reference for ImageJ macro

#### CLIJ_absolute

Computes the absolute value of every individual pixel x in a given image.

```
f(x) = |x|
```

**Parameters**: Image source, Image destination

**Available for**: 2D, 3D

**Macro example**:

```
run("CLIJ Macro Extensions", "cl_device=");
Ext.CLIJ_push(source);
Ext.CLIJ_absolute(source, destination);
Ext.CLIJ_pull(destination);
```

#### CLIJ_addImageAndScalar

Adds a scalar value s to all pixels x of a given image X.

```
f(x, s) = x + s
```

**Parameters**: Image source, Image destination, Number scalar

**Available for**: 2D, 3D

**Macro example**:

```
run("CLIJ Macro Extensions", "cl_device=");
Ext.CLIJ_push(source);
Ext.CLIJ_addImageAndScalar(source, destination, scalar);
Ext.CLIJ_pull(destination);
```

#### CLIJ_addImages

Calculates the sum of pairs of pixels x and y of two images X and Y.

```
f(x, y) = x + y
```

**Parameters**: Image summand1, Image summand2, Image destination

**Available for**: 2D, 3D

**Macro example**:

```
run("CLIJ Macro Extensions", "cl_device=");
Ext.CLIJ_push(summand1);
Ext.CLIJ_push(summand2);
Ext.CLIJ_addImages(summand1, summand2, destination);
Ext.CLIJ_pull(destination);
```

### CLIJ_addImagesWeighted

Calculates the sum of pairs of pixels x and y from images X and Y weighted with factors a and b.

```
f(x, y, a, b) = x * a + y * b
```

**Parameters**: Image summand1, Image summand2, Image destination, Number factor1, Number factor2

**Available for**: 2D, 3D

**Macro example**:

```
run("CLIJ Macro Extensions", "cl_device=");
Ext.CLIJ_push(summand1);
Ext.CLIJ_push(summand2);
Ext.CLIJ_addImagesWeighted(summand1, summand2, destination, factor1, factor2);
Ext.CLIJ_pull(destination);
```

### CLIJ_affineTransform

CLIJ affineTransform is deprecated. Use affineTransform2D or affineTransform3D instead.

Applies an affine transform to a 3D image. Individual transforms must be separated by spaces.

Supported transforms:

- center: translate the coordinate origin to the center of the image
- -center: translate the coordinate origin back to the initial origin
- rotate=[angle]: rotate in X/Y plane (around Z-axis) by the given angle in degrees
- rotateX=[angle]: rotate in Y/Z plane (around X-axis) by the given angle in degrees

- rotateY=[angle]: rotate in X/Z plane (around Y-axis) by the given angle in degrees
- rotateZ=[angle]: rotate in X/Y plane (around Z-axis) by the given angle in degrees
- scale=[factor]: isotropic scaling according to given zoom factor
- scaleX=[factor]: scaling along X-axis according to given zoom factor
- scaleY=[factor]: scaling along Y-axis according to given zoom factor
- scaleZ=[factor]: scaling along Z-axis according to given zoom factor
- shearXY=[factor]: shearing along X-axis in XY plane according to given factor
- shearXZ=[factor]: shearing along X-axis in XZ plane according to given factor
- shearYX=[factor]: shearing along Y-axis in XY plane according to given factor
- shearYZ=[factor]: shearing along Y-axis in YZ plane according to given factor
- shearZX=[factor]: shearing along Z-axis in XZ plane according to given factor
- shearZY=[factor]: shearing along Z-axis in YZ plane according to given factor
- translateX=[distance]: translate along X-axis by distance given in pixels
- translateY=[distance]: translate along X-axis by distance given in pixels
- translateZ=[distance]: translate along X-axis by distance given in pixels

Example transform: transform = "center scale=2 rotate=45 -center";

**Parameters**: Image source, Image destination, String transform

**Available for**: 2D, 3D

**Macro example**:

```
run("CLIJ Macro Extensions", "cl_device=");
Ext.CLIJ_push(source);
Ext.CLIJ_affineTransform(source, destination, transform);
Ext.CLIJ_pull(destination);
```

### CLIJ_affineTransform2D

Applies an affine transform to a 2D image. Individual transforms must be separated by spaces.

Supported transforms:

- center: translate the coordinate origin to the center of the image
- -center: translate the coordinate origin back to the initial origin
- rotate=[angle]: rotate in X/Y plane (around Z-axis) by the given angle in degrees
- scale=[factor]: isotropic scaling according to given zoom factor
- scaleX=[factor]: scaling along X-axis according to given zoom factor
- scaleY=[factor]: scaling along Y-axis according to given zoom factor
- shearXY=[factor]: shearing along X-axis in XY plane according to given factor
- translateX=[distance]: translate along X-axis by distance given in pixels
- translateY=[distance]: translate along X-axis by distance given in pixels

Example transform: transform = "center scale=2 rotate=45 -center";

**Parameters**: Image source, Image destination, String transform

**Available for**: 2D

**Macro example**:

```
run("CLIJ Macro Extensions", "cl_device=");
Ext.CLIJ_push(source);
Ext.CLIJ_affineTransform2D(source, destination, transform);
Ext.CLIJ_pull(destination);
```

### CLIJ_affineTransform3D

Applies an affine transform to a 3D image. Individual transforms must be separated by spaces.

Supported transforms:

- center: translate the coordinate origin to the center of the image
- -center: translate the coordinate origin back to the initial origin
- rotate=[angle]: rotate in X/Y plane (around Z-axis) by the given angle in degrees
- rotateX=[angle]: rotate in Y/Z plane (around X-axis) by the given angle in degrees
- rotateY=[angle]: rotate in X/Z plane (around Y-axis) by the given angle in degrees
- rotateZ=[angle]: rotate in X/Y plane (around Z-axis) by the given angle in degrees
- scale=[factor]: isotropic scaling according to given zoom factor
- scaleX=[factor]: scaling along X-axis according to given zoom factor
- scaleY=[factor]: scaling along Y-axis according to given zoom factor
- scaleZ=[factor]: scaling along Z-axis according to given zoom factor
- shearXY=[factor]: shearing along X-axis in XY plane according to given factor
- shearXZ=[factor]: shearing along X-axis in XZ plane according to given factor
- shearYX=[factor]: shearing along Y-axis in XY plane according to given factor
- shearYZ=[factor]: shearing along Y-axis in YZ plane according to given factor
- shearZX=[factor]: shearing along Z-axis in XZ plane according to given factor
- shearZY=[factor]: shearing along Z-axis in YZ plane according to given factor
- translateX=[distance]: translate along X-axis by distance given in pixels
- translateY=[distance]: translate along X-axis by distance given in pixels
- translateZ=[distance]: translate along X-axis by distance given in pixels

Example transform: transform = "center scale=2 rotate=45 -center";

**Parameters**: Image source, Image destination, String transform

**Available for**: 3D

**Macro example**:

```
run("CLIJ Macro Extensions", "cl_device=");
Ext.CLIJ_push(source);
```

```
Ext.CLIJ_affineTransform3D(source, destination, transform);
Ext.CLIJ_pull(destination);
```

### CLIJ_applyVectorField2D

Deforms an image according to distances provided in the given vector images. It is recommended to use 32-bit images for input, output and vector images.

**Parameters**: Image source, Image vectorX, Image vectorY, Image destination

**Available for**: 2D

**Macro example**:

```
run("CLIJ Macro Extensions", "cl_device=");
Ext.CLIJ_push(source);
Ext.CLIJ_push(vectorX);
Ext.CLIJ_push(vectorY);
Ext.CLIJ_applyVectorField2D(source, vectorX, vectorY, destination);
Ext.CLIJ_pull(destination);
```

### CLIJ_applyVectorField3D

Image source, Image vectorX, Image vectorY, Image vectorZ, Image destination

**Parameters**: Deforms an image stack according to distances provided in the given vector image stacks. It is recommended to use 32-bit image stacks for input, output and vector image stacks.

**Available for**: 3D

**Macro example**:

```
run("CLIJ Macro Extensions", "cl_device=");
Ext.CLIJ_applyVectorField3D(an, and);
```

### CLIJ_argMaximumZProjection

Determines the maximum projection of an image stack along Z. Furthermore, another 2D image is generated with pixels containing the z-index where the maximum was found (zero based).

**Parameters**: Image source, Image destination_max, Image destination_arg_max

**Available for**: 3D

**Macro example**:

```
run("CLIJ Macro Extensions", "cl_device=");
Ext.CLIJ_push(source);
Ext.CLIJ_argMaximumZProjection(source, destination_max, destination_arg_max);
Ext.CLIJ_pull(destination_max);
Ext.CLIJ_pull(destination_arg_max);
```

### CLIJ_automaticThreshold

The automatic thresholder utilizes the threshold methods from ImageJ on a histogram determined on the GPU to create binary images as similar as possible to ImageJ 'Apply Threshold' method. Enter one of these methods in the method text field: [Default, Huang, Intermodes, IsoData, IJ_IsoData, Li, MaxEntropy, Mean, MinError, Minimum, Moments, Otsu, Percentile, RenyiEntropy, Shanbhag, Triangle, Yen]

**Parameters**: Image input, Image destination, String method

**Available for**: 2D, 3D

**Macro example**:

```
run("CLIJ Macro Extensions", "cl_device=");
Ext.CLIJ_push(input);
Ext.CLIJ_automaticThreshold(input, destination, method);
Ext.CLIJ_pull(destination);
```

### CLIJ_binaryAnd

Computes a binary image (containing pixel values 0 and 1) from two images X and Y by connecting pairs of pixels x and y with the binary AND operator &. All pixel values except 0 in the input images are interpreted as 1.

```
f(x, y) = x & y
```

**Parameters**: Image operand1, Image operand2, Image destination

**Available for**: 2D, 3D

**Macro example**:

```
run("CLIJ Macro Extensions", "cl_device=");
Ext.CLIJ_push(operand1);
Ext.CLIJ_push(operand2);
```

```
Ext.CLIJ_binaryAnd(operand1, operand2, destination);
Ext.CLIJ_pull(destination);
```

### CLIJ_binaryNot

Computes a binary image (containing pixel values 0 and 1) from an image X by negating its pixel values x using the binary NOT operator ! All pixel values except 0 in the input image are interpreted as 1.

```
f(x) = !x
```

**Parameters**: Image source, Image destination

**Available for**: 2D, 3D

**Macro example**:

```
run("CLIJ Macro Extensions", "cl_device=");
Ext.CLIJ_push(source);
Ext.CLIJ_binaryNot(source, destination);
Ext.CLIJ_pull(destination);
```

### CLIJ_binaryOr

Computes a binary image (containing pixel values 0 and 1) from two images X and Y by connecting pairs of pixels x and y with the binary OR operator |. All pixel values except 0 in the input images are interpreted as 1.

```
f(x, y) = x | y
```

**Parameters**: Image operand1, Image operand2, Image destination

**Available for**: 2D, 3D

**Macro example**:

```
run("CLIJ Macro Extensions", "cl_device=");
Ext.CLIJ_push(operand1);
Ext.CLIJ_push(operand2);
Ext.CLIJ_binaryOr(operand1, operand2, destination);
Ext.CLIJ_pull(destination);
```

### CLIJ_binaryXOr

Computes a binary image (containing pixel values 0 and 1) from two images X and Y by connecting pairs of pixels x and y with the binary operators AND &, OR | and NOT ! implementing the XOR operator. All pixel values except 0 in the input images are interpreted as 1.

```
f(x, y) = (x & !y) | (!x & y)
```

**Parameters**: Image operand1, Image operand2, Image destination

**Available for**: 2D, 3D

**Macro example**:

```
run("CLIJ Macro Extensions", "cl_device=");
Ext.CLIJ_push(operand1);
Ext.CLIJ_push(operand2);
Ext.CLIJ_binaryXOr(operand1, operand2, destination);
Ext.CLIJ_pull(destination);
```

### CLIJ_blur2D

Computes the Gaussian blurred image of an image given two sigma values in X and Y. Thus, the filterkernel can have non-isotropic shape.

The implementation is done separable. In case a sigma equals zero, the direction is not blurred.

**Parameters**: Image source, Image destination, Number sigmaX, Number sigmaY

**Available for**: 2D

**Macro example**:

```
run("CLIJ Macro Extensions", "cl_device=");
Ext.CLIJ_push(source);
Ext.CLIJ_blur2D(source, destination, sigmaX, sigmaY);
Ext.CLIJ_pull(destination);
```

### CLIJ_blur3D

Computes the Gaussian blurred image of an image given two sigma values in X, Y and Z. Thus, the filterkernel can have non-isotropic shape.

The implementation is done separable. In case a sigma equals zero, the direction is not blurred.

**Parameters**: Image source, Image destination, Number sigmaX, Number sigmaY, Number sigmaZ

**Available for**: 3D

**Macro example**:

```
run("CLIJ Macro Extensions", "cl_device=");
Ext.CLIJ_push(source);
Ext.CLIJ_blur3D(source, destination, sigmaX, sigmaY, sigmaZ);
Ext.CLIJ_pull(destination);
```

### CLIJ_blur3DSliceBySlice

Computes the Gaussian blurred image of an image given two sigma values in X and Y. Thus, the filterkernel can have non-isotropic shape.

The Gaussian blur is applied slice by slice in 2D.

**Parameters**: Image source, Image destination, Number sigmaX, Number sigmaY

**Available for**: 3D

**Macro example**:

```
run("CLIJ Macro Extensions", "cl_device=");
Ext.CLIJ_push(source);
Ext.CLIJ_blur3DSliceBySlice(source, destination, sigmaX, sigmaY);
Ext.CLIJ_pull(destination);
```

### CLIJ_centerOfMass

Determines the center of mass of an image or image stack and writes the result in the results table in the columns MassX, MassY and MassZ.

**Parameters**: Image source

**Available for**: 2D, 3D

**Macro example**:

```
run("CLIJ Macro Extensions", "cl_device=");
Ext.CLIJ_push(source);
Ext.CLIJ_centerOfMass(source);
```

### CLIJ_clInfo

Outputs information about available OpenCL devices.

**Parameters**:

**Available for**:

**Macro example**:

```
run("CLIJ Macro Extensions", "cl_device=");
Ext.CLIJ_clInfo();
```

### CLIJ_clear

Resets the GPUs memory by deleting all cached images.

**Parameters**:

**Available for**:

**Macro example**:

```
run("CLIJ Macro Extensions", "cl_device=");
Ext.CLIJ_clear();
```

### CLIJ_convertFloat

Convert the input image to a float image with 32 bits per pixel. The target image should not exist with a different type before this method is called.

**Parameters**: Image source, Image destination

**Available for**: 2D, 3D

**Macro example**:

```
run("CLIJ Macro Extensions", "cl_device=");
Ext.CLIJ_push(source);
Ext.CLIJ_convertFloat(source, destination);
Ext.CLIJ_pull(destination);
```

### CLIJ_convertUInt16

Convert the input image to a unsigned integer image with 16 bits per pixel. Pixel values are copied as they are. Use multiplyImageWithScalar in order to scalepixel values when reducing bit-depth to prevent cutting-off intensity ranges. The target image should not exist with a different type before this method is called.

**Parameters**: Image source, Image destination

**Available for**: 2D, 3D

**Macro example**:

```
run("CLIJ Macro Extensions", "cl_device=");
Ext.CLIJ_push(source);
Ext.CLIJ_convertUInt16(source, destination);
Ext.CLIJ_pull(destination);
```

### CLIJ_convertUInt8

Convert the input image to a unsigned integer image with 8 bits per pixel. Pixel values are copied as they are. Use multiplyImageWithScalar in order to scalepixel values when reducing bit-depth to prevent cutting-off intensity ranges. The target image should not exist with a different type before this method is called.

**Parameters**: Image source, Image destination

**Available for**: 2D, 3D

**Macro example**:

```
run("CLIJ Macro Extensions", "cl_device=");
Ext.CLIJ_push(source);
Ext.CLIJ_convertUInt8(source, destination);
Ext.CLIJ_pull(destination);
```

### CLIJ_copy

Copies an image.

```
f(x) = x
```

**Parameters**: Image source, Image destination

**Available for**: 2D, 3D

**Macro example**:

```
run("CLIJ Macro Extensions", "cl_device=");
Ext.CLIJ_push(source);
Ext.CLIJ_copy(source, destination);
Ext.CLIJ_pull(destination);
```

### CLIJ_copySlice

This method has two purposes: It copies a 2D image to a given slice z position in a 3D image stack or It copies a given slice at position z in an image stack to a 2D image.

The first case is only available via ImageJ macro. If you are using it, it is recommended that the target 3D image already pre-exists in GPU memory before calling this method. Otherwise, CLIJ create the image stack with z planes.

**Parameters**: Image source, Image destination, Number sliceIndex

**Available for**: 3D -> 2D and 2D -> 3D

**Macro example**:

```
run("CLIJ Macro Extensions", "cl_device=");
Ext.CLIJ_push(source);
Ext.CLIJ_copySlice(source, destination, sliceIndex);
Ext.CLIJ_pull(destination);
```

### CLIJ_countNonZeroPixels2DSphere

Counts non-zero pixels in a sphere around every pixel.Put the number in the result image.

**Parameters**: Image source, Image destination, Number radiusX, Number radiusY

**Available for**: 2D

**Macro example**:

```
run("CLIJ Macro Extensions", "cl_device=");
Ext.CLIJ_push(source);
Ext.CLIJ_countNonZeroPixels2DSphere(source, destination, radiusX, radiusY);
Ext.CLIJ_pull(destination);
```

### CLIJ_countNonZeroPixelsSliceBySliceSphere

Counts non-zero pixels in a sphere around every pixel slice by slice in a stack and puts the resulting number in the destination image stack.

**Parameters**: Image source, Image destination, Number radiusX, Number radiusY

**Available for**: 3D

**Macro example**:

```
run("CLIJ Macro Extensions", "cl_device=");
Ext.CLIJ_push(source);
Ext.CLIJ_countNonZeroPixelsSliceBySliceSphere(source, destination, radiusX, radiusY);
Ext.CLIJ_pull(destination);
```

### CLIJ_countNonZeroVoxels3DSphere

Counts non-zero voxels in a sphere around every voxel.Put the number in the result image.

**Parameters**: Image source, Image destination, Number radiusX, Number radiusY, Number radiusZ

**Available for**: 3D

**Macro example**:

```
run("CLIJ Macro Extensions", "cl_device=");
Ext.CLIJ_push(source);
Ext.CLIJ_countNonZeroVoxels3DSphere(source, destination, radiusX, radiusY, radiusZ);
Ext.CLIJ_pull(destination);
```

### CLIJ_create2D

Allocated memory for a new 2D image in the GPU memory. BitDepth must be 8 (unsigned byte), 16 (unsigned short) or 32 (float).

**Parameters**: Image destination, Number width, Number height, Number bitDepth

**Available for**: 2D

**Macro example**:

```
run("CLIJ Macro Extensions", "cl_device=");
Ext.CLIJ_create2D(destination, width, height, bitDepth);
Ext.CLIJ_pull(destination);
```

### CLIJ_create3D

Allocated memory for a new 3D image in the GPU memory. BitDepth must be 8 (unsigned byte), 16 (unsigned short) or 32 (float).

**Parameters**: Image destination, Number width, Number height, Number depth, Number bitDepth

**Available for**: 3D

**Macro example**:

```
run("CLIJ Macro Extensions", "cl_device=");
Ext.CLIJ_create3D(destination, width, height, depth, bitDepth);
Ext.CLIJ_pull(destination);
```

### CLIJ_crop2D

Crops a given rectangle out of a given image.

Note: If the destination image pre-exists already, it will be overwritten and keep it's dimensions.

**Parameters**: Image source, Image destination, Number startX, Number startY, Number width, Number height

**Available for**: 3D

**Macro example**:

```
run("CLIJ Macro Extensions", "cl_device=");
Ext.CLIJ_push(source);
Ext.CLIJ_crop2D(source, destination, startX, startY, width, height);
Ext.CLIJ_pull(destination);
```

### CLIJ_crop3D

Crops a given sub-stack out of a given image stack.

Note: If the destination image pre-exists already, it will be overwritten and keep it's dimensions.

**Parameters**: Image source, Image destination, Number startX, Number startY, Number startZ, Number width, Number height, Number depth

**Available for**: 3D

**Macro example**:

```
run("CLIJ Macro Extensions", "cl_device=");
Ext.CLIJ_push(source);
Ext.CLIJ_crop3D(source, destination, startX, startY, startZ, width, height, depth);
Ext.CLIJ_pull(destination);
```

### CLIJ_detectMaximaBox

Detects local maxima in a given square/cubic neighborhood. Pixels in the resulting image are set to 1 if there is no other pixel in a given radius which has a higher intensity, and to 0 otherwise.

**Parameters**: Image source, Image destination, Number radius

**Available for**: 2D, 3D

**Macro example**:

```
run("CLIJ Macro Extensions", "cl_device=");
Ext.CLIJ_push(source);
Ext.CLIJ_detectMaximaBox(source, destination, radius);
Ext.CLIJ_pull(destination);
```

### CLIJ_detectMaximaSliceBySliceBox

Detects local maxima in a given square neighborhood of an input image stack. The input image stack is processed slice by slice. Pixels in the resulting image are set to 1 if there is no other pixel in a given radius which has a higher intensity, and to 0 otherwise.

**Parameters**: Image source, Image destination, Number radius

**Available for**: 3D

**Macro example**:

```
run("CLIJ Macro Extensions", "cl_device=");
Ext.CLIJ_push(source);
Ext.CLIJ_detectMaximaSliceBySliceBox(source, destination, radius);
Ext.CLIJ_pull(destination);
```

### CLIJ_detectMinimaBox

Detects local minima in a given square/cubic neighborhood. Pixels in the resulting image are set to 1 if there is no other pixel in a given radius which has a lower intensity, and to 0 otherwise.

**Parameters**: Image source, Image destination, Number radius

**Available for**: 2D, 3D

**Macro example**:

```
run("CLIJ Macro Extensions", "cl_device=");
Ext.CLIJ_push(source);
Ext.CLIJ_detectMinimaBox(source, destination, radius);
Ext.CLIJ_pull(destination);
```

### CLIJ_detectMinimaSliceBySliceBox

Detects local minima in a given square neighborhood of an input image stack. The input image stack is processed slice by slice. Pixels in the resulting image are set to 1 if there is no other pixel in a given radius which has a lower intensity, and to 0 otherwise.

**Parameters**: Image source, Image destination, Number radius

**Available for**: 3D

**Macro example**:

```
run("CLIJ Macro Extensions", "cl_device=");
Ext.CLIJ_push(source);
Ext.CLIJ_detectMinimaSliceBySliceBox(source, destination, radius);
Ext.CLIJ_pull(destination);
```

### CLIJ_dilateBox

Computes a binary image with pixel values 0 and 1 containing the binary dilation of a given input image. The dilation takes the Moore-neighborhood (8 pixels in 2D and 26 pixels in 3d) into account. The pixels in the input image with pixel value not equal to 0 will be interpreted as 1.

This method is comparable to the 'Dilate' menu in ImageJ in case it is applied to a 2D image. The only difference is that the output image contains values 0 and 1 instead of 0 and 255.

**Parameters**: Image source, Image destination

**Available for**: 2D, 3D

**Macro example**:

```
run("CLIJ Macro Extensions", "cl_device=");
Ext.CLIJ_push(source);
Ext.CLIJ_dilateBox(source, destination);
Ext.CLIJ_pull(destination);
```

### CLIJ_dilateBoxSliceBySlice

Computes a binary image with pixel values 0 and 1 containing the binary dilation of a given input image. The dilation takes the Moore-neighborhood (8 pixels in 2D and 26 pixels in 3d) into account. The pixels in the input image with pixel value not equal to 0 will be interpreted as 1.

This method is comparable to the 'Dilate' menu in ImageJ in case it is applied to a 2D image. The only difference is that the output image contains values 0 and 1 instead of 0 and 255.

This filter is applied slice by slice in 2D.

**Parameters**: Image source, Image destination

**Available for**: 2D, 3D

**Macro example**:

```
run("CLIJ Macro Extensions", "cl_device=");
Ext.CLIJ_push(source);
Ext.CLIJ_dilateBoxSliceBySlice(source, destination);
Ext.CLIJ_pull(destination);
```

### CLIJ_dilateSphere

Computes a binary image with pixel values 0 and 1 containing the binary dilation of a given input image. The dilation takes the von-Neumann-neighborhood (4 pixels in 2D and 6 pixels in 3d) into account. The pixels in the input image with pixel value not equal to 0 will be interpreted as 1.

**Parameters**: Image source, Image destination

**Available for**: 2D, 3D

**Macro example**:

```
run("CLIJ Macro Extensions", "cl_device=");
Ext.CLIJ_push(source);
Ext.CLIJ_dilateSphere(source, destination);
Ext.CLIJ_pull(destination);
```

### CLIJ_dilateSphereSliceBySlice

Computes a binary image with pixel values 0 and 1 containing the binary dilation of a given input image. The dilation takes the von-Neumann-neighborhood (4 pixels in 2D and 6 pixels in 3d) into account. The pixels in the input image with pixel value not equal to 0 will be interpreted as 1.

This filter is applied slice by slice in 2D.

**Parameters**: Image source, Image destination

**Available for**: 2D, 3D

**Macro example**:

```
run("CLIJ Macro Extensions", "cl_device=");
Ext.CLIJ_push(source);
Ext.CLIJ_dilateSphereSliceBySlice(source, destination);
Ext.CLIJ_pull(destination);
```

### CLIJ_divideImages

Divides two images X and Y by each other pixel wise.

```
f(x, y) = x / y
```

**Parameters**: Image divident, Image divisor, Image destination

**Available for**: 2D, 3D

**Macro example**:

```
run("CLIJ Macro Extensions", "cl_device=");
Ext.CLIJ_push(divident);
Ext.CLIJ_push(divisor);
Ext.CLIJ_divideImages(divident, divisor, destination);
Ext.CLIJ_pull(destination);
```

### CLIJ_downsample2D

Scales an image using given scaling factors for X and Y dimensions. The nearest-neighbor method is applied. In ImageJ the method which is similar is called 'Interpolation method: none'.

**Parameters**: Image source, Image destination, Number factorX, Number factorY

**Available for**: 2D

**Macro example**:

```
run("CLIJ Macro Extensions", "cl_device=");
Ext.CLIJ_push(source);
Ext.CLIJ_downsample2D(source, destination, factorX, factorY);
Ext.CLIJ_pull(destination);
```

### CLIJ_downsample3D

Scales an image using given scaling factors for X and Y dimensions. The nearest-neighbor method is applied. In ImageJ the method which is similar is called 'Interpolation method: none'.

**Parameters**: Image source, Image destination, Number factorX, Number factorY, Number factorZ

**Available for**: 3D

**Macro example**:

```
run("CLIJ Macro Extensions", "cl_device=");
Ext.CLIJ_push(source);
Ext.CLIJ_downsample3D(source, destination, factorX, factorY, factorZ);
Ext.CLIJ_pull(destination);
```

### CLIJ_downsampleSliceBySliceHalfMedian

Scales an image using scaling factors 0.5 for X and Y dimensions. The Z dimension stays untouched. Thus, each slice is processed separately. The median method is applied. Thus, each pixel value in the destination image equals to the median of four corresponding pixels in the source image.

**Parameters**: Image source, Image destination

**Available for**: 3D

**Macro example**:

```
run("CLIJ Macro Extensions", "cl_device=");
Ext.CLIJ_push(source);
Ext.CLIJ_downsampleSliceBySliceHalfMedian(source, destination);
Ext.CLIJ_pull(destination);
```

### CLIJ_erodeBox

Computes a binary image with pixel values 0 and 1 containing the binary erosion of a given input image. The erosion takes the Moore-neighborhood (8 pixels in 2D and 26 pixels in 3d) into account. The pixels in the input image with pixel value not equal to 0 will be interpreted as 1.

This method is comparable to the 'Erode' menu in ImageJ in case it is applied to a 2D image. The only difference is that the output image contains values 0 and 1 instead of 0 and 255.

**Parameters**: Image source, Image destination

**Available for**: 2D, 3D

**Macro example**:

```
run("CLIJ Macro Extensions", "cl_device=");
Ext.CLIJ_push(source);
Ext.CLIJ_erodeBox(source, destination);
Ext.CLIJ_pull(destination);
```

### CLIJ_erodeBoxSliceBySlice

Computes a binary image with pixel values 0 and 1 containing the binary erosion of a given input image. The erosion takes the Moore-neighborhood (8 pixels in 2D and 26 pixels in 3d) into account. The pixels in the input image with pixel value not equal to 0 will be interpreted as 1.

This method is comparable to the 'Erode' menu in ImageJ in case it is applied to a 2D image. The only difference is that the output image contains values 0 and 1 instead of 0 and 255.

This filter is applied slice by slice in 2D.

**Parameters**: Image source, Image destination

**Available for**: 2D, 3D

**Macro example**:

```
run("CLIJ Macro Extensions", "cl_device=");
Ext.CLIJ_push(source);
Ext.CLIJ_erodeBoxSliceBySlice(source, destination);
Ext.CLIJ_pull(destination);
```

### CLIJ_erodeSphere

Computes a binary image with pixel values 0 and 1 containing the binary erosion of a given input image. The erosion takes the von-Neumann-neighborhood (4 pixels in 2D and 6 pixels in 3d) into account. The pixels in the input image with pixel value not equal to 0 will be interpreted as 1.

**Parameters**: Image source, Image destination

**Available for**: 2D, 3D

**Macro example**:

```
run("CLIJ Macro Extensions", "cl_device=");
Ext.CLIJ_push(source);
Ext.CLIJ_erodeSphere(source, destination);
Ext.CLIJ_pull(destination);
```

### CLIJ_erodeSphereSliceBySlice

Computes a binary image with pixel values 0 and 1 containing the binary erosion of a given input image. The erosion takes the von-Neumann-neighborhood (4 pixels in 2D and 6 pixels in 3d) into account. The pixels in the input image with pixel value not equal to 0 will be interpreted as 1.

This filter is applied slice by slice in 2D.

**Parameters**: Image source, Image destination

**Available for**: 2D, 3D

**Macro example**:

```
run("CLIJ Macro Extensions", "cl_device=");
Ext.CLIJ_push(source);
Ext.CLIJ_erodeSphereSliceBySlice(source, destination);
Ext.CLIJ_pull(destination);
```

### CLIJ_flip2D

Flips an image in X and/or Y direction depending on boolean flags.

**Parameters**: Image source, Image destination, Boolean flipX, Boolean flipY

**Available for**: 2D

**Macro example**:

```
run("CLIJ Macro Extensions", "cl_device=");
Ext.CLIJ_push(source);
Ext.CLIJ_flip2D(source, destination, flipX, flipY);
Ext.CLIJ_pull(destination);
```

### CLIJ_flip3D

Flips an image in X, Y and/or Z direction depending on boolean flags.

**Parameters**: Image source, Image destination, Boolean flipX, Boolean flipY, Boolean flipZ

**Available for**: 3D

**Macro example**:

```
run("CLIJ Macro Extensions", "cl_device=");
Ext.CLIJ_push(source);
Ext.CLIJ_flip3D(source, destination, flipX, flipY, flipZ);
Ext.CLIJ_pull(destination);
```

### CLIJ_gradientX

Computes the gradient of gray values along X. Assuming a, b and c are three adjacent pixels in X direction. In the target image will be saved as:

```
b' = c - a;
```

**Parameters**: Image source, Image destination

**Available for**: 2D, 3D

**Macro example**:

```
run("CLIJ Macro Extensions", "cl_device=");
Ext.CLIJ_push(source);
Ext.CLIJ_gradientX(source, destination);
Ext.CLIJ_pull(destination);
```

### CLIJ_gradientY

Computes the gradient of gray values along Y. Assuming a, b and c are three adjacent pixels in Y direction. In the target image will be saved as:

```
b' = c - a;
```

**Parameters**: Image source, Image destination

**Available for**: 2D, 3D

**Macro example**:

```
run("CLIJ Macro Extensions", "cl_device=");
Ext.CLIJ_push(source);
Ext.CLIJ_gradientY(source, destination);
Ext.CLIJ_pull(destination);
```

### CLIJ_gradientZ

Computes the gradient of gray values along Z. Assuming a, b and c are three adjacent pixels in Z direction. In the target image will be saved as:

```
b' = c - a;
```

**Parameters**: Image source, Image destination

**Available for**: 3D

**Macro example**:

```
run("CLIJ Macro Extensions", "cl_device=");
Ext.CLIJ_push(source);
Ext.CLIJ_gradientZ(source, destination);
Ext.CLIJ_pull(destination);
```

### CLIJ_help

Searches in the list of CLIJ commands for a given pattern. Lists all commands in case"" is handed over as parameter.

**Parameters**: String searchFor

**Available for**: -

**Macro example**:

```
run("CLIJ Macro Extensions", "cl_device=");
Ext.CLIJ_help(searchFor);
```

### CLIJ_histogram

Determines the histogram of a given image.

**Parameters**: Image source, Image destination, Number numberOfBins, Number minimumGreyValue, Number maximumGreyValue, Boolean determineMinAndMax

**Available for**: 2D, 3D

**Macro example**:

```
run("CLIJ Macro Extensions", "cl_device=");
Ext.CLIJ_push(source);
Ext.CLIJ_histogram(source, destination, numberOfBins, minimumGreyValue,
maximumGreyValue, determineMinAndMax);
Ext.CLIJ_pull(destination);
```

### CLIJ_invert

Computes the negative value of all pixels in a given image. It is recommended to convert images to 32-bit float before applying this operation.

```
f(x) = - x
```

For binary images, use binaryNot.

**Parameters**: Image source, Image destination

**Available for**: 2D, 3D

**Macro example**:

```
run("CLIJ Macro Extensions", "cl_device=");
Ext.CLIJ_push(source);
Ext.CLIJ_invert(source, destination);
Ext.CLIJ_pull(destination);
```

### CLIJ_localThreshold

Computes a binary image with pixel values 0 and 1 depending on if a pixel value x in image X was above of equal to the pixel value m in mask image M.

```
f(x) = (1 if (x >=  m)); (0 otherwise)
```

**Parameters**: Image source, Image localThreshold, Image destination

**Available for**: 2D, 3D

**Macro example**:

```
run("CLIJ Macro Extensions", "cl_device=");
Ext.CLIJ_push(source);
Ext.CLIJ_push(localThreshold);
Ext.CLIJ_localThreshold(source, localThreshold, destination);
Ext.CLIJ_pull(destination);
```

### CLIJ_mask

Computes a masked image by applying a mask to an image. All pixel values x of image X will be copied to the destination image in case pixel value m at the same position in the mask image is not equal to zero.

```
f(x,m) = (x if (m != 0); (0 otherwise))
```

**Parameters**: Image source, Image mask, Image destination

**Available for**: 2D, 3D

**Macro example**:

```
run("CLIJ Macro Extensions", "cl_device=");
Ext.CLIJ_push(source);
Ext.CLIJ_push(mask);
Ext.CLIJ_mask(source, mask, destination);
Ext.CLIJ_pull(destination);
```

### CLIJ_maskStackWithPlane

Computes a masked image by applying a 2D mask to an image stack. All pixel values x of image X will be copied to the destination image in case pixel value m at the same spatial position in the mask image is not equal to zero.

```
f(x,m) = (x if (m != 0); (0 otherwise))
```

**Parameters**: Image source, Image mask, Image destination

**Available for**: 3D (first parameter), 2D (second parameter), 3D (result)

**Macro example**:

```
run("CLIJ Macro Extensions", "cl_device=");
Ext.CLIJ_push(source);
Ext.CLIJ_push(mask);
Ext.CLIJ_maskStackWithPlane(source, mask, destination);
Ext.CLIJ_pull(destination);
```

### CLIJ_maximum2DBox

Computes the local maximum of a pixels rectangular neighborhood. The rectangles size is specified by its half-width and half-height (radius).

**Parameters**: Image source, Image destination, Number radiusX, Number radiusY

**Available for**: 2D

**Macro example**:

```
run("CLIJ Macro Extensions", "cl_device=");
Ext.CLIJ_push(source);
Ext.CLIJ_maximum2DBox(source, destination, radiusX, radiusY);
Ext.CLIJ_pull(destination);
```

### CLIJ_maximum2DSphere

Computes the local maximum of a pixels ellipsoidal neighborhood. The ellipses size is specified by its half-width and half-height (radius).

**Parameters**: Image source, Image destination, Number radiusX, Number radiusY

**Available for**: 2D

**Macro example**:

```
run("CLIJ Macro Extensions", "cl_device=");
Ext.CLIJ_push(source);
Ext.CLIJ_maximum2DSphere(source, destination, radiusX, radiusY);
Ext.CLIJ_pull(destination);
```

### CLIJ_maximum3DBox

Computes the local maximum of a pixels cube neighborhood. The cubes size is specified by its half-width, half-height and half-depth (radius).

**Parameters**: Image source, Image destination, Number radiusX, Number radiusY, Number radiusZ

**Available for**: 3D

**Macro example**:

```
run("CLIJ Macro Extensions", "cl_device=");
Ext.CLIJ_push(source);
Ext.CLIJ_maximum3DBox(source, destination, radiusX, radiusY, radiusZ);
Ext.CLIJ_pull(destination);
```

### CLIJ_maximum3DSphere

Computes the local maximum of a pixels spherical neighborhood. The spheres size is specified by its half-width, half-height and half-depth (radius).

**Parameters**: Image source, Image destination, Number radiusX, Number radiusY, Number radiusZ

**Available for**: 3D

**Macro example**:

```
run("CLIJ Macro Extensions", "cl_device=");
Ext.CLIJ_push(source);
Ext.CLIJ_maximum3DSphere(source, destination, radiusX, radiusY, radiusZ);
Ext.CLIJ_pull(destination);
```

### CLIJ_maximumImageAndScalar

Computes the maximum of a constant scalar s and each pixel value x in a given image X.

```
f(x, s) = max(x, s)
```

**Parameters**: Image source, Image destination, Number scalar

**Available for**: 2D, 3D

**Macro example**:

```
run("CLIJ Macro Extensions", "cl_device=");
Ext.CLIJ_push(source);
Ext.CLIJ_maximumImageAndScalar(source, destination, scalar);
Ext.CLIJ_pull(destination);
```

### CLIJ_maximumImages

Computes the maximum of a pair of pixel values x, y from two given images X and Y.

```
f(x, y) = max(x, y)
```

**Parameters**: Image source1, Image source2, Image destination

**Available for**: 2D, 3D

**Macro example**:

```
run("CLIJ Macro Extensions", "cl_device=");
Ext.CLIJ_push(source1);
Ext.CLIJ_push(source2);
Ext.CLIJ_maximumImages(source1, source2, destination);
Ext.CLIJ_pull(destination);
```

### CLIJ_maximumOfAllPixels

Determines the maximum of all pixels in a given image. It will be stored in a new row of ImageJs Results table in the column 'Max'.

**Parameters**: Image source

**Available for**: 2D, 3D

**Macro example**:

```
run("CLIJ Macro Extensions", "cl_device=");
Ext.CLIJ_push(source);
Ext.CLIJ_maximumOfAllPixels(source);
```

### CLIJ_maximumSliceBySliceSphere

Computes the local maximum of a pixels ellipsoidal 2D neighborhood in an image stack slice by slice. The ellipses size is specified by its half-width and half-height (radius).

This filter is applied slice by slice in 2D.

**Parameters**: Image source, Image destination, Number radiusX, Number radiusY

**Available for**: 3D

**Macro example**:

```
run("CLIJ Macro Extensions", "cl_device=");
Ext.CLIJ_push(source);
Ext.CLIJ_maximumSliceBySliceSphere(source, destination, radiusX, radiusY);
Ext.CLIJ_pull(destination);
```

### CLIJ_maximumXYZProjection

Determines the maximum projection of an image along a given dimension. Furthermore, the X and Y dimesions of the resulting image must be specified by the user according to its definition: X = 0 Y = 1 Z = 2

**Parameters**: Image source, Image destination_max, Number dimensionX, Number dimensionY, Number projectedDimension

**Available for**: 3D

**Macro example**:

```
run("CLIJ Macro Extensions", "cl_device=");
Ext.CLIJ_push(source);
Ext.CLIJ_maximumXYZProjection(source, destination_max, dimensionX, dimensionY,
projectedDimension);
Ext.CLIJ_pull(destination_max);
```

### CLIJ_maximumZProjection

Determines the maximum projection of an image along Z.

**Parameters**: Image source, Image destination_max

**Available for**: 3D

**Macro example**:

```
run("CLIJ Macro Extensions", "cl_device=");
Ext.CLIJ_push(source);
Ext.CLIJ_maximumZProjection(source, destination_max);
Ext.CLIJ_pull(destination_max);
```

### CLIJ_mean2DBox

Computes the local mean average of a pixels rectangular neighborhood. The rectangles size is specified by its half-width and half-height (radius).

**Parameters**: Image source, Image destination, Number radiusX, Number radiusY

**Available for**: 2D

**Macro example**:

```
run("CLIJ Macro Extensions", "cl_device=");
Ext.CLIJ_push(source);
Ext.CLIJ_mean2DBox(source, destination, radiusX, radiusY);
Ext.CLIJ_pull(destination);
```

### CLIJ_mean2DSphere

Computes the local mean average of a pixels ellipsoidal neighborhood. The ellipses size is specified by its half-width and half-height (radius).

**Parameters**: Image source, Image destination, Number radiusX, Number radiusY

**Available for**: 2D

**Macro example**:

```
run("CLIJ Macro Extensions", "cl_device=");
Ext.CLIJ_push(source);
Ext.CLIJ_mean2DSphere(source, destination, radiusX, radiusY);
Ext.CLIJ_pull(destination);
```

### CLIJ_mean3DBox

Computes the local mean average of a pixels cube neighborhood. The cubes size is specified by its half-width, half-height and half-depth (radius).

**Parameters**: Image source, Image destination, Number radiusX, Number radiusY, Number radiusZ

**Available for**: 3D

**Macro example**:

```
run("CLIJ Macro Extensions", "cl_device=");
Ext.CLIJ_push(source);
Ext.CLIJ_mean3DBox(source, destination, radiusX, radiusY, radiusZ);
Ext.CLIJ_pull(destination);
```

### CLIJ_mean3DSphere

Computes the local mean average of a pixels spherical neighborhood. The spheres size is specified by its half-width, half-height and half-depth (radius).

**Parameters**: Image source, Image destination, Number radiusX, Number radiusY, Number radiusZ

**Available for**: 3D

**Macro example**:

```
run("CLIJ Macro Extensions", "cl_device=");
Ext.CLIJ_push(source);
Ext.CLIJ_mean3DSphere(source, destination, radiusX, radiusY, radiusZ);
Ext.CLIJ_pull(destination);
```

### CLIJ_meanOfAllPixels

Determines the mean average of all pixels in a given image. It will be stored in a new row of ImageJs Results table in the column 'Mean'.

**Parameters**: Image source

**Available for**: 2D, 3D

**Macro example**:

```
run("CLIJ Macro Extensions", "cl_device=");
Ext.CLIJ_push(source);
Ext.CLIJ_meanOfAllPixels(source);
```

### CLIJ_meanSliceBySliceSphere

Computes the local mean average of a pixels ellipsoidal 2D neighborhood in an image stack slice by slice. The ellipses size is specified by its half-width and half-height (radius).

This filter is applied slice by slice in 2D.

**Parameters**: Image source, Image destination, Number radiusX, Number radiusY

**Available for**: 3D

**Macro example**:

```
run("CLIJ Macro Extensions", "cl_device=");
Ext.CLIJ_push(source);
Ext.CLIJ_meanSliceBySliceSphere(source, destination, radiusX, radiusY);
Ext.CLIJ_pull(destination);
```

### CLIJ_meanZProjection

Determines the mean average projection of an image along Z.

**Parameters**: Image source, Image destination

**Available for**: 3D

**Macro example**:

```
run("CLIJ Macro Extensions", "cl_device=");
Ext.CLIJ_push(source);
Ext.CLIJ_meanZProjection(source, destination);
Ext.CLIJ_pull(destination);
```

### CLIJ_median2DBox

Computes the local median of a pixels rectangular neighborhood. The rectangle is specified by its half-width and half-height (radius).

For technical reasons, the area of the rectangle must have less than 1000 pixels.

**Parameters**: Image source, Image destination, Number radiusX, Number radiusY

**Available for**: 2D

**Macro example**:

```
run("CLIJ Macro Extensions", "cl_device=");
Ext.CLIJ_push(source);
Ext.CLIJ_median2DBox(source, destination, radiusX, radiusY);
Ext.CLIJ_pull(destination);
```

### CLIJ_median2DSphere

Computes the local median of a pixels ellipsoidal neighborhood. The ellipses size is specified by its half-width and half-height (radius).

For technical reasons, the area of the ellipse must have less than 1000 pixels.

**Parameters**: Image source, Image destination, Number radiusX, Number radiusY

**Available for**: 2D

**Macro example**:

```
run("CLIJ Macro Extensions", "cl_device=");
Ext.CLIJ_push(source);
Ext.CLIJ_median2DSphere(source, destination, radiusX, radiusY);
Ext.CLIJ_pull(destination);
```

### CLIJ_median3DBox

Computes the local median of a pixels cuboid neighborhood. The cuboid size is specified by its half-width, half-height and half-depth (radius).

For technical reasons, the volume of the cuboid must contain less than 1000 voxels.

**Parameters**: Image source, Image destination, Number radiusX, Number radiusY, Number radiusZ

**Available for**: 3D

**Macro example**:

```
run("CLIJ Macro Extensions", "cl_device=");
Ext.CLIJ_push(source);
Ext.CLIJ_median3DBox(source, destination, radiusX, radiusY, radiusZ);
Ext.CLIJ_pull(destination);
```

### CLIJ_median3DSphere

Computes the local median of a pixels spherical neighborhood. The spheres size is specified by its half-width, half-height and half-depth (radius).

For technical reasons, the volume of the sphere must contain less than 1000 voxels.

**Parameters**: Image source, Image destination, Number radiusX, Number radiusY, Number radiusZ

**Available for**: 3D

**Macro example**:

```
run("CLIJ Macro Extensions", "cl_device=");
Ext.CLIJ_push(source);
Ext.CLIJ_median3DSphere(source, destination, radiusX, radiusY, radiusZ);
Ext.CLIJ_pull(destination);
```

### CLIJ_medianSliceBySliceBox

Computes the local median of a pixels rectangular neighborhood. This is done slice-by-slice in a 3D image stack. The rectangle is specified by its half-width and half-height (radius).

For technical reasons, the area of the rectangle must have less than 1000 pixels.

**Parameters**: Image source, Image destination, Number radiusX, Number radiusY

**Available for**: 3D

**Macro example**:

```
run("CLIJ Macro Extensions", "cl_device=");
Ext.CLIJ_push(source);
Ext.CLIJ_medianSliceBySliceBox(source, destination, radiusX, radiusY);
Ext.CLIJ_pull(destination);
```

### CLIJ_medianSliceBySliceSphere

Computes the local median of a pixels ellipsoidal neighborhood. This is done slice-by-slice in a 3D image stack. The ellipses size is specified by its half-width and half-height (radius).

For technical reasons, the area of the ellipse must have less than 1000 pixels.

**Parameters**: Image source, Image destination, Number radiusX, Number radiusY

**Available for**: 3D

**Macro example**:

```
run("CLIJ Macro Extensions", "cl_device=");
```

```
Ext.CLIJ_push(source);
Ext.CLIJ_medianSliceBySliceSphere(source, destination, radiusX, radiusY);
Ext.CLIJ_pull(destination);
```

### CLIJ_minimum2DBox

Computes the local minimum of a pixels rectangular neighborhood. The rectangles size is specified by its half-width and half-height (radius).

**Parameters**: Image source, Image destination, Number radiusX, Number radiusY

**Available for**: 2D

**Macro example**:

```
run("CLIJ Macro Extensions", "cl_device=");
Ext.CLIJ_push(source);
Ext.CLIJ_minimum2DBox(source, destination, radiusX, radiusY);
Ext.CLIJ_pull(destination);
```

### CLIJ_minimum2DSphere

Computes the local minimum of a pixels ellipsoidal neighborhood. The ellipses size is specified by its half-width and half-height (radius).

**Parameters**: Image source, Image destination, Number radiusX, Number radiusY

**Available for**: 2D

**Macro example**:

```
run("CLIJ Macro Extensions", "cl_device=");
Ext.CLIJ_push(source);
Ext.CLIJ_minimum2DSphere(source, destination, radiusX, radiusY);
Ext.CLIJ_pull(destination);
```

### CLIJ_minimum3DBox

Computes the local minimum of a pixels cube neighborhood. The cubes size is specified by its half-width, half-height and half-depth (radius).

**Parameters**: Image source, Image destination, Number radiusX, Number radiusY, Number radiusZ

**Available for**: 3D

**Macro example**:

```
run("CLIJ Macro Extensions", "cl_device=");
Ext.CLIJ_push(source);
Ext.CLIJ_minimum3DBox(source, destination, radiusX, radiusY, radiusZ);
Ext.CLIJ_pull(destination);
```

### CLIJ_minimum3DSphere

Computes the local minimum of a pixels spherical neighborhood. The spheres size is specified by its half-width, half-height and half-depth (radius).

**Parameters**: Image source, Image destination, Number radiusX, Number radiusY, Number radiusZ

**Available for**: 3D

**Macro example**:

```
run("CLIJ Macro Extensions", "cl_device=");
Ext.CLIJ_push(source);
Ext.CLIJ_minimum3DSphere(source, destination, radiusX, radiusY, radiusZ);
Ext.CLIJ_pull(destination);
```

### CLIJ_minimumImageAndScalar

Computes the maximum of a constant scalar s and each pixel value x in a given image X.

```
f(x, s) = min(x, s)
```

**Parameters**: Image source, Image destination, Number scalar

**Available for**: 2D, 3D

**Macro example**:

```
run("CLIJ Macro Extensions", "cl_device=");
Ext.CLIJ_push(source);
Ext.CLIJ_minimumImageAndScalar(source, destination, scalar);
Ext.CLIJ_pull(destination);
```

### CLIJ_minimumImages

Computes the minimum of a pair of pixel values x, y from two given images X and Y.

```
f(x, y) = min(x, y)
```

**Parameters**: Image source1, Image source2, Image destination

**Available for**: 2D, 3D

**Macro example**:

```
run("CLIJ Macro Extensions", "cl_device=");
Ext.CLIJ_push(source1);
Ext.CLIJ_push(source2);
Ext.CLIJ_minimumImages(source1, source2, destination);
Ext.CLIJ_pull(destination);
```

### CLIJ_minimumOfAllPixels

Determines the minimum of all pixels in a given image. It will be stored in a new row of ImageJs Results table in the column 'Min'.

**Parameters**: Image source

**Available for**: 2D, 3D

**Macro example**:

```
run("CLIJ Macro Extensions", "cl_device=");
Ext.CLIJ_push(source);
Ext.CLIJ_minimumOfAllPixels(source);
```

### CLIJ_minimumSliceBySliceSphere

Computes the local minimum of a pixels ellipsoidal 2D neighborhood in an image stack slice by slice. The ellipses size is specified by its half-width and half-height (radius).

This filter is applied slice by slice in 2D.

**Parameters**: Image source, Image destination, Number radiusX, Number radiusY

**Available for**: 3D

**Macro example**:

```
run("CLIJ Macro Extensions", "cl_device=");
Ext.CLIJ_push(source);
Ext.CLIJ_minimumSliceBySliceSphere(source, destination, radiusX, radiusY);
Ext.CLIJ_pull(destination);
```

### CLIJ_minimumZProjection

Determines the minimum projection of an image along Z.

**Parameters**: Image source, Image destination_sum

**Available for**: 3D

**Macro example**:

```
run("CLIJ Macro Extensions", "cl_device=");
Ext.CLIJ_push(source);
Ext.CLIJ_minimumZProjection(source, destination_sum);
Ext.CLIJ_pull(destination_sum);
```

### CLIJ_multiplyImageAndScalar

Multiplies all pixels value x in a given image X with a constant scalar s.

```
f(x, s) = x * s
```

**Parameters**: Image source, Image destination, Number scalar

**Available for**: 2D, 3D

**Macro example**:

```
run("CLIJ Macro Extensions", "cl_device=");
Ext.CLIJ_push(source);
Ext.CLIJ_multiplyImageAndScalar(source, destination, scalar);
Ext.CLIJ_pull(destination);
```

### CLIJ_multiplyImages

Multiplies all pairs of pixel values x and y from two image X and Y.

```
f(x, y) = x * y
```

**Parameters**: Image factor1, Image factor2, Image destination

**Available for**: 2D, 3D

**Macro example**:

```
run("CLIJ Macro Extensions", "cl_device=");
Ext.CLIJ_push(factor1);
Ext.CLIJ_push(factor2);
Ext.CLIJ_multiplyImages(factor1, factor2, destination);
Ext.CLIJ_pull(destination);
```

### CLIJ_multiplyStackWithPlane

Multiplies all pairs of pixel values x and y from an image stack X and a 2D image Y. x and y are at the same spatial position within a plane.

```
f(x, y) = x * y
```

**Parameters**: Image sourceStack, Image sourcePlane, Image destination

**Available for**: 3D (first parameter), 2D (second parameter), 3D (result)

**Macro example**:

```
run("CLIJ Macro Extensions", "cl_device=");
Ext.CLIJ_push(sourceStack);
Ext.CLIJ_push(sourcePlane);
Ext.CLIJ_multiplyStackWithPlane(sourceStack, sourcePlane, destination);
Ext.CLIJ_pull(destination);
```

### CLIJ_power

Computes all pixels value x to the power of a given exponent a.

```
f(x, a) = x ^ a
```

**Parameters**: Image source, Image destination, Number exponent

**Available for**: 2D, 3D

**Macro example**:

```
run("CLIJ Macro Extensions", "cl_device=");
Ext.CLIJ_push(source);
Ext.CLIJ_power(source, destination, exponent);
Ext.CLIJ_pull(destination);
```

### CLIJ_pull

Copies an image specified by its name from GPU memory back to ImageJ and shows it.

**Parameters**: String image

**Available for**: 2D, 3D

**Macro example**:

```
run("CLIJ Macro Extensions", "cl_device=");
Ext.CLIJ_pull(image);
```

### CLIJ_pullBinary

Copies a binary image specified by its name from GPU memory back to ImageJ and shows it. This binary image will have 0 and 255 pixel intensities as needed for ImageJ to interpret it as binary.

**Parameters**: String image

**Available for**: 2D, 3D

**Macro example**:

```
run("CLIJ Macro Extensions", "cl_device=");
Ext.CLIJ_pullBinary(image);
```

### CLIJ_push

Copies an image specified by its name to GPU memory in order to process it there later.

**Parameters**: String image

**Available for**: 2D, 3D

**Macro example**:

```
run("CLIJ Macro Extensions", "cl_device=");
Ext.CLIJ_push(image);
```

### CLIJ_pushCurrentSlice

Copies an image specified by its name to GPU memory in order to process it there later.

**Parameters**: String image

**Available for**: 2D, 3D

**Macro example**:

```
run("CLIJ Macro Extensions", "cl_device=");
Ext.CLIJ_pushCurrentSlice(image);
```

### CLIJ_release

Frees memory of a specified image in GPU memory.

**Parameters**: String image

**Available for**: 2D, 3D

**Macro example**:

```
run("CLIJ Macro Extensions", "cl_device=");
Ext.CLIJ_release(image);
```

### CLIJ_reportMemory

Prints a list of all images cached in the GPU to ImageJs log window together with a sum of memory consumption.

**Parameters**:

**Available for**: -

**Macro example**:

```
run("CLIJ Macro Extensions", "cl_device=");
Ext.CLIJ_reportMemory();
```

### CLIJ_resliceBottom

Flippes Y and Z axis of an image stack. This operation is similar to ImageJs 'Reslice [/]' method but offers less flexibility such as interpolation.

**Parameters**: Image source, Image destination

**Available for**: 3D

**Macro example**:

```
run("CLIJ Macro Extensions", "cl_device=");
Ext.CLIJ_push(source);
Ext.CLIJ_resliceBottom(source, destination);
Ext.CLIJ_pull(destination);
```

### CLIJ_resliceLeft

Flippes X, Y and Z axis of an image stack. This operation is similar to ImageJs 'Reslice [/]' method but offers less flexibility such as interpolation.

**Parameters**: Image source, Image destination

**Available for**: 3D

**Macro example**:

```
run("CLIJ Macro Extensions", "cl_device=");
Ext.CLIJ_push(source);
Ext.CLIJ_resliceLeft(source, destination);
Ext.CLIJ_pull(destination);
```

### CLIJ_resliceRadial

Computes a radial projection of an image stack. Starting point for the line is the center in any X/Y-plane of a given input image stack. This operation is similar to ImageJs 'Radial Reslice' method but offers less flexibility.

**Parameters**: Image source, Image destination, Number numberOfAngles, Number angleStepSize

**Available for**: 3D

**Macro example**:

```
run("CLIJ Macro Extensions", "cl_device=");
Ext.CLIJ_push(source);
Ext.CLIJ_resliceRadial(source, destination, numberOfAngles, angleStepSize);
Ext.CLIJ_pull(destination);
```

### CLIJ_resliceRight

Flippes X, Y and Z axis of an image stack. This operation is similar to ImageJs 'Reslice [/]' method but offers less flexibility such as interpolation.

**Parameters**: Image source, Image destination

**Available for**: 3D

**Macro example**:

```
run("CLIJ Macro Extensions", "cl_device=");
Ext.CLIJ_push(source);
Ext.CLIJ_resliceRight(source, destination);
Ext.CLIJ_pull(destination);
```

### CLIJ_resliceTop

Flippes Y and Z axis of an image stack. This operation is similar to ImageJs 'Reslice [/]' method but offers less flexibility such as interpolation.

**Parameters**: Image source, Image destination

**Available for**: 3D

**Macro example**:

```
run("CLIJ Macro Extensions", "cl_device=");
Ext.CLIJ_push(source);
Ext.CLIJ_resliceTop(source, destination);
Ext.CLIJ_pull(destination);
```

### CLIJ_rotate2D

Rotates an image in plane. All angles are entered in degrees. If the image is not rotated around the center, it is rotated around the coordinate origin.

It is recommended to apply the rotation to an isotropic image.

**Parameters**: Image source, Image destination, Number angle, Boolean rotateAroundCenter

**Available for**: 2D

**Macro example**:

```
run("CLIJ Macro Extensions", "cl_device=");
Ext.CLIJ_push(source);
```

```
Ext.CLIJ_rotate2D(source, destination, angle, rotateAroundCenter);
Ext.CLIJ_pull(destination);
```

### CLIJ_rotate3D

Rotates an image stack in 3D. All angles are entered in degrees. If the image is not rotated around the center, it is rotated around the coordinate origin.

It is recommended to apply the rotation to an isotropic image stack.

**Parameters**: Image source, Image destination, Number angleX, Number angleY, Number angleZ, Boolean rotateAroundCenter

**Available for**: 3D

**Macro example**:

```
run("CLIJ Macro Extensions", "cl_device=");
Ext.CLIJ_push(source);
Ext.CLIJ_rotate3D(source, destination, angleX, angleY, angleZ, rotateAroundCenter);
Ext.CLIJ_pull(destination);
```

### CLIJ_rotateLeft

Rotates a given input image by 90 degrees counter-clockwise. For that, X and Y axis of an image stack are flipped. This operation is similar to ImageJs 'Reslice [/]' method but offers less flexibility such as interpolation.

**Parameters**: Image source, Image destination

**Available for**: 2D, 3D

**Macro example**:

```
run("CLIJ Macro Extensions", "cl_device=");
Ext.CLIJ_push(source);
Ext.CLIJ_rotateLeft(source, destination);
Ext.CLIJ_pull(destination);
```

### CLIJ_rotateRight

Rotates a given input image by 90 degrees clockwise. For that, X and Y axis of an image stack are flipped. This operation is similar to ImageJs 'Reslice [/]' method but offers less flexibility such as interpolation.

**Parameters**: Image source, Image destination

**Available for**: 2D, 3D

**Macro example**:

```
run("CLIJ Macro Extensions", "cl_device=");
Ext.CLIJ_push(source);
Ext.CLIJ_rotateRight(source, destination);
Ext.CLIJ_pull(destination);
```

### CLIJ_scale

DEPRECATED: CLIJ scale() is deprecated. Use scale2D or scale3D instead!

**Parameters**: Image source, Image destination, Number scaling_factor, Boolean scale_to_center

**Available for**: 3D

**Macro example**:

```
run("CLIJ Macro Extensions", "cl_device=");
Ext.CLIJ_push(source);
Ext.CLIJ_scale(source, destination, scaling_factor, scale_to_center);
Ext.CLIJ_pull(destination);
```

### CLIJ_scale2D

Scales an image with a given factor.

**Parameters**: Image source, Image destination, Number scaling_factor, Boolean scale_to_center

**Available for**: 2D

**Macro example**:

```
run("CLIJ Macro Extensions", "cl_device=");
Ext.CLIJ_push(source);
Ext.CLIJ_scale2D(source, destination, scaling_factor, scale_to_center);
Ext.CLIJ_pull(destination);
```

### CLIJ_scale3D

Scales an image with a given factor.

**Parameters**: Image source, Image destination, Number scaling_factor, Boolean scale_to_center

**Available for**: 3D

**Macro example**:

```
run("CLIJ Macro Extensions", "cl_device=");
Ext.CLIJ_push(source);
Ext.CLIJ_scale3D(source, destination, scaling_factor, scale_to_center);
Ext.CLIJ_pull(destination);
```

### CLIJ_set

Sets all pixel values x of a given image X to a constant value v.

```
f(x) = v
```

**Parameters**: Image source, Number value

**Available for**: 2D, 3D

**Macro example**:

```
run("CLIJ Macro Extensions", "cl_device=");
Ext.CLIJ_push(source);
Ext.CLIJ_set(source, value);
```

### CLIJ_subtractImages

Subtracts one image X from another image Y pixel wise.

```
f(x, y) = x - y
```

**Parameters**: Image subtrahend, Image minuend, Image destination

**Available for**: 2D, 3D

**Macro example**:

```
run("CLIJ Macro Extensions", "cl_device=");
Ext.CLIJ_push(subtrahend);
Ext.CLIJ_push(minuend);
Ext.CLIJ_subtractImages(subtrahend, minuend, destination);
Ext.CLIJ_pull(destination);
```

### CLIJ_sumOfAllPixels

Determines the sum of all pixels in a given image. It will be stored in a new row of ImageJs Results table in the column 'Sum'.

**Parameters**: Image source

**Available for**: 2D, 3D

**Macro example**:

```
run("CLIJ Macro Extensions", "cl_device=");
Ext.CLIJ_push(source);
Ext.CLIJ_sumOfAllPixels(source);
```

### CLIJ_sumZProjection

Determines the sum projection of an image along Z.

**Parameters**: Image source, Image destination_sum

**Available for**: 3D

**Macro example**:

```
run("CLIJ Macro Extensions", "cl_device=");
Ext.CLIJ_push(source);
Ext.CLIJ_sumZProjection(source, destination_sum);
Ext.CLIJ_pull(destination_sum);
```

### CLIJ_threshold

Computes a binary image with pixel values 0 and 1. All pixel values x of a given input image with value larger or equal to a given threshold t will be set to 1.

f(x,t) = (1 if (x >= t); (0 otherwise))

This plugin is comparable to setting a raw threshold in ImageJ and using the 'Convert to Mask' menu.

**Parameters**: Image source, Image destination, Number threshold

**Available for**: 2D, 3D

**Macro example**:

```
run("CLIJ Macro Extensions", "cl_device=");
Ext.CLIJ_push(source);
Ext.CLIJ_threshold(source, destination, threshold);
Ext.CLIJ_pull(destination);
```

### CLIJ_translate2D

Translate an image stack in X and Y.

**Parameters**: Image source, Image destination, Number translateX, Number translateY

**Available for**: 2D

**Macro example**:

```
run("CLIJ Macro Extensions", "cl_device=");
Ext.CLIJ_push(source);
Ext.CLIJ_translate2D(source, destination, translateX, translateY);
Ext.CLIJ_pull(destination);
```

### CLIJ_translate3D

Translate an image stack in X, Y and Z.

**Parameters**: Image source, Image destination, Number translateX, Number translateY, Number translateZ

**Available for**: 3D

**Macro example**:

```
run("CLIJ Macro Extensions", "cl_device=");
Ext.CLIJ_push(source);
Ext.CLIJ_translate3D(source, destination, translateX, translateY, translateZ);
Ext.CLIJ_pull(destination);
```

121 plugins documented.
