## Supplementary Listing 2: Benchmarked operations for "CLIJ: GPU-accelerated image processing for everyone"

### Supplementary Listing 2: List of benchmarked operations

This list shows benchmarked operations and short Java code visualising how the operations were called. The full implementation of individual operation calls can be found in the respective Java classes:https://github.com/clij/clij-benchmarking-jmh/tree/master/src/main/java/net/haesleinhuepf/clij/benchmark/jmh

| **Operation** | **ImageJ** | **CLIJ** |
|---|---|---|
| AddImagesWeighted2D | `ImageCalculator ic = new ImageCalculator();`<br>`output = ic.run("Add create", input1, input2);` | `clij.op().addImagesWeighted(input1, input2,`<br>` output, 1f, 1f);` |
| AddImagesWeighted3D | `ImageCalculator ic = new ImageCalculator();`<br>`output = ic.run("Add create stack", input1,`<br>` input2);` | `clij.op().addImagesWeighted(input1, input2,`<br>` output, 1f, 1f);` |
| AddScalar2D | `IJ.run(input, "Add...", "value=1");` | `clij.op().addImageAndScalar(input, output,`<br>` 1f);` |
| AddScalar3D | `IJ.run(input, "Add...", "value=1 stack");` | `clij.op().addImageAndScalar(input, output,`<br>` 1f);` |
| AutoThreshold2D | `IJ.setAutoThreshold(input, "Default dark");`<br>`IJ.run(input, "Convert to Mask",`<br>` "method=Default background=Dark black");` | `clij.op().automaticThreshold(input, output,`<br>` "Default");` |
| AutoThreshold3D | `IJ.setAutoThreshold(input, "Default dark");`<br>`IJ.run(input, "Convert to Mask",`<br>` "method=Default background=Dark black");` | `clij.op().automaticThreshold(input, output,`<br>` "Default");` |
| BinaryAnd2D | `ImageCalculator ic = new ImageCalculator();`<br>`output = ic.run("AND create", input1, input2);` | `CLIJ` |
| BinaryAnd3D | `ImageCalculator ic = new ImageCalculator();`<br>`output = ic.run("AND create stack", input1,`<br>` input2);` | `clij.op().binaryAnd(input1, input2, output;` |
| Operation | `ImageJ` | `clij.op().binaryAnd(input1, input2, output;` |
| Erode2D | `IJ.run(input, "Erode", "");` | `clij.op().erodeSphere(input, output);` |

| | | |
|---|---|---|
| Erode3D | `import process3d.Erode_;`<br>`output = new Erode_().erode(input, 1, true);` | `clij.op().erodeSphere(input, output);` |
| FixedThreshold2D | `IJ.setThreshold(input, 128, 255);`<br>`IJ.run(input, "Convert to Mask",`<br>`"method=Default background=Dark black");` | `clij.op().threshold(input, output, 128f);` |
| FixedThreshold3D | `IJ.setThreshold(input, 128, 255);`<br>`IJ.run(input, "Convert to Mask",`<br>`"method=Default background=Dark black");` | `clij.op().threshold(input, output, 128f);` |
| Flip2D | `IJ.run(input, "Flip Horizontally", "");` | `clij.op().flip(input, output, true, false);` |
| Flip3D | `IJ.run(input, "Flip Horizontally", "stack");` | `clij.op().flip(clb3Da, clb3Dc, true, false,`<br>`false);` |
| GaussianBlur2D | `IJ.run(input, "Gaussian Blur...", "sigma=" +`<br>`sigma);` | `clij.op().blur(input, output, sigma, sigma);` |
| GaussianBlur3D | `IJ.run(input, "Gaussian Blur 3D...", "x=" +`<br>`sigma + " y=" + sigma + " z=" + sigma);` | `clij.op().blur(input, output, sigma, sigma,`<br>`sigma);` |
| MaximumZProjection | `IJ.run(input, "Z Project...",`<br>`"projection=[Max Intensity]");` | `clij.op().maximumZProjection(input, output);` |
| Mean2D | `IJ.run(input, "Mean...", "radius=" + radius);` | `kernelSize = CLIJUtilities.radiusToKernelSize(`<br>`radius);`<br>`clij.op().meanSphere(input, output, kernelSize,`<br>`kernelSize);` |
| Mean3D | `IJ.run(imp3D, "Mean 3D...", "x=" + radius +`<br>`" y=" + radius + " z=" + radius);` | `kernelSize = CLIJUtilities.radiusToKernelSize(`<br>`radius);`<br>`clij.op().meanSphere(input, output, kernelSize,`<br>`kernelSize, kernelSize);` |
| Median2D | `IJ.run(input, "Median...", "radius=" + radius);` | `kernelSize = CLIJUtilities.radiusToKernelSize(`<br>`radius);`<br>`clij.op().medianSphere(input, output,`<br>`kernelSize, kernelSize);` |
| Median3D | `IJ.run(imp3D, "Median 3D...", "x=" + radius +`<br>`" y=" + radius + " z=" + radius);` | `kernelSize = CLIJUtilities.radiusToKernelSize(`<br>`radius);`<br>`clij.op().medianSphere(clb3Da, clb3Dc,`<br>`kernelSize, kernelSize, kernelSize);` |
| Minimum2D | `IJ.run(imp, "Minimum...", "radius=" + radius);` | `kernelSize = CLIJUtilities.radiusToKernelSize(`<br>`radius);` |

| | | |
|---|---|---|
| | | `clij.op().minimumSphere(input, output,`<br>`kernelSize, kernelSize);` |
| Minimum3D | `IJ.run(input, "Minimum 3D...", "x=" + radius +`<br>`" y=" + radius + " z=" + radius);` | `kernelSize = CLIJUtilities.radiusToKernelSize(`<br>`radius);`<br>`clij.op().minimumSphere(input, output,`<br>`kernelSize, kernelSize, kernelSize);` |
| MultiplyScalar2D | `IJ.run(input, "Multiply...", "value=2");` | `clij.op().multiplyImageAndScalar(input,`<br>`output, 2f);` |
| MultiplyScalar3D | `IJ.run(input, "Multiply...", "value=2 stack");` | `clij.op().multiplyImageAndScalar(input, output,`<br>`2f);` |
| RadialReslice | `input.setRoi(new Line(input.getWidth() / 2,`<br>`input.getWidth() / 2, 0, 0));`<br>`Radial_Reslice_Copy rr =`<br>`new Radial_Reslice_Copy();`<br>`rr.setup("", input);`<br>`rr.run(input.getProcessor());` | `numberOfAngles = 360;`<br>`angleStepSize = 1.0f;`<br>`effectiveNumberOfAngles = (int)((float)`<br>`numberOfAngles / angleStepSize);`<br>`maximumRadius = (int)Math.sqrt(`<br>`Math.pow(input.getWidth() / 2, 2) +`<br>`Math.pow(input.getHeight() / 2, 2));`<br>`ClearCLBuffer output = clij.createCLBuffer(`<br>`new long[]{maximumRadius,`<br>`input.getDepth(),`<br>`effectiveNumberOfAngles},`<br>`input.getNativeType());`<br>`clij.op().radialProjection(input, output,`<br>`angleStepSize);` |
| Rotate2D | `IJ.run(imp2D, "Rotate 90 Degrees Left", "");` | `clij.op().rotateLeft(input, output);` |
| Rotate3D | `IJ.run(imp2D, "Rotate 90 Degrees Left", "");` | `clij.op().rotateLeft(input, output);` |
