## Supplementary Listing 3: Frequently Asked Questions for "CLIJ: GPU-accelerated image processing for everyone"

#### Which GPUs are supported by CLIJ?

CLIJ was successfully tested on a variety of Intel, Nvidia and AMD GPUs. See the full list of tested systems: https://clij.github.io/clij-docs/testedsystems

#### Do I have to buy a dedicated GPU in order to benefit from GPU-acceleration using CLIJ?

No. Common Intel Core and AMD Ryzen processors contain built-in GPUs which are compatible with CLIJ. However, as dedicated graphics cards come with their own GDDR-memory, additional speed-up can be gained by utilizing dedicated GPUs though.

#### With which operating systems is CLIJ compatible?

CLIJ was succesfully tested on Windows, MacOS, Fedora linux and Ubuntu linux. Current GPU and OpenCL drivers must be installed.

#### How can I achieve optimal performance using CLIJ?

In order to exploit GPU-accelerated image processing, one should

- Run as many operations as possible in a block without back and forth pulling/pushing image data to/from GPU memory.
- Process images larger than 10 MB (rule of thumb, depends on actual CPU/GPU hardware). Background: Image processing on the CPU can be pretty fast if the accessed memory is smaller than the cache of the CPU. When processing exceeds this cache size, using GPU might become beneficial.
- Process many images of the same size and type subsequently, because in that way compiled GPU-code can be reused.
- Reuse memory. Releasing and allocating memory takes time. Try to reuse memory if possible.

- Use a dedicated graphics card. When deciding for the right GPU, check the memory bandwidth. Image processing is usually memory-bound. The faster the memory access, the faster images can be processed. The computing power / clock rate of the GPU and number of compute cores is of secondary interest.
- Some CLIJ marked with "Box" in their name filters are implemented separable (Gaussian blur, minimum, maximum, mean filters). Separable filters are faster than others (e.g. marked with "Sphere").
- Further speedup can be achieved by combining filters on OpenCL kernel level. This means implementing OpenCL kernels containing whole workflows. This custom OpenCL code can be distributed as custom CLIJ plugin. A plugin template can be found here: https://github.com/clij/clij-plugin-template/

### How can I measure the speedup of my workflows?

The simplest way for measuring the speedup of workflows is using time measurements before and after execution, e.g. in ImageJ macro:

```
time = getTime(); // gives current time in milliseconds
// ...
// my workflow
// ...
print("Processing the workflow took " + (getTime() - time) + " msec"));
```

However, in order to make these measurements reliable, some hints shall be given:

- Measure the timing of execution in a loop several times. The first exection(s) may be slower than subsequent exeutions because of so called *warmup* effects.
- Exclude file input/output from the time measurements to exclude harddrive read/write speed from the performance benchmarking of your workflow.
- Also measure the similarity of the ImageJ and CLIJ workflows results. For example: Some CLIJ_*Box filters are potentially much faster than CLIJ_*Sphere filters, which are more similar to ImageJs filters. In this case, performance can be gained by paying with reduced workflow result similarity.

ImageJ macros benchmarking CPU/GPU performance can be found here:

- https://github.com/clij/clij-docs/blob/master/src/main/macro/benchmarking.ijm
- https://github.com/clij/clij-benchmarking/tree/master/src/main/macro_benchmarking_workflow

For more professional benchmarking, we recommend the OpenJDK Java Microbenchmark Harness (JMH). As the name suggests, this involves Java programming. You find more details here:https://openjdk.java.net/projects/code-tools/jmh/

To give an overview, some of CLIJs operations have been benchmarked with JMH: https://github.com/clij/clij-benchmarking-jmh

### Is CLIJ compatible with ImageJ without Fiji?

With some limitations, yes. You find details and installation instructions here: https://github.com/clij/clij-legacy/

### Does reusing memory bring additional speed-up?

Yes. When processing images of the same size and type, it is recommended to reuse memory instead of releasing memory and reallocating memory in every iteration. An example macro demonstrating this can be found here: https://github.com/clij/clij-docs/blob/master/src/main/macro/memory_reuse_versus_reallocation.ijm

### Are results of CLIJ filters expected to be exactly the same as when using ImageJ?

No. While algorithms on the CPU can make use of double-precision, common GPUs only support single precision for floating point numbers. Furthermore, following priorities were set while developing CLIJs filters:

- Mathematical correctness
- Consistency, e.g. results in 2D and 3D should be reasonably similar
- Simplicity of code to ease maintenance
- Performance
- Similarity of results generated with ImageJ

For example, the minimum filter of ImageJ takes different neighborhoods into account when being applied in 2D and 3D. CLIJs filters are consistend in 2D and 3D. Thus,

results may differ between ImageJ and CLIJ as shown here: Comparing CLIJs mean filter (center) and ImageJs mean filter (right) in 2D (top) and 3D (bottom). The result can be reproduced by running the following macro with radius = 1: https://github.com/clij/clij-docs/blob/master/src/main/macro/mean_detailed_comparison_IJ_CLIJ.ijm

### Which pixel values does CLIJ take into account when processing edge pixels of the image?

CLIJ in general uses the strategy `clamp to edge` assuming pixels outside the image have the same pixel value as the closest border pixel of the image. For transforms such as rotation, translation, scaling, and affine transforms, 'zero-padding' is applied assuming pixels having value 0 out of the image.

### Does CLIJ take physical pixel units into account?

No. All numeric spatial parameters in CLIJ such as radius and sigma are always entered in pixels. There is no operation in CLIJ which makes use of any physical units.

### Are pixel positions 0- or 1-indiced?

Pixel coordinates in X, Y and Z are zero-based indiced.

### Are multi-channel images and timelapse data supported by CLIJ?

In general no. CLIJ supports two and three dimensional images. If the third dimension represents channels or frames, these images can be processed using CLIJs 3D filters. When processing 4D or 5D images, it is recommended to split them into 3D blocks.

### Are in-place operations supported?

No. There are no in-place operations implemented in CLIJ. No built-in operation overwrites its input images. However, when implementing your own custom OpenCL-code and wrapping it into CLIJ plugins, in-place operations may be supported depending on used hardware, driver version and supported OpenCL version.

### Does it matter which is the currently active image window in ImageJ?

No. The currently active image window in ImageJ plays no role in CLIJ. Input and output images must be specified in macros by name explicitly.

### What happens in ImageJ macro if a specified output image doesn't exist?

If a specified output image does not exist in GPU memory, it will be generated automatically with a size defined by the executed operation with respect to input image and given parameters.

### What happens in ImageJ macro if a specified output image exists already?

If a specified output image exists already in GPU memory, it will be overwritten. If the output image has the wrong size, it will not be changed.

### What is the return value of Ext.CLIJ_... methods in ImageJ macro?

CLIJ operations called from ImageJ macro have no return values. They either process pixels and save results to images or they save their results to ImageJs results table.

### How are binary image characterized in CLIJ?

Binary output images are filled with pixel values 0 and 1. Any input image can serve as binary image and will be interpreted by differentiating 0 and non-zero values. In order to pull a binary image back to ImageJ which is compatible, use `pullBinary()`. This delivers a binary 8-bit image with 0 and 255 as pixel values.

### Are there performance benefits expected when calling OpenCL kernels directly via ClearCL instead of CLIJ?

Yes. CLIJ brings OpenCL-kernel caching and the possibility of image/pixel-type-independent OpenCL. These benefits come with small performance loss. Calling an OpenCL kernel via ClearCL directly may be about *a millisecond* faster than calling it via CLIJ. Example code demonstrating this is available here: [https://github.com/clij/clij-benchmarking/blob/master/src/main/java/net/haesleinhuepf/clij/benchmark/clearclclijcomparison/ClearCLVersusCLIJComparison.java](https://github.com/clij/clij-benchmarking/blob/master/src/main/java/net/haesleinhuepf/clij/benchmark/clearclclijcomparison/ClearCLVersusCLIJComparison.java)

### The CLIJ Java API offers methods for processing ClearCLBuffers and ClearCLImages. What's the difference?

Images and buffers are defined in the OpenCL standard. We tried to have as many operations as possible compatible to both, images and buffers. Differences are:

- When applying affine transforms and warping to images, linear interpolation is used. When using buffers, the nearest neighbor pixel delivers the resulting intensity.
- Images are not generally supported by GPU devices runnning OpenCL 1.1.
- For filters which access the local neighborhood of pixels, using images brings performance gain.

We recommend using buffers in general for maximum device compatibility.
