## Supplementary Listing 4: Used raw data and resulting figures for "CLIJ: GPU-accelerated image processing for everyone"

### Supplementary Listing 4: Raw data, processing scripts and resulting Figures

| [Raw] data item | Processing program / script | Figure / resulting data |
|---|---|---|
| #1 Images data used for benchmarking operations. Available online: https://git.mpi-cbg.de/rhaase/clij-benchmarking-data/tree/master/data/operation_benchmark_input | clij-benchmarking-JMH Java code and batch scripts available online: https://github.com/clij/clij-benchmarking-jmh | Benchmarking time measurements raw data:<br><br>#2 IJ and CLIJ Processing time versus image size<br>#3 IJ and CLIJ Processing time versus radii / sigma |
| #2 IJ and CLIJ Processing time versus image size https://github.com/clij/clij-benchmarking/tree/master/data/benchmarking-jmh/imagesize | Python Jupyter notebook https://github.com/clij/clij-benchmarking/blob/master/plotting_jmh/python/plotting_ij_clij_imagesize_comparison.ipynb | Figure 1b, Supplemental Figures 1 and 2 |
| #2 IJ and CLIJ Processing time versus image size https://github.com/clij/clij-benchmarking/tree/master/data/benchmarking-jmh/imagesize | Python Jupyter notebook https://github.com/clij/clij-benchmarking/blob/master/plotting_jmh/python/speedup_table.ipynb | Figure 1c |
| #3 IJ and CLIJ Processing time versus radii / sigma https://github.com/clij/clij-benchmarking/tree/master/data/benchmarking-jmh/kernelsize | Python Jupyter notebook https://github.com/clij/clij-benchmarking/blob/master/plotting_jmh/python/plotting_ij_clij_radii_comparison.ipynb | Figure 1b<br>Supplemental Figure 3 |
| #4 Image data used for workflow benchmarking. Available online: https://bds.mpi-cbg.de/CLIJ_benchmarking_data/ | ImageJ / ImageJ+CLIJ workflows https://github.com/clij/clij-benchmarking/tree/master/src/main | #5 Workflow time measurements<br>#6 Workflow resulting images<br>#7 Workflow cell count measurements |
| #5 Workflow time measurements https://github.com/clij/clij-benchmarking/tree/master/data/benchmarking/cellcount/ | Python Jupyter notebook https://github.com/clij/clij-benchmarking/blob/master/plotting/python/AnalyseWorkflowBTimes.ipynb | Time/speedup measurements for workflows in main text |
| #6 Workflow resulting images https://git.mpi-cbg.de/rhaase/clij-benchmarking-data/tree/master/data/benchmark | ImageJ Macros https://github.com/clij/clij-benchmarking/blob/master/src/main/macro_visualising_workflow_results/ | Supplemental video 1<br>Supplemental figure 6 |
| #7 Workflow cell count measurements https://github.com/clij/clij-benchmarking/tree/master/data/benchmarking/cellcount/ | Python Jupyter notebook https://github.com/clij/clij-benchmarking/blob/master/plotting/python/Analyse_differences.ipynb | Supplemental video 1 |
